# The Evolutionary History of Metallothioneins in Horseshoe Crabs, Scorpions, Spiders and other Chelicerate Species

**DOI:** 10.1101/2025.01.04.631296

**Authors:** Òscar Palacios, Mercè Capdevila, Ricard Albalat

## Abstract

Metallothioneins (MTs) are a heterogenous family of proteins involved in metal homeostasis and detoxification, playing a crucial role in the adaptation of organisms to varying conditions of metal bioavailability across ecosystems. Despite their significance, MT characterization across animal groups remains uneven. While MTs have been extensively studied in vertebrates and gastropods, their presence and function in many arthropods, particularly within the Chelicerata subphylum, remain largely unexplored. Chelicerates, comprising thousands of species of spiders, scorpions and ticks, are a highly diverse group of animals inhabiting a wide range of environments. Identifying and characterizing their MTs is therefore essential for understanding how they manage metal availability of diverse habitats. In this study, we have identified over 400 chelicerate MTs, classifying them into three distinct types –types 1, 2, and 3– each exhibiting significant structural and phylogenetic differences. Our findings have revealed that most chelicerate MTs are bidomain proteins, with notable exceptions of monodomain forms and larger variants. Evolutionary analysis suggests that type 1 MTs represent the ancestral form, while types 2 and 3 likely emerged during the evolution of spiders. These results pave the way of a deeper understanding of the metal homeostasis and detoxification in chelicerates, laying the groundwork for future studies on the origin, evolution and functional diversification of MTs in arthropods and other metazoan lineages.

## 1. Introduction

Heavy metals such as zinc (Zn) and copper (Cu) are essential for many biological processes, but harmful at high concentrations, while others such as cadmium (Cd) or lead (Pb) are toxic even at low concentrations (Tchounwou, et al. 2012). Having physiological mechanisms to control the homeostasis of the essential metals and to counteract the harmful effects of the toxic ones is, therefore, vital for all living beings. One of these mechanisms relies on a specialized type of metal-binding proteins called metallothioneins (MTs) that were first described in mammals by Margoshes and Vallee in 1957 (Margoshes and Vallee 1957). Classically, MTs have been considered small proteins (<100 amino acids) able to coordinate metal ions thanks to the metal-thiolate bonds of their abundant (up to 30%) cysteine (C) residues (Capdevila, et al. 2012; Ziller and Fraissinet-Tachet 2018). At a structural level, most MTs are modular proteins formed by one or two domains (MT domains) defined by the number and distribution of the CC, CxC or CCC motifs in which their cysteines are grouped (Braun, et al. 1986). This modular structure is thought to have been key in the evolvability of these proteins (Calatayud, et al. 2022) since it would have facilitated both the appearance of large MTs (>100 amino acids) resulting from tandem domain repeats, and the evolution of new MTs made with *de novo* emerged domains or with new rearrangements of the pre-existing ones (Calatayud, et al. 2018; Pedrini-Martha, et al. 2020; Calatayud, et al. 2021a; Calatayud, et al. 2021b; Calatayud, et al. 2021c; Garcia-Risco, et al. 2023; Dallinger, et al. 2024; Dallinger, et al. 2025).

In Metazoans, MTs have been characterized mostly in species of the Chordata and Mollusca phyla (Calatayud, et al. 2021b; Calatayud, et al. 2021c), whereas the number of MTs identified in other animal groups remains scarce and unevenly distributed (Blindauer and Leszczyszyn 2010; Blindauer 2014). For instance, in Arthropoda –the animal phylum with the largest number of species–, MTs have been identified only in some crustacean (malacostracans and brachiopods) (Brouwer, et al. 1995; Syring, et al. 2000; Valls, et al. 2001; Shaw, et al. 2007) and hexapod (insects) species (Luo, et al. 2020; and references therein), and remain unknown for most taxa, including two entire arthropod subphyla, that is, Chelicerata and Myriapoda.

The Chelicerata subphylum is the sister clade of all the other subphyla of Arthropoda and includes thousands of species of scorpions, ticks, mites, opiliones and spiders together with the “living fossil” horseshoe crabs (Giribet and Edgecombe 2012). The analysis of MTs from this animal clade is therefore not only key for understanding how this important group of animals deals with heavy metals, but also to achieve a comprehensive perspective of metal metabolism in arthropods, and the evolution and diversification of the MTs in the phylum.

With this purpose, in this work we aim to identify and characterize the chelicerate MTs, analyzing them in different species of the three classes in which this group is classified (**Figure 1**): class Pycnogonida, which includes the sea spiders; class Merostomata, which comprises the horseshoe crabs; and class Arachnida, the largest class within the subphylum, which includes spiders, scorpions, mites and ticks. Through exhaustive surveys of public databases, including raw data from transcriptomic and genomic high-throughput sequencing projects, we have been able to determine the sequence of potential MTs in numerous species distributed among the three classes of chelicerates, providing our analyses with a broad phylogenetic context.

**Figure 1.**
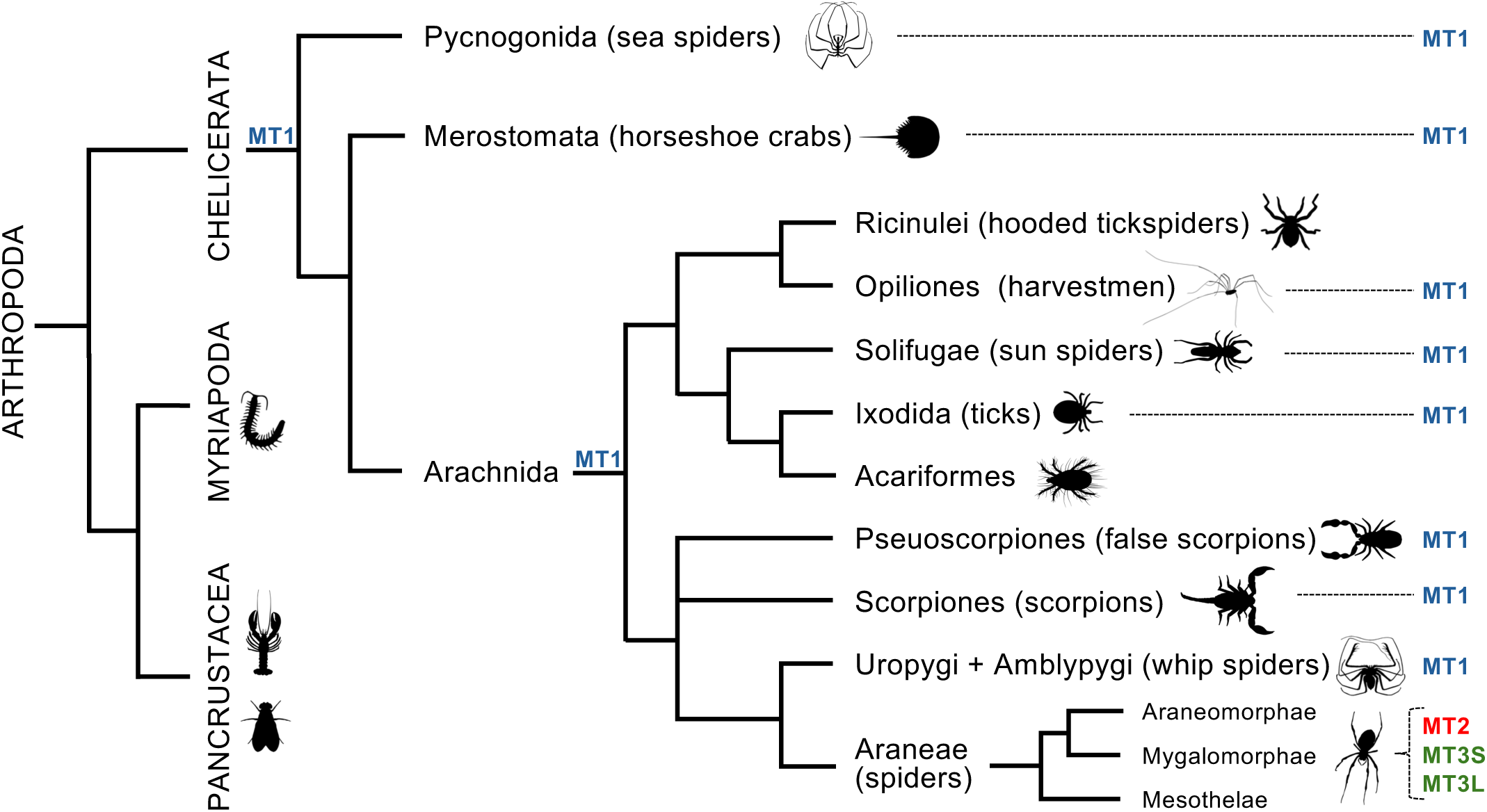
Metallothioneins in chelicerates. The Chelicerata subphylum, the sister group to the other arthropod clades (Myriapoda and Pancrustacea), is divided into three classes: Pycnogonida, Merostomata, and Arachnida. Type 1 MTs (MT1, in blue) are present across all chelicerate lineages except in the Araneae order, which contains type 2 (MT2, in red) and type 3 (MT3S and MT3L, in green) forms. This pattern suggests that MT1 is the ancestral form, lost in spiders, where MT2 and MT3 emerged as lineage-specific innovations. Animal silhouettes come from Phylopic (phylopic.org/).

We have identified over 400 potential MTs in chelicerates, revealing an unanticipated diversity due to the identification of three distinct MT types –types 1, 2 and 3– each exhibiting significant structural and phylogenetic differences. We have evaluated the metal-binding function of eight potential MTs, which were selected to represent the three MT types distributed among the three taxonomic classes. Chemical analyses of these MTs have confirmed their MT nature and have also revealed some differences in their metal selectivity and metal-binding capacity. Interestingly, structural analyses based on the cysteine motifs have exposed a modular organization for the chelicerate MTs, with a majority of bidomain proteins although with some notable exceptions of monodomain forms and large MT variants. Finally, evolutionary analyses have suggested that type 1 MTs would be the ancestral form, while type 2 and 3 MTs probably arose during the evolution of spiders. In summary, we have compiled the most extensive catalog of chelicerate MTs to date, paving the way for a deeper understanding of the metal homeostasis and detoxification in this important group of animals, while also laying the groundwork for future studies on the evolution and function of MTs across arthropods.

## 2. Results

### 2.1. Identification of Chelicerate MTs

Although the existence of chelicerate MTs had been inferred based on immunological and chemical assays (Wilczek 2005; Eraly, et al. 2010; Babczynska, et al. 2011a; Eraly, et al. 2011; Wilczek, et al. 2013; Aziz and Butt 2020; Lv, et al. 2021; Wisniewska, et al. 2022), the MTs of this group of animals had not been identified and characterized. Therefore, we began our analyses by searching public databases for potential MT sequences. We collected 429 putative MTs (**Table S1**) belonging to species distributed among the three Chelicerata classes: Pycnogonida, Merostomata and Arachnida (**Figure 1**). Based on the number and distribution of their cysteine motifs (CxC, CC, and CCC), which are used to predict the MT domains, we classified Chelicerata MTs in 3 types: MT of type1 (i.e. MT1), MT2 and MT3 (**Figure 2**). Each Chelicerata MT identified in this work was named according to the species and type to which it belonged (**Table S1**). The MT of the cross spider, for instance, was named AdiaMT2. “Adia” stood for the species name *<u>A</u>raneus <u>dia</u>dematus*, and “MT2” indicated that it was a type 2 form. In species with more than one isoform of the same type, each isoform was catalogued as “a”, “b”, “c” (e.g. isoforms of type 2 MT were named MT2a, MT2b, etc…). This naming convention helps to classify and distinguish the metallothioneins found across different chelicerate species based on their structural characteristics.

**Figure 2.**
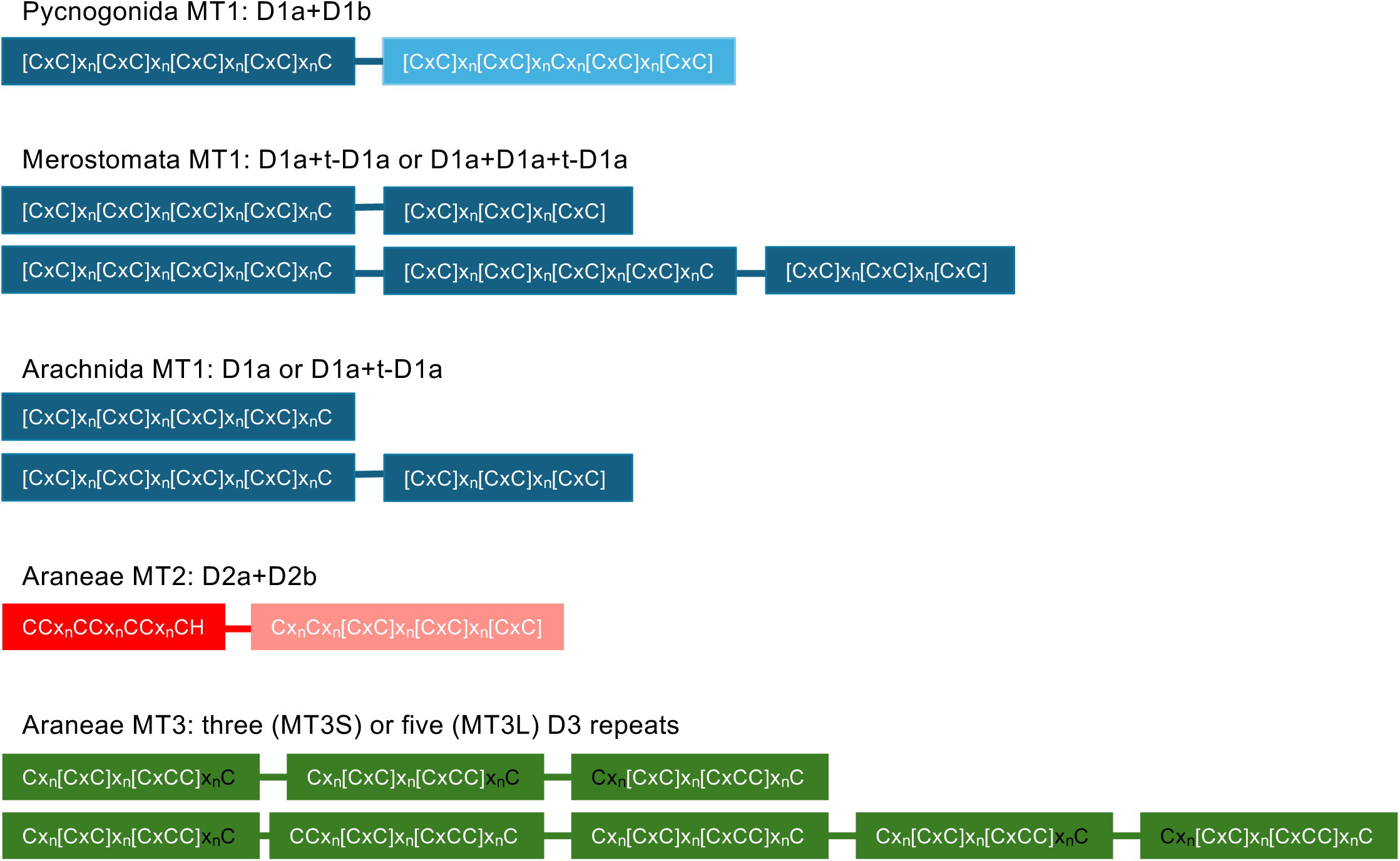
Domain organization of chelicerate metallothioneins. Based on the cysteine motif patterns, chelicerate MTs are classified into three types: MT1, MT2, and MT3. In Pycnogonida, MT1s consist of two domains, D1a (dark blue box) and D1b (light blue box). In Merostomata, MT1s typically comprise one or two complete D1a domains and a truncated D1a sequence (t-D1a) at the carboxyl-terminus. Arachnida MT1s exhibit a D1/t-D1 organization, similar to that of Merostomata, except in tick species and some harvestmen and sun spiders, which possess monodomain D1a forms. In Araneae, MT2s consist of two domains, D2a (red box) and D2b (pink box), while MT3s are composed of either three (MT3S) or five imperfect repeats (MT3L) of a 7-Cys D3 sequence (green box), with some repeats missing one cysteine (marked in black).

In the Pycnogonida class, we recognized seven putative MTs in seven species of sea spiders (**Figure S1A and Table S1**). We classified Pycnogonida MTs as type 1 and based on the pattern of their cysteine motifs we predicted that they were bidomain MTs made up of two distinct nine-cysteine domains, named D1a ([CxC]x_4_[CxC]x_3_[CxC]x_2_[CxC]x_2_C) and D1b ([CxC]x_5_[CxC]x_3_Cx_5_[CxC]x_3-2_[CxC]) (**Figure 2**). In the Merostomata class, we identified 10 putative MT1s in four species of horseshoe crabs (**Figure S1B and Table S1**). Merostomata MTs consisted of a variable number of domains, typically one or two complete D1a domains plus one truncated D1a sequence (t-D1a) at the carboxyl-end with only 6 cysteines (**Figure 2**). In the Arachnida class, our searches identified 57 putative MT1s in species of the orders Scorpiones (7 MT1s), Amblypygi (6 MT1s), Solifugae (4 MT1s), Opiliones (15 MT1s) and Ixodida (25 MT1s) (**Figure S1C and Table S1**). Most Arachnida MT1 exhibited a D1a/t-D1a organization, similar to Merostomata MT1s, with the exception of several monodomain D1a forms in all tick species (25 MT1s) and in some species of harvestmen (Opiliones) and sun spiders (Solifugae) (**Figure 2**).

Remarkably, MT1s were not found in any species of Araneae order (spiders), which is in fact the largest group of arachnids. Spiders, in contrast, exhibited MTs of type 2 and 3. We identified 114 putative MT2s from 110 spider species (**Figure S2 and Table S1**), which based on the pattern of their cysteine motifs would be bidomain MTs with an amino-terminal D2a domain with seven cysteines and one highly conserved histidine (i.e., CCx_5_CCx_4_CCx_6_CH), and a carboxyl-terminal D2b domain with eight cysteines (Cx_3_Cx_3_[CxC]x_5_[CxC]x_4_[CxC]) (**Figure 2**). These motifs were conserved in most of the 104 sequences of the Araneomorphae suborder but showed minor differences in the MTs of the Mesothelae (4 sequences: CCx_5_CCx_4_CCx_7_CC + CCx_8_[CxC]x_5_[CxC]x_4_[CxC]) and Mygalomorphae (6 sequences: CCx_5_CCx_4_Cx_6_CH + CCx_8_[CxC]x_5_[CxC]x_4_[CxC]) suborders (**Figure S2**). We also identified 241 putative MTs of type 3 across various spider species (**Figure S3 and Table S1**). These MT3s were composed of imperfect repeats of a D3 sequence (Cx_3_[CxC]x_2_[CxCC]x_4_C), typically conserving six out of seven cysteines. These MT3s appeared in two forms: short (S) and long (L). The short forms (151 MT3S in 118 species) consisted of three imperfect D3 repeats, while the long forms (90 MT3L in 87 species) generally contained five D3 repeats (**Figure 2 and Figure S3**), except for two MTs with six repeats in the Mygalomorphae class. The low number of cysteine residues in the D3 sequence –only 7– and the frequent loss of one cysteine in many repeats raise questions about whether D3 constitutes a structurally and functionally independent domain. While future experiments are needed to address this question, the identification of chelicerate MT3S and MT3L forms reinforces the notion that tandem sequence repetition is a recurrent and effective strategy in MT evolution for generating large variants with high metal-binding capacity (Dallinger, et al. 2025).

Interestingly, we found 59 species that contained all three Araneae MT forms: MT2, MT3S and MT3L (e.g. LcoeMT2, LcoeMT3Sa and b, and LcoeMT3L in the wolf spider *Lycosa coelestis*), suggesting a high degree of MT multiplicity across many spiders. Furthermore, the presence of MT2, MT3S and MT3L in species from the three Araneae suborders (Araneomorphae, Mesothelae and Mygalomorphae) (**Figure 1**) suggests that this multiplicity predated the diversification of spiders, over 350 million years ago (Lozano-Fernandez, et al. 2020).

### 2.2. Chemical Characterization of Chelicerata metal-MT complexes

#### 2.2.1. The MT nature of chelicerate proteins

To confirm the metallothionein nature of the putative chelicerate MTs, we chemical analyzed the metal-protein complexes of eight putative MTs representing the three Chelicerata classes –Pycnogonida, Merostomata, and Arachnida–. Five of these MTs belonged to type 1 MTs: NgraMT1 from the sea spider *Nymphon gracile*, representing the Pycnogonida MTs; LpolMT1a and LpolMT1b from the horseshoe crab *Limulus polyphemus*, as Merostomata MTs; and IricMT1a from the tick *Ixodes ricinus* and HpauMT1 from the scorpion *Hadogenes paucidens*, as Arachnida forms. The remaining three MTs were spider-specific MTs type 2 and type 3 from to the wolf spider *Lycosa coelestis*: LcoeMT2, LcoeMT3Sa and LcoeMT3L.

We produced the metal-MT complexes by heterologous expression of each MT in *Escherichia coli* cells grown in media supplemented with Zn, Cd, and Cu salts. The analysis of recombinant metal-protein complexes is a reliable method for determining their structural and functional characteristics, and it allows for assessing whether a cysteine-rich protein is a *bona fide* MT. This approach offers significant practical advantages over the isolation and purification of the complexes from native sources, while effectively minimizing methodological artifacts ((Cols, et al. 1997; Bofill, et al. 2001; Orihuela, et al. 2010; Palacios, et al. 2011; Dallinger, et al. 2024), for a critical review, see (Blindauer and Leszczyszyn 2010)). The complexes were purified and characterized using inductively coupled plasma atomic emission spectroscopy (ICP-AES) for protein quantification and metal-to-protein stoichiometry analysis (**Table 1**), and electrospray ionization mass spectrometry (ESI-MS) to determine the molecular mass of the formed metal-MT species, that is, the speciation of the samples (**Table 2**, **Figures 3 and 4** for analysis at neutral pH, observing total metal content, and **Figure S4** for data recorded at pH 2.4, observing the apo forms and Cu-bound proteins). The experimental masses of the apo-MTs after demetallation through acidification of the different productions (**Figure S4**) were consistently in agreement with the theoretical masses, confirming the identity of the recombinantly expressed proteins.

**Figure 3.**
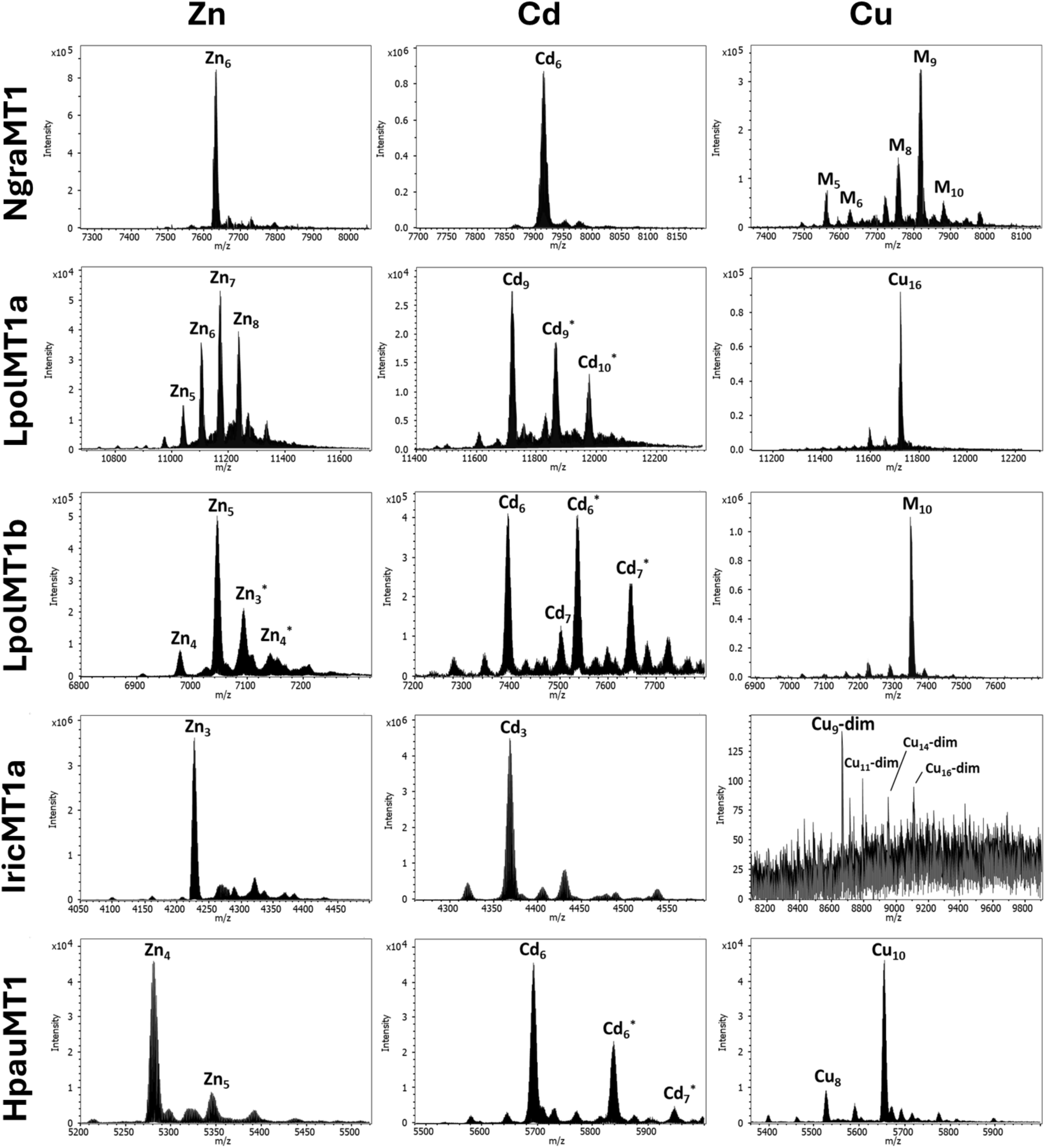
Deconvoluted ESI-MS spectra of the type 1 MTs (NrgaMT1, LpolMT1a, LpolMT1b, IricMT1a, and HpauMT1), recorded at pH 7.0, recombinantly produced in Zn-, Cd– and Cu-enriched media. M stands for Cu+Zn. * glycosylated species containing one hexamer unit.

**Figure 4.**
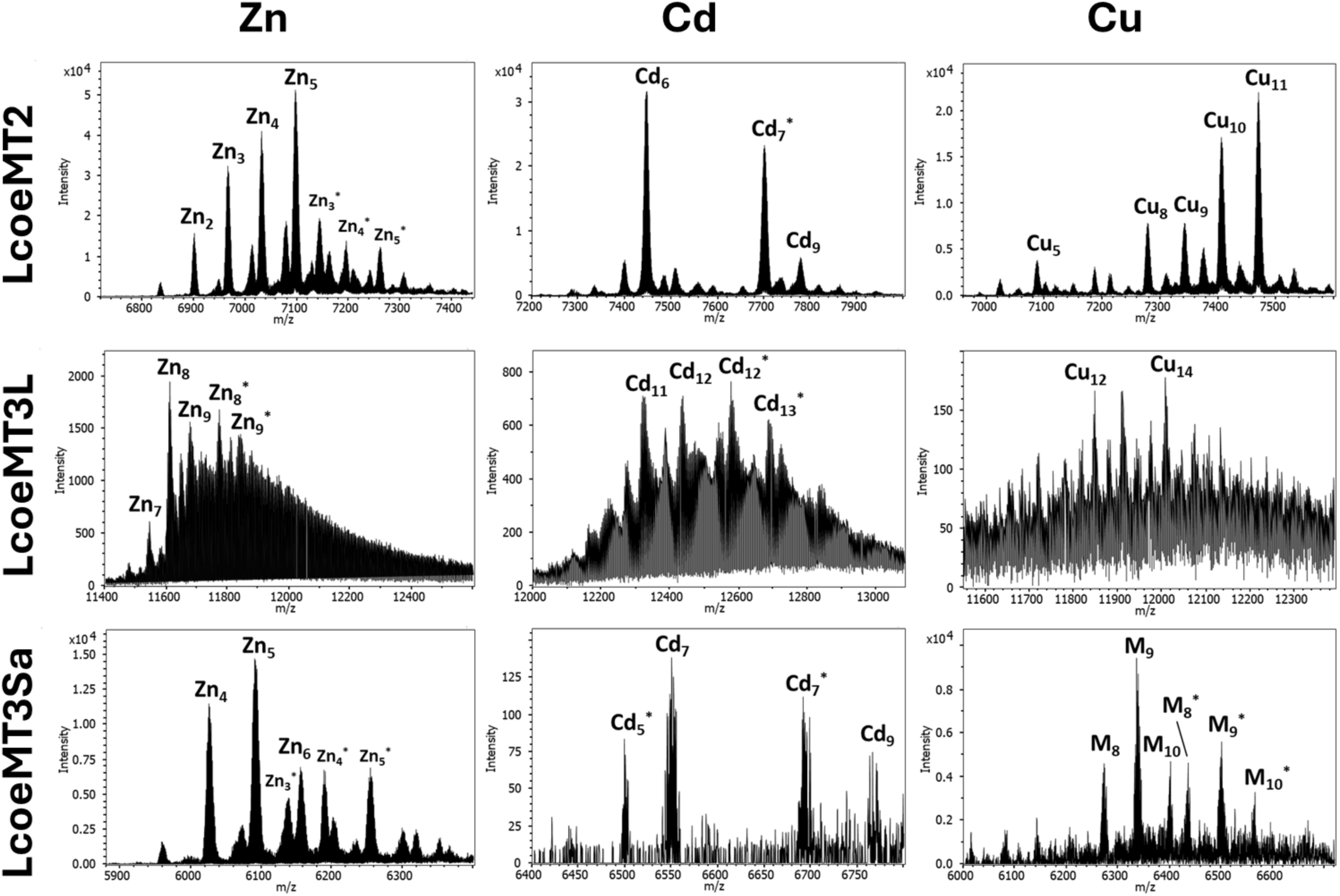
Deconvoluted ESI-MS spectra of the LcoeMT2, LcoeMT3L, and LcoeMT3Sa, recorded at pH 7.0, recombinantly produced in Zn-, Cd– and Cu-enriched media. M stands for Cu+Zn. * Glycosylated species containing 1 hexamer unit.

**Table 1.** ICP-AES results of the recombinant preparation of chelicerate MTs. Molar concentration of the protein measured by quantification of the S content and considering the number of Cys and Met of each protein, as well as metal/protein molar ratios measured considering the concentration detected for each metal.

| Protein | Metal supp. | [MT] x 10<sup>4</sup> (M) | Zn/MT | Cd/MT | Cu/MT |
| --- | --- | --- | --- | --- | --- |
| NgraMT1 | Zn | 1.92 | 5.2 | -- | -- |
|  | Cd | 1.62 | -- | 5.8 | -- |
|  | Cu | 0.79 | 3.7 | -- | 8.2 |
| LpolMT1a | Zn | 1.76 | 3.6 | -- | -- |
|  | Cd | 0.42 | -- | 8.6 | -- |
|  | Cu | 1.12 | -- | -- | 9.9 |
| LpolMT1b | Zn | 5.61 | 1.7 | -- | -- |
|  | Cd | 6.67 | -- | 1.4 | -- |
|  | Cu | 1.62 | 1.2 | -- | 8.6 |
| IricMT1 | Zn | 5.02 | 1.1 | -- | -- |
|  | Cd | 7.05 | -- | 1.3 | -- |
|  | Cu | 0.12 | -- | -- | 12.1 |
| HpauMT1 | Zn | 5.34 | 1.3 | -- | -- |
|  | Cd | 3.2 | -- | 3.2 | -- |
|  | Cu | 0.98 | -- | -- | 9.5 |
| LcoeMT2 | Zn | 2.73 | 1.7 | -- | -- |
|  | Cd | 4.39 | -- | 1.8 | -- |
|  | Cu | 1.21 | -- | -- | 9 |
| LcoeMT3L | Zn | 2.73 | 1.7 | -- | -- |
|  | Cd | 4.39 | -- | 1.8 | -- |
|  | Cu | 0.45 | -- | -- | 7.4 |
| LcoeMT3Sa | Zn | 0.16 | < 2.0 | -- | -- |
|  | Cd | 0.11 | -- | 7.1 | -- |
|  | Cu | 0.06 | -- | -- | 7.8 |

**Table 2.** Overview of the features and experimental data collected in the characterization of the chelicerate MT productions. The name of the proteins, number of amino acids (aa), cysteines (C) and histidines (H) are indicated in the three first columns. The type of domain organization, number of domains (#domains), and distribution of the coordinating amino acids among domains is summarized in the 4^th^ column. The presence of glycosylated species in the metal-MT preparations is indicated under the heading “Glycosylation”. The two next columns include the metal-MT species identified in the diGerent preparations as well as their corresponding Cys/M^2+^ ratios referred to the most important species. The last column assigns the Zn-, Cd-, or Cu-thionein character to each chelicerate MT.

| Name | aa | C/H | Domain type (# domains)<br># Cys/His | Glycosylation | Metal-species at neutral pH<br>(Cu-MT species at pH 2.4)<sup>a</sup> | Cys/M<sup>2+</sup> | Zn-/Cd-/Cu-thionein<br>character |
| --- | --- | --- | --- | --- | --- | --- | --- |
| NgraMT1 | 68 | 18 | D1a + D1b (2)<br>9 Cys + 9 Cys | Cu-species | Zn<sub>6</sub><br>Cd<sub>6</sub><br>M<sub>9</sub>>M<sub>8</sub>>M<sub>5</sub>... (Cu<sub>4</sub>>Cu<sub>8</sub>) | 3.0<br>3.0<br>-- | Zn-th |
| LpolMT1a | 100 | 24 | D1a + D1a + t-D1a (3)<br>9 Cys + 9 Cys + 6 Cys | Zn-species<br>Cd-species | Zn<sub>7</sub>>Zn<sub>8</sub>>Zn<sub>6</sub>...<br>Cd<sub>9</sub>>Cd<sub>9</sub>*>Cd<sub>10</sub>*<br>Cu<sub>16</sub> (*idem*) | 3.4<br>2.7<br>-- | Cu-th |
| LpolMT1b | 64 | 15 | D1a + t-D1a (2)<br>9 Cys + 6 Cys | Zn-species<br>Cd-species | Zn<sub>5</sub>>Zn<sub>3</sub>*...<br>Cd<sub>6</sub>≈Cd<sub>6</sub>*>Cd<sub>7</sub>*>Cd<sub>7</sub><br>M<sub>10</sub> (Cu<sub>10</sub>>>Cu<sub>8</sub>>Cu<sub>4</sub>) | 3.0<br>2.5<br>-- | Close to Cu-th |
| IricMT1a | 39 | 9 | D1a (1)<br>9 Cys | --- | Zn<sub>3</sub><br>Cd<sub>3</sub><br>Dimeric Cu-MT species (*idem*) | 3.0<br>3.0<br>-- | Cd-th |
| HpauMT1 | 51 | 15 | D1a + t-D1a (2)<br>9 Cys + 6 Cys | Zn-species<br>Cd-species<br>Cu-species | Zn<sub>4</sub>>Zn<sub>5</sub><br>Cd<sub>6</sub>>Cd<sub>6</sub>*>Cd<sub>7</sub>*<br>Cu<sub>10</sub>>>Cu<sub>8</sub> (*idem*) | 3.8<br>2.5<br>-- | Cu-th |
| LcoeMT2 | 65 | 15/1 | D2a + D2b (2)<br>7 Cys / 1 His + 8 Cys | Zn-species<br>Cd-species | Zn<sub>5</sub>>Zn<sub>4</sub>>Zn<sub>3</sub>...<br>Cd<sub>6</sub>>Cd<sub>7</sub>*>>Cd<sub>9</sub><br>Cu<sub>11</sub>>Cu<sub>10</sub>>Cu<sub>9</sub>... (Cu<sub>11</sub>>Cu<sub>8</sub>>Cu<sub>7</sub>...) | 3.0<br>2.5<br>-- | Close to Cd-th |
| LcoeMT3L | 112 | 33/1 | D3+D3+D3+D3+D3 (5) | Zn-species<br>Cd-species<br>Cu-species | Zn<sub>8</sub>>Zn<sub>8</sub>*>Zn<sub>9</sub>...<br>Cd<sub>11</sub>≈Cd<sub>12</sub>≈Cd<sub>12</sub>*...<br>Cu<sub>12</sub>≈Cu<sub>14</sub> (*idem*) | 3.7<br>3.0<br>-- | Multipurpose |
| LcoeMT3Sa | 57 | 18 | D3+D3+D3 (3) | Zn-species<br>Cd-species<br>Cu-species | Zn<sub>5</sub>>Zn<sub>4</sub>...<br>Cd<sub>7</sub>>Cd<sub>7</sub>*>Cd<sub>5</sub>*>Cd<sub>9</sub><br>M<sub>9</sub>>M<sub>9</sub>*... (Cu<sub>8</sub>>Cu<sub>8</sub>*>Cu<sub>9</sub>...) | 3.6<br>2.6<br>-- | Multipurpose |

<sup>a</sup>Asterisk (\*) indicates metal-MT glycosylated species. *idem* means that the speciation at acidic pH is the same than at neutral pH.

The metallothionein nature of putative chelicerate MTs was determined using the criterion of the proteins’ ability to form metal-protein clusters with Cys/M^2+^ ratios lower than 4:1, a feature that endows MTs with their high metal coordination capacity. These lower ratios result from the fact that in metallothioneins, metal clusters incorporate both terminal and bridging cysteine ligands, which organize the protein backbone around the metal ions. Consequently, the 4:1 Cys/M²⁺ molar ratio (four cysteines per divalent metal ion, M = Zn or Cd) expected if the divalent metal centers adopted a tetrahedral coordination with cysteine residues acting solely as terminal ligands, is reduced in MTs to 3:1 or less due to the involvement of some cysteine residues as bridging ligands. This reduction has been well documented for numerous MTs, including gastropod MTs forming two clusters with a 3:1 ratio (9 cysteines coordinating 3 Zn²⁺)(Perez-Rafael, et al. 2012; Dvorak, et al. 2018; Garcia-Risco, et al. 2021), and mammalian MTs with two clusters exhibiting 2.75:1 and 3:1 ratios (11 cysteines coordinating 4 Zn²⁺ and 9 cysteines coordinating 3 Zn²⁺, respectively)(Otvos and Armitage 1980; Braun, et al. 1986; Singh, et al. 2023). The eight chelicerate MTs analyzed satisfied the “metallothionein” criterion by rendering metal-protein preparations with Cys/M^2+^ ratios ranging from 2.5 to 3.0 for at least one of the two divalent metal ions (Zn^2+^ or Cd^2+^) (**Table 2**), thereby confirming their metallothionein nature.

#### 2.2.2. Metal binding capacity and selectivity

We also analyzed the binding capacity and metal selectivity of chelicerate MTs. The binding capacity, quantitatively described by the metal-to-protein stoichiometry, varied among chelicerate MTs, although the Cys/M²⁺ ratios consistently remained ≤ 3:1 for all of them. Most chelicerate MTs, such as NgraMT1, LpomMT1b, HpauMT1, and LcoMT2, were bidomain proteins with a metal-to-protein stoichiometry of 5:1 to 6:1, corresponding to 5–6 divalent ions per protein molecule containing 15–18 cysteines (i.e. 3:1 to 2.5:1 Cys/M²⁺ ratios; **Table 2** and **Figures 3 and 4**). In contrast, small monodomain forms, such as IricMT1a, exhibited a halved metal-to-protein stoichiometry of 3:1, coordinating 3 divalent ions per protein molecule containing 9 cysteines, whereas larger MTs like LcoeMT3L had a nearly doubled stoichiometry of 11:1 to 13:1, binding 11–13 ions per MT molecule with 33 cysteines and one histidine (**Table 2** and **Figures 3 and 4**).

Regarding metal selectivity, our results revealed that chelicerate MTs exhibit diverse metal preferences (**Table 2** and **Figures 3 and 4**). NgraMT1, IricMT1a, and LcoeMT2 showed a clear preference for divalent ions and were classified as Zn-or Cd-thioneins (Zn-th or Cd-th), while LpolMT1a, LpolMT1b, and HpauMT1 were categorized as Cu-th due to their preference for monovalent metal ions. In contrast, LcoeMT3Sa and LcoMT3L did not show a clear preference for either divalent or monovalent ions, classifying them as multipurpose proteins. This classification was based on metal-protein complexes formed when expressed recombinantly in Zn-, Cd-or Cu-enriched culture media. For example, NgraMT1 was classified as a Zn-th because it formed single, well-folded species with divalent ions (i.e. Zn_6_-NgraMT1), but produced mixtures of heteronuclear Zn,Cu-MT species in presence of Cu^+^ ions (i.e. Zn,Cu_5-10_-NgraMT1; **Figures 3, top row**). IricMT1a and LcoeMT2 were categorized as Cd-th because they also produced well-folded species with divalent ions (i.e. Cd_3_-IricMT1a and Cd_6_-LcoeMT2), but produced mixtures of homonuclear Cu-MT species in Cu-enriched productions (i.e. Cu_9-16_-IricMT1a and Cu_8-11_-LcoeMT2; **Figures 3, fourth row and Figure 4 top row**). In contrast, LpolMT1a, LpolMT1b, and HpauMT1 formed single, well-folded homonuclear Cu-MT species in Cu-enriched media (i.e. Cu_16_-LpolMT1a, Cu_10_-LpolMT1b, and Cu_10_-HpauMT1) but produced mixtures of species in Zn– and Cd-enriched cultures (**Figures 3, rows 2, 3 and 5**), thus classifying them as Cu-th. Finally, LcoeMT3Sa and LcoeMT3L were considered multipurpose MTs as they did not form single, well-folded species with any metal (**Figure 4 middle and bottom rows**).

Further support for this classification came from analyzing glycosylated species in recombinant metal-MT productions. In certain cases, MTs produced in *E. coli* in the presence of non-cognate metal ions are glycosylated with hexoses, leading to an increase in their molecular weight by increments of 162 Da compared to the expected apo-form (Garcia-Risco, et al. 2022). Conversely, glycosylation is negligible when MTs are produced with their cognate metals, as the resulting unique metalated species exhibits a highly structured conformation that sterically hindered enzymes and sugar nucleotides access to amino acid side chains (Garcia-Risco, et al. 2022). Therefore, our assignment of metal selectivity of chelicerate MTs was consistent with the negligible glycosylation levels observed in the apo-forms of NgraMT1 in the presence of Zn, IricMT1a and LcoeMT2 in the presence of Cd, and LpolMT1a, LpolMT1b, and HpauMT1 in the presence of Cu (**Figure S4**). In contrast, the multipurpose MTs, LcoeMT3a and LcoeMT3b, exhibited significant glycosylation of their apo-forms (**Figure S4**), likely due to their less tightly structured and more flexible conformation, indicating the absence of a strong metal-binding preference.

In summary, the chemical analyses of metal-protein complexes confirmed the metallothionein nature of the eight selected chelicerate proteins, validating our approach for searching, identifying, and characterizing new MTs across different animal groups. Our analyses further revealed the remarkable structural and functional diversity of chelicerate MTs, as they displayed different metal coordination capacities –ranging from 3 to 13 divalent metal ions per molecule– and distinct metal selectivities –from Zn-, Cd-, and Cu-thioneins to multipurpose forms– likely reflecting functional adaptations of the MTs to animals with different metal-specific needs.

## 3. Discussion

Chelicerates exhibit substantial resistance to heavy metals. For instance, horseshoe crabs display significantly higher tolerance to Cu and Zn compared to other marine arthropods (Botton, et al. 1998), and spiders thrive in metal-polluted habitats (Marczyk, et al. 1993; Wilczek and Migula 1996; Wilczek and Babczyńska 2000; Hendrickx, et al. 2003; Babczynska, et al. 2011a; Jung and Lee 2012). However, the underlying mechanisms of metal tolerance in chelicerates remain poorly understood. While immunological and chemical analyses suggest a potential role for chelicerate MTs in metal homeostasis and detoxification (Wilczek 2005; Eraly, et al. 2010; Babczynska, et al. 2011a; Babczynska, et al. 2011b; Eraly, et al. 2011; Wilczek, et al. 2013; Aziz and Butt 2020; Lv, et al. 2021; Wisniewska, et al. 2022), the lack of specific data on chelicerate MTs restricts this possibility to mere hypotheses. In this context, our work is particularly significant, as it provides the first identification and characterization of chelicerate MTs (**Figures 1 and 2; Figures S1, S2 and S3 and Table S1**), revealing a remarkable diversity in their metal coordination capacities and metal selectivities (**Table 2 and Figures 3 and 4**).

### 3.1. Metal-binging Features: MT Domains and Coordination Capacity

The coordination capacity of an MT correlates directly with its number of domains. MTs are usually composed of two domains. For instance, vertebrate MTs have a β-domain and an α-domain that coordinate three and four divalent metal ions, respectively (Otvos and Armitage 1980; Braun, et al. 1986). Similarly, most mollusk MTs are bidomain proteins that combine two of six possible domains – α, β1, β2, β3, ψ and 8– in lineage-specific arrangements to coordinate 6-7 metal ions (Calatayud, et al. 2021c; Calatayud, et al. 2022). Most chelicerate MTs have also a bidomain structure. For instance, based on their cysteine motifs, Pycnogonida MT1s would be composed of D1a and D1b domains, while Arachnida MT2s would consist of D2a and D2b domains (**Figure 2; Figures S1A and S2**). These bidomain MTs have a metal-binding capacity of around six divalent ions, i.e. three ions per domain. However, during evolution, monodomain MTs with reduced coordination capacity arose in certain animal lineages (Egli, et al. 2006; Atrian 2009; Calatayud, et al. 2021b), with chelicerates being one of them. Specifically, in all tick species and in some species of Opiliones and Solifugae, MT1s are monodomain proteins, composed of a single D1a domain with nine cysteines (**Figure 2**; **Figure S1C**). The chemical characterization of IricMT1a from the castor bean tick demonstrates the structural and functional independence of the D1a domain and confirms the reduced metal-to-protein stoichiometry of these monodomain MTs (**Figure 3**). The domain organization of other Arachnida MT1s is, however, more complex, although they always maintain the Cys/M²⁺ ratio of ≤ 3:1 in their stoichiometries. Thus, some MT1s contain a full D1a domain alongside a truncated version (t-D1a) with six or fewer cysteines, a configuration similar to that observed in Merostomata MT1s (**Figure 2; Figure S1B and C**). Since the D1a/D1b organization in Pycnogonida MT1s likely represents the ancestral structure (see below), our findings suggest that the D1b domain was lost in chelicerates after the divergence of the Pycnogonida lineage. Following the loss of D1b, the remaining D1a domain appears to have duplicated once or twice in Merostomata and Arachnida species. In these lineages, the carboxyl-terminal copy gradually degenerated into a truncated form (t-D1a) or disappeared entirely, resulting in the monodomain MTs observed in ticks and other arachnid species.

In contrast to the small monodomain MTs, other chelicerate MTs exhibit enhanced binding capacities due to a significant increase in size (>100 amino acids), resulting from multiple tandem repeats of their sequence. We have identified several large MTs in spiders, specifically the MT3L forms. For example, LcoeMT3L consists of 110 amino acids and can coordinate up to 11–13 divalent metal ions per molecule (**Figure 4**), which is almost twice the binding capacity of a standard bidomain form. Recent studies have also identified large MTs with tandem domain repeats in species of chordate tunicates and mollusk gastropods (Baumann, et al. 2017; Calatayud, et al. 2018; Pedrini-Martha, et al. 2020; Calatayud, et al. 2021b; Calatayud, et al. 2021c), and thereby, our findings extend the evolution of large MTs to the arthropod phylum. Overall, our results support the idea that small monodomain MTs and large multidomain MTs have evolved independently across various animal groups, from tunicates and insects to gastropods and arachnids (Egli, et al. 2006; Baumann, et al. 2017; Calatayud, et al. 2018; Pedrini-Martha, et al. 2020; Calatayud, et al. 2021a; Calatayud, et al. 2021b; Calatayud, et al. 2021c), likely favored by the flexibility and versatility provided by their modular architecture (Calatayud, et al. 2022; Dallinger, et al. 2025). However, the biological factors driving the reiterative evolution of MTs with reduced (monodomain) or enhanced (multidomain) metal-binding capacities across animal lineages remain unclear, and identifying them will pose a significant challenge for environmental toxicology and evolutionary biology in the coming years.

### 3.2. Metal-binging Features: Metal Selectivity

Regarding metal selectivity, we have found that chelicerate MTs exhibit distinct metal preferences, from Zn-, Cd-, and Cu-thioneins to multipurpose forms. Specifically, sea spider NgraMT1 behaves as a Zn-th, while tick IricMT1a is a clear Cd-th. Spider LcoeMT2 shows a non-extreme Cd-th character. Horseshoe crab LpolMT1a and scorpion HpauMT1 are genuine Cu-th while LpolMT1b prefers Cu over Zn and Cd but not in an extreme manner. Finally, the spider LcoeMT3Sa and LcoeMT3L should be considered multipurpose MTs, this is without a clear metal preference, which means that they can be used by the organisms to deal with a variety of metal ions, toxic and essential, in a set of physiological conditions. It has been proposed that when an MT has never been isolated and no relevant functional data is available, determining its “metal selectivity” may be the best option for determining its potential physiological function, that is, its “metal specificity”(Dallinger 2024). This makes our results particularly relevant, as information on metal physiology in chelicerates is very limited. For example, a set of proteins called metallothionein-like proteins (MTLPs) is known to be induced in metal-contaminated habitats (Babczynska, et al. 2011a), though these proteins have not yet been characterized. It has also been proposed that MTLPs may transport Zn and Pb to type A granules, and Cd and Cu to type B granules (Babczynska, et al. 2011a), where spiders store these metals (Jung and Lee 2012), but this transport has not yet been characterized. Our findings could allow to link these uncharacterized MTLPs to the specific MTs identified in this study and investigate which MTs mediate metal transport to A and B granules. Overall, our results pave the way for a deeper understanding of metal physiology in chelicerates and the functional innovations this group of animals has had to develop to thrive in a wide range of ecosystems.

### 3.3. ORIGIN OF CHELICERATE MTS AND *DE NOVO* EVOLUTION OF MTS IN SPIDERS

Piecing together the history of metallothioneins remains a significant challenge due to their high structural heterogeneity and possible independent evolutionary origins (Capdevila and Atrian 2011; Blindauer 2014; Isani and Carpene 2014; Ziller and Fraissinet-Tachet 2018; Calatayud, et al. 2021b; Calatayud, et al. 2021c). This heterogeneity and lack of homology invalidate evolutionary analyses based on conventional phylogenetic methods that assume a common ancestor among the aligned sequences, and call for alternative approaches. In this context, it has been suggested that alternative molecular markers, such as the patterns of cysteine motifs (CxC, CC, and CCC) in the MT domains, could be valuable for inferring homology relationships and conducting evolutionary studies (Calatayud, et al. 2021b). MTs with similar patterns are likely to share ancestry and be homologous, while those with distinct patterns would be analogous proteins with independent evolutionary origins.

Based on the patterns of cysteine motifs, we classified Chelicerata MTs into three types: MT1, MT2 and MT3 (**Figure 2**). The presence of MT1s across the three chelicerate classes –Pycnogonida, Merostomata and Arachnida (**Figure 1**)– suggest that type 1 represents the ancestral form. In contrast, type 2 and type 3 MTs, which are only found in species of the Araneae order, appear to be specific innovations of spiders. Surprisingly, the cysteine pattern in other reported arthropod MTs totally differs from that of chelicerate MTs. However, as mentioned above, arthropod MTs have been identified only in a limited number of species from the crustacean and hexapod subphyla, leaving many taxonomic groups unexamined. To address this gap, we have initiated a project to identify and characterize the MTs of myriapods, the fourth Arthropoda subphylum and the sister group of both crustaceans and hexapods. Preliminary results have revealed putative MT sequences in two myriapod species, *Lithobius forficatus* and *Glomeridella minima*, with a D1a/D1b organization similar to chelicerate MT1s found in the Pycnogonida class (**Figure S5**). The most parsimonious scenario to explain this phylogenetic distribution is that D1a/D1b MT1s are the ancestral forms in Arthropoda, and thus, they represent the ancestral chelicerate type. Supporting this possibility, comparison of the cysteine pattern of chelicerate and myriapod MT1s with that of other metazoan MTs reveals similarities with mollusk and chordate MTs (**Figure S5**).

Specifically, the patterns of D1a and D1b domains are identical to the patterns of β2 and β1 domains of mollusk MTs (Calatayud, et al. 2021c), and similar to β and α domains of vertebrate and cephalochordate forms (Calatayud, et al. 2021b). All these results support the idea that bidomain MT1 represents the ancestral MT of arthropods as they probably appeared before the diversification of bilaterians.

Having established the ancient origin of MT1s, further analysis is needed to understand the emergence of MT2 and MT3 forms. Given that these MTs seem restricted to spiders, the most plausible scenario is that they evolved between 450 and 350 MYA, after spiders diverged from Uropygi and Amblypygi species, but before the diversification of the Araneae order (Lozano-Fernandez, et al. 2020). We have previously proposed that *de novo* evolution of MT proteins or domains is a frequent event (Calatayud, et al. 2021b; Calatayud, et al. 2021c; Calatayud, et al. 2022), as the primary requirements for a peptide to function as a metal ion chelator only are the presence of a number of coordinating residues (such as cysteines) and a relatively short length that facilitates proper folding (Capdevila and Atrian 2011; Calatayud, et al. 2021c). The source of new MT sequences remains unknown, but the structural characteristics of the peptides encoded by short open reading frames (sORFs) in long non-coding RNAs (lncRNAs) concur well with those of MTs: small size, intrinsically disorder, and a poor conservation among lineages due to their diverse origins (Reinhardt, et al. 2013; Ruiz-Orera, et al. 2014; Ji, et al. 2015; Ruiz-Orera, et al. 2020). It can therefore be hypothesized that through the high evolutionary potential of sORFs to explore the functional value of new peptides, spiders would have generated novel MTs. These new MTs would have facilitated them the colonization of habitats with a wide range of metal availability –from deserts to rainforests, mountains to caves, and even urban areas– enabling spiders to become the most diverse group of chelicerates, capable of inhabiting nearly every terrestrial ecosystem on Earth.

To conclude, it is important to note that several important questions regarding the evolutionary history of MTs remain unresolved: What ecological or physiological factors account for the disparity in MT diversity and multiplicity among chelicerate species? What insights can the evolutionary history of chelicerate MTs provide about MT evolution in arthropods, in many of which MTs have not yet been identified? Is it feasible to reconstruct the evolutionary history of MTs in metazoans? Addressing these questions will require additional research that combine genetic, biochemical, and structural analyses of numerous MTs from a wide variety of organisms, with ecological, toxicological and physiological studies of the species that possess them.

## 4. Material and Methods

### 4.1. Database Searches and Sequence Analyses

Chelicerata MT sequences were identified by low-restrictive tblastn searches in Transcriptome Shotgun Assemble (TSA) and other NCBI databases with a high value of expect threshold and no filtering for low complexity sequences, using the arthropod MTs and a selection of other animal MTs as queries. Retrieved sequences were manually inspected and promising MT sequences were used as queries for additional tblastn or blastn searches in Sequence Read Archive (SRA) of RNA databases. Raw sequence data retrieved from the SRA searches were assembled using SeqMan Ultra 17.5.0 software from the DNASTAR Lasergene package, and manually inspected in order to reconstruct new MT sequences. Sequence alignments were generated by MUSCLE and reviewed manually with the package AliView v1.28 (Larsson 2014).

### 4.2. Production and Purification of Recombinant Metal-MT Complexes

Production and purification of recombinant metal-MT complexes were performed as described elsewhere (Calatayud, et al. 2018; Calatayud, et al. 2021b). In brief, synthetic cDNAs codifying the different constructs were provided by Synbio Technologies (Monmouth Junction, NJ, USA), cloned in the pGEX-4T-1 expression vector (GE Healthcare, Chicago, IL, USA) and transformed in protease-deficient *E. coli* BL21 strain. Metal-MT complexes were produced in *E. coli* BL21 cultures expressing the recombinant plasmids, after induction with isopropyl-β-D-thiogalactopyranoside (100 μM) and supplementation with ZnCl_2_ (300 μM), CdCl_2_ (300 μM) or CuSO_4_ (500 μM). Metal-MT complexes were purified from the soluble protein fraction of sonicated bacteria by affinity purification of the GST-tagged proteins, and digestion with thrombin. The metal-MT complexes were concentrated with a 3 kDa Centripep Low Concentrator (Amicon, Merck), and fractionated on a Superdex-75 FPLC column (GE Healthcare) equilibrated with 20 mM Tris-HCl, pH 7.0 and run at 0.8 mL min^−1^. The protein-containing fractions, identified by their absorbance at 254 nm, were pooled and stored at −80°C until use.

### 4.3. Analysis of Metal-MT Complexes

The metal-MT complexes were analyzed by ICP-AES for protein quantification and element composition with an Optima 4300DV (Perkin-Elmer, MA, USA) apparatus (S, 182.040 nm; Zn, 213.856 nm; Cd, 228.802 nm; and Cu, 324.803 nm) under conventional conditions following an already established method (Bongers, et al. 1988). Molecular weights were determined by ESI-MS, in a MicroTof-Q instrument (Bruker Daltonics Gmbh, Bremen, Germany) connected to a Series 1100 HPLC pump (Agilent Technologies) controlled by the Compass Software. The instrument was calibrated with ESI-L Low Concentration Turning Mix (Agilent Technologies, Santa Clara, CA). Metallated forms were detected under native conditions: 20 μL of sample injected through a PEEK tube at 30-50 μL min^-1^ in a 3.5–5.0-kV capillary-counter voltage, at 90-110 °C of desolvation temperature, and with dry gas at 6 L min^-1^. Spectra were recorded between a m/z range from 800 to 3,000. The liquid carrier was a 90:10 mixture of 15 mM ammonium acetate at pH 7.0 and acetonitrile. Apo-forms and Cu-species resistant at acidic pH were analyzed under similar ESI-MS conditions but using an 80:20 mixture of aqueous formic acid solution at pH 2.4 and acetonitrile. All molecular masses were calculated according to the bibliography (Fabris, et al. 1996). It is important to take into consideration that due to the close mass between Cu(I) and Zn(II) ions, in the analysis of protein they cannot be easily distinguished by MS, thus, in ESI-MS data recorded at neutral pH, when the presence of both ions was confirmed by ICP-AES, the heteronuclear species detected were denoted as M (M = Zn+Cu).

## 5. Supplementary Material

Supplementary material includes Table S1, and Figures S1, S2, S3, S4 and S5, and they are available online.

## 6. Acknowledgements

We acknowledge the Centres Científics i Tecnològics (CCiT) de la Universitat de Barcelona (DNA sequencing) and the Servei d’Anàlisi Química (SAQ) de la Universitat Autònoma de Barcelona (ICP-AES, ESI-MS) for allocating instrument time. We also thank to Sebastian Artime for experimental support.

## 7. Funding

R.A. was supported by Grant PID2021-123258NB-I00 funded by MICIU/AEI/ 10.13039/501100011033 and by ERDF/EU. M.C. and Ò.P. were supported by BIO2015-67358-C2-2-P and PID2022-138479NB-I00 from the Spanish Ministerio de Ciencia e Innovación. R.A. was member of the “Grup de Recerca de la Generalitat de Catalunya” ref. 2021 SGR 00372, and M.C. and Ò.P. of the ref. 2021SGR00668.

## 8. Author Contributions

Conceptualization: M.C., Ò.P. and R.A.; Methodology, Ò.P. and R.A.; Formal analysis, Ò.P., M.C. and R.A.; Investigation, Ò.P. and R.A.; Funding acquisition: M.C., Ò.P. and R.A.; Project administration, Ò.P. and R.A.; Writing (original draft) R.A.; Writing (review and editing) M.C., Ò.P. and R.A.; All authors have read and agreed to the published version of the manuscript.

## 9. Data Availability

The data underlying this article are available in NCBI database: https://www.ncbi.nlm.nih.gov/

## 10. Conflicts of interest

The authors declare no conflict of interest.

## Notes

### Competing Interest Statement

The authors have declared no competing interest.

