## Supplementary figures and images for "The Evolutionary History of Metallothioneins in Horseshoe Crabs, Scorpions, Spiders and other Chelicerate Species"

### Figure S1

#### Figure S1

##### A) Pycnogonida MT1

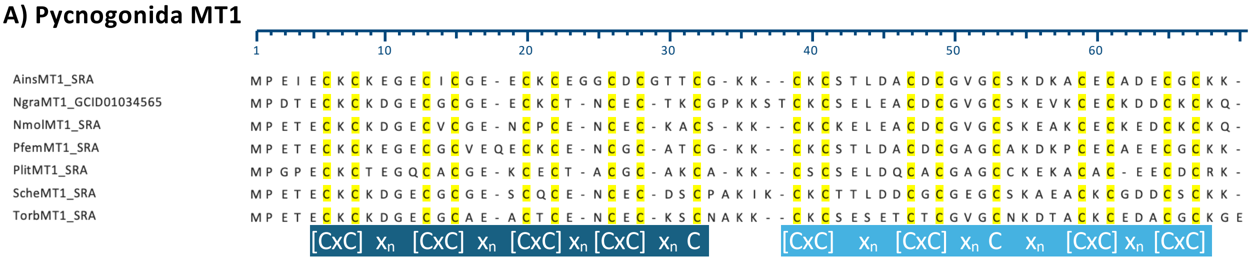


##### B) Merostomata MT1

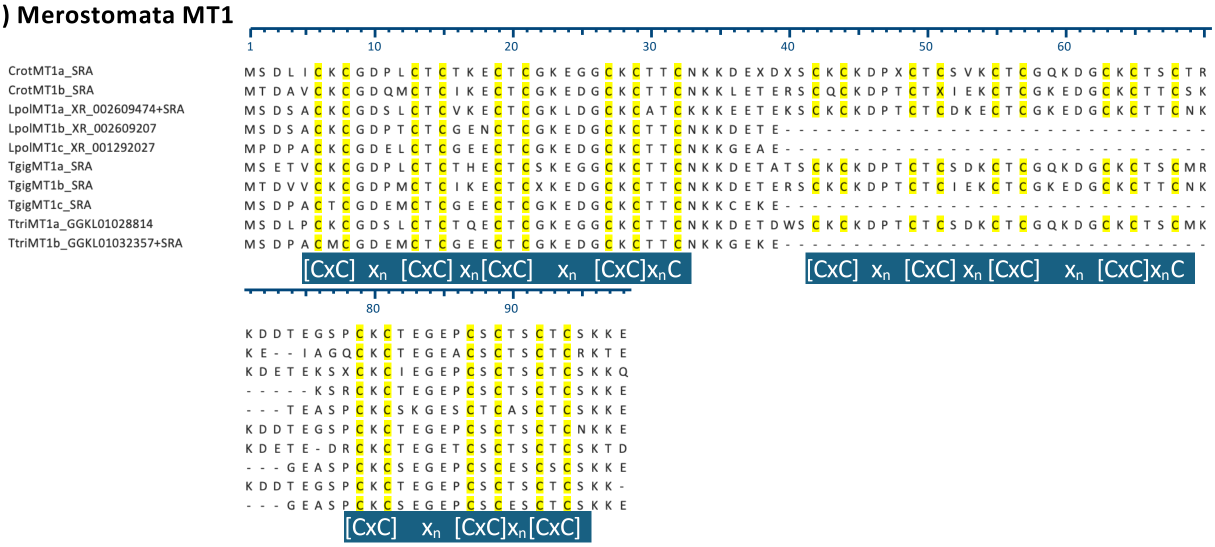


##### C) Arachnida MT1

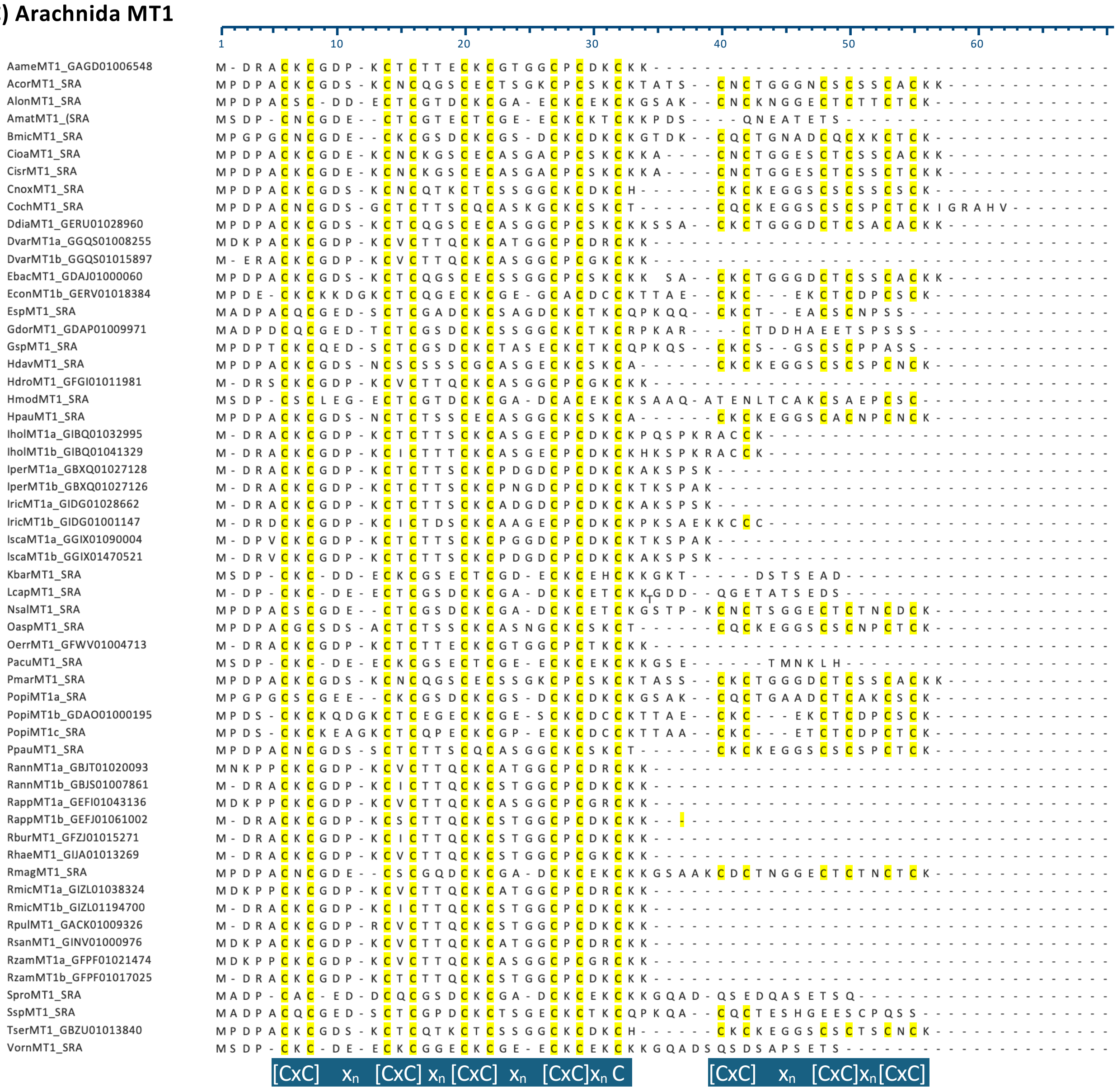

### Figure S4

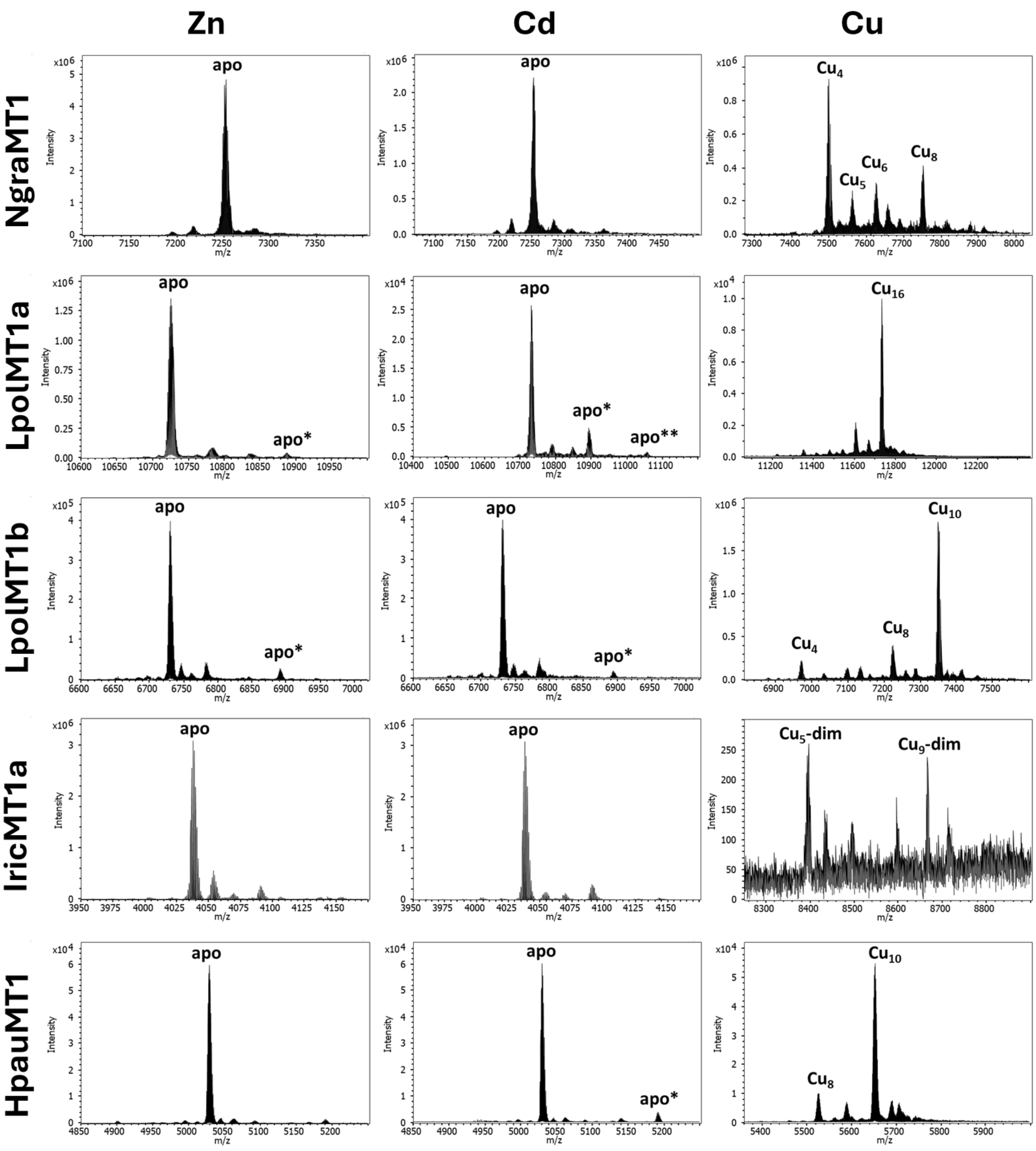

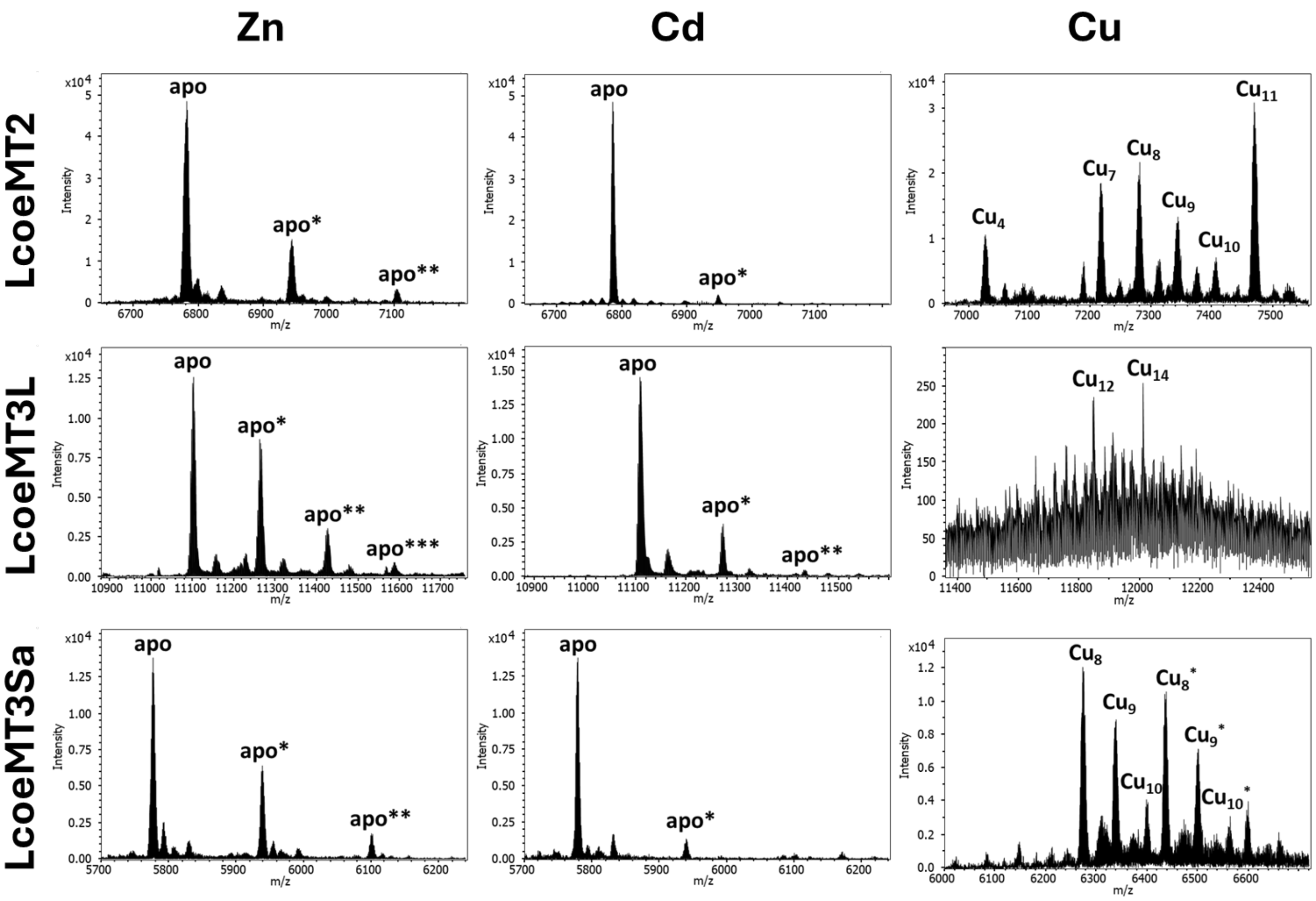
