## Supplementary material for "The Evolutionary History of Metallothioneins in Horseshoe Crabs, Scorpions, Spiders and other Chelicerate Species": Figure S2

### Araneae MT2

```
                              1        10        20        30        40        50        60
Araneomorphae
AaquMT2_GIWR01226499          MADVEKTTSCCTDPKECCKDD-ACCEQGQGP-CHTGKEE-CKDTCH-KK-ACGCKAGEDCKC-SGGKCGC
AargMT2_GIWY01029574          MADVEK-KTCCAEPKECCKDE-SCCQDGKGT-CHTEKEG-CADSCQ-KK-PCGCKAGEDCKC-KDGKCAC
AbruMT2_KAF8785867            MADVEK-KACCAEPKECCKDE-SCCQEGKGP-CHTEKEG-CADSCQ-KK-PCGCKAGEDCKC-KDGKCAC
AdiaMT2_GERP01021930          MADVEK-KSCCAEPKECCKDE-SCCQEGKGP-CHNEKEV-CADSCQ-KK-PCGCKAGEDCKC-KDGKCAC
AipsMT2_IBYV01021947          MADVEK-TSCCTEPKECCKDE-SCCENGQGA-CHTGKEE-CKDSCH-KK-ACGCKAGEECKC-TDGKCGC
AlogMT2_IBSH01016593          MADVEK-KPCCAEPQECCKDE-SCCQEGKGP-CHTDKDG-CADSCQ-KK-LCGCKSGEDCKC-TDGKCAC
AmelMT2_IAPN01004235          MADVEK-KPCCDEPKECCKDE-SCCQEGKGP-CHDEKDG-CADSCQ-KK-PCGCKAGEDCKC-SEGKCAC
AnaeMT2_IBOW01043046          MADVEK-KSSCTEPRECCKDA-SCCDEGKGE-CHTGKEE-CKDTCH-EK-SCGCK--EDCKC-SENKCKC
ApotMT2_IBWF01015665          MADVEK-KSSCTEPRECCKDA-SCCDEGKGE-CHTGKEE-CKDTCH-EK-SCGCK--EDCKC-SENKCKC
AtanMT2_IBAW01019844          MADVEK-TTCCTESKECCKDE-SCCENGQGA-CHTGKEE-CKDSCH-KK-PCGCKTGEECKC-SDGKCGC
CalbMT2_IAZE01025958          MADVEK-KTCCAEPKECCKDE-SCCQEGKGP-CHTEKEG-CADSCQ-KK-PCGCKAGEDCKC-KDGKCAC
CbasMT2_IBFW01007189          MADVEK-TSCCTEPKECCKDE-SCCENGQGA-CHTGKEE-CKDTCH-KK-ACGCKGGEDCKC-SEGKCGC
CdarMT2_GGTX01063157          MADVEK--PCCAEPKECCKDE-SCCDEGKGP-CHTDGKEDCADSCH-KK-PCGCKAGEDCKC-KDGKCAC
CforMT2_IBYP01014520          MADVEN-KSCCAEPKECCKDE-SCCQEGKGP-CHTEKEG-CADSCQ-KK-PCGCKAGEDCKC-QDGKCAC
CfulMT2_IBOC01038815          MADVEK-TSCCTEPKECCKDE-SCCENGQGE-CHTGKEG-CKDTCH-KK-ACGCKADEECKC-TEGKCAC
ClasMT2_IBQA01031776          MADVEK-KSCCAEPKECCKDE-SCCQEGKGP-CHTEKEG-CADSCQ-KK-PCGCKAGEDCKC-TDGKCAC
DcanMT2_ICHF01017029          MADVEK-TSCCTEPKECCKDE-SCCENGQGA-CHTGKEE-CKDTCH-KK-PCGCKAGEDCKC-SEGKCAC
DfolMT2_IAXI01039432          MADVEN-KSCCAEPKECCKDE-SCCQEGKGP-CHTDKDSSCADSCQ-KK-ACGCKSGEDCKC-KDGKCDC
DmarMT2_GIWQ01214392          MADVEKTTSCCTDPKECCKDD-ACCEQGQGP-CHTGKEE-CKDTCH-KK-ACGCKAGEDCKC-SGGKCGC
DsubMT2_ICGF01004055          MADVEK-KSCCAEPKECCKDE-SCCQEGKGP-CHTEKEG-CADSCQ-KK-PCGCKAGEDCKC-TDGKCAC
DtriMT2_GGRN01026674          MADVEK-TSCCTEAKECCKDE-SCCENGQGA-CHTGKEE-CKDSCH-KK-ACGCKAGEDCKC-SEGKCAC
EkitMT2_IBEG01009253          MADVEN-KSCCAEPKECCKDE-SCCQEGKGP-CHTEKEG-CADSCQ-KK-PCGCKAGEDCKC-KDGKCAC
EkumMT2_IANA01004259          MADVEK-TSCCTEPKECCKDE-SCCENGQGE-CHTGKEE-CKDSCH-KK-ACGCKADEECKC-SEGKCAC
EsakMT2_IBEM01035248          MADVEK-KSCCAEPKECCKDE-SCCQEGKGP-CHSEKEG-CADSCQ--K-PCGCKAGEDCKC-KDGKCSC
FgraMT2_IBPA01038432          MADVEK-TSCCTEPKECCKDE-SCCENGQGA-CHTGKEE-CKETCH-KK-PCGCKAGEDCKC-SEGKCAC
GbitMT2_IBVP01006635          MADVEK-KSCCAEPKECCKDE-SCCQEGKGP-CHTEKEG-CADSCQ-KK-PCGCKAGEDCKC-KDGKCAC
HadaMT2_GJQJ01025370          MADVEK-KTCCTEPKECCKDE-SCCQNGQGE-CHTGKEE-CKDSCQ-KK-ACGCKAGEDCKC-SDGKCGC
HmelMT2_IAJG01014351          MADVEK-TSCCTEPKECCKDE-SCCENGQGA-CHTGKEE-CKDSCH-KK-PCGCKEGEDCKC-SDGKCGC
HsanMT2_IATX01006580          MADVEN-KSCCAEPKECCKDE-SCCQDGKGP-CHNEKEG-CADSCQ-KK-ACGCKAGEDCKC-KDGKCGC
HtanMT2_IAQG01046202          MADVEK-KS-CAESKDCCKDDPECCQEGKGP-CHTDKDSSCADSCQ-KK-ACGCKSGEDCKC-KDGKCEC
HvenMT2_IBFQ01010827          MADVEK-TSCCTEPKECCKDE-SCCENGQGA-CHTGKEE-CKDTCH-KK-PCGCKAGEDCKC-SEGKCAC
LcoeMT2_IBIG01002848          MADVEK-TSCCTEPKECCKDE-SCCENGQGA-CHTGKEE-CKDSCH-KK-ACGCKAGEDCKC-SDGKCGC
LcorMT2_IBDW01045585          MADVEK-TSCCTESKECSKDE-SCCENGQGE-CHTGEEK-CKDTCH-KK-ACGCKADEECKC-SEGKCAC
LgeoMT2_GFCZ01010332          MADVEK--PCCPEPKECCKDE-SCCDKGKGP-CHAGEGG--GEACQ-QK-PCGCKGGEECKC-KEGKCGC
LhesMT2_GBJN01124720          MADVEK--PCCPEPKECCKDE-SCCDQGKGP-CHAGEGG--GDACQ-QK-PCGCKAGEDCKC-KEGKCGC
LlaeMT2_EY188643              MADVEK-APCCDSPKECCKDA-TCCDEGKGP-CH-GPDEKCSDSAS-KK-KCGCKAGEDCKC-SEGKCAC
LmaiMT2_ICAK01041707          MADVEK-TSCCTEPKECCKDE-SCCENGQGE-CHTGEEK-CKDTCH-KK-ACGCKADEECKC-AEGKCAC
LthoMT2_ICGP01036048          MADVEN-KSCCAEPKECCKDE-SCCQEGKGP-CHTEKEG-CADSCQ-KK-PCGCKAGEDCKC-QDGKCAC
MbelMT2_IAGN01028376          MADVEK-TPCCTEAKECCKD-----QKGQGE-CHTGKEE-CKDTCH-KK-ACGCKADEECKC-SEGKCAC
MgraMT2_IAWP01018183          MADVEK-KPCCAEPKECCKDE-SCCQEGKGP-CHTEKDG-CADSCQ-TK-PCGCKAGEDCKC-TDGKCSC
MhasMT2_IBKA01027009          MADVEN-KSCCAEPKECCKDE-SCCQEGKGP-CHTEKEG-CADSCQ-KK-PCGCKAGEDCKC-QDGKCAC
MineMT2_ICJC01044708          MADVEK--PCCAEPKECCKDE-SCCQEGKGP-CHTEKEG-CGDSCH-KK-PCGCKAGEDCKC-KDGKCAC
MlemMT2_ICBE01019609          MADVEK-KSCCAEPKECCKDE-SCCQEGKGP-CHTEKDG-CADSCQ-KK-PCGCKAGEDCKC-KDGKCAC
MmusMT2_GJIM01328652          MADVEK-KSCCTEAKECCKDE-SCCQNGQGT-CHTGKEE-CKDSCK-TK-ACGCKAGEDCKC-SDGKCAC
MplaMT2_IBXG01004564          MADVEK-KSCCAEPKECCKDE-TCCQEGKGP-CHTEKE----SCQ-KK-PCGCSTGEECKC-KDGKCAC
MpseMT2_ICBP01032588          MADVEK-K-CCAEPKDCCKEDPSCCQDGKGS-CHTDKDS-CADSCD-KK-ACGCKAGEDCKC-KDGKCAC
MshiMT2_IAKA01018968          MADVEK-TSCCTEAKECCKDE-SCCQNGQGS-CHTGKEE-CKDSCH-KK-PCGCKASEECKC-SDGKCPC
MyawMT2_IBNG01005425          MADVEK-KSDCTESKECCKNE-----MPHEP-CHTDKED-CKDCCK-TK-ECGCKAGEDCKC-SDGKCGC
NaraMT2_IBED01024418          MADVEK-KSCCAEPKECCKDE-SCCQEGKGP-CHTEKEG-CADSCQ-KK-PCGCKAGEDCKC-TDGKCAC
NclaMT2a_GFKT010927754        MADVEK--PCCAEPKECCKDE-SCCDEGKGP-CHTDGKEDCADSCH-KK-PCGCKAGEDCKC-KDGKCAC
NclaMT2b_GFKT012135641        MADVEK-TACCAEPKECCKDE-SCCQEGKGP-CHTEKES-CGDSCH-KK-PCGCKAGEDCKC-KDGKCAC
NcruMT2_GEWZ01004248          MADVEK-TACCAEPKECCKDE-SCCQEGKGP-CHTEKES-CGDSCH-KK-PCGCKAGEDCKC-KDGKCAC
NmunMT2_IAVV01016525          MADVEK--PCCAEPKECCKDE-SCCQEGKGP-CHTEKEG-CADSCQ-KK-PCGCKAGEDCKC-KDGKCAC
NyesMT2_IBBO01018781          MADVEK-K-CCDEPKECCKDE-SCCQEGKGS-CHTEKDGSCADKCHAKK-ACGCTAGEECKC-KEGKCGC
OgraMT2_IBKN01001004          MADVEK--SCCAETKDCCDVDANCCENGKGT-CHEKDADCCKDGCHTKK-PCGCKANEACKC-SEGKCAC
OgrpMT2_ICKD01013852          MADVEK-TSCCTEAKECCKDE-SCCENGQGA-CHTGKEE-CKDSCH-KK-ACGCKAGEDCKC-SEGKCAC
OhosMT2_IBYW01036832          MADVEK-TSCCTESKECCKDE-SCCENGQGE-CHTGKEE-CKDTCH-KK-ACGCKADEECKC-SEGKCAC
OlunMT2_ICGC01035958          MADVEK-TSCCTDAKEC-KDD-SCCDNGQAA-CHTGKEE-CAETCH-KK-PCGCKAGEDCKC-SDGKCAC
OmatMT2_IBFR01041374          MADVEK-TSCCTETKECCKDE-SCCENGQGE-CHTGKEG-CKDTCH-KK-ACGCKAGEECKC-TEGKCAC
OnavMT2_IBWY01033478          MADVEN-KSCCAEPKECCKDE-SCCQEGKGP-CHTEKEG-CADSCQ-KK-PCGCKAGEDCKC-QDGKCAC
OokiMT2_IALW01017951          MADVEK--SCCAETKDCCDVDANCCENGKGT-CHEKDADCCKDGCHTKK-PCGCKANEACKC-SEGKCAC
OsybMT2_IAZZ01001752          MADVEK--SCCAETKDCCDVDANCCENGKGT-CHEKDADCCKDGCHTKK-PCGCKANEACKC-SEGKCAC
OvarMT2_IAST01008113          MADVEK--SCCAETKDCCDVDANCCENGKGT-CHEKDADCCKDGCHTKK-PCGCKANEACKC-SEGKCAC
OyesMT2_IAQA01003517          MADVEK--SCCAETKDCCDVDANCCENGKGT-CHEKDADCCKDGCHTKK-PCGCKANEACKC-SEGKCAC
PameMT2_GJIU01181368          MADVEK-TSCCTESKECCKDE-SCCENGQGA-CHTGKEE-CKDSCH-KK-ACGCKAGEDCKC-SDGKCGC
PbicMT2_ICDN01011823          MADVEK-TSCCTESKECCKDD-SCCENGKAA-CHAGKEE-CKESCHQKDSPCKCKSGEECK--KEG-CSC
PbreMT2_IAWA01034742          MADVEK-TSCCTESKECCKDE-SCCENGQGA-CHTGKEE-CKDSCH-KK-ACGCKADEDCKC-AEGKCAC
PceyMT2_IAYL01007552          MADVEK-TSCCTEPKECCKDE-SCCENGQGK-CHTGKEE-CKDTCH-KK-SCGCKADEECKC-SEGKCAC
PclaMT2_ICBG01047316          MADVEK-KSCCTEPKECCKDE-SCCQNGQGS-CHTGKEE-CKDSCQ-TK-ACGCKAGEDCKC-SDGKCAC
PemaMT2_IBPM01002942          MADVEK-TSCCTEPKECCKDE-SCCQNGQGA-CHTDKEE-CKESCH-KK-PCGCKTGEECKCSSDGKCAC
PjapMT2_IAZW01000630          MADVEN-KSCCAEPKECCKDE-SCCQEGKGP-CHTDKDSSCADSCQ-KK-ACGCKSGEDCKC-KDGKCDC
PkatMT2_IAOS01018527          MADVEK-TSCCTEPKECCKDE-SCCENGQGE-CHTGKEE-CKDTCH-KK-ACGCKADEECKC-SEGKCAC
PlamMT2_IAQF01020027          MADVEK-KACCAEPKECCKDE-SCCQEGKGP-CHTEKEG-CADSCQ-KK-PCGCKAGEDCKC-KDGKCAC
PnigMT2_GFUY01017528          MADVEK-TSCCTEQKECCKDE-SCCENGQGA-CHTGKEE-CKDTCH-KK-ACGCKAGEDCKC-SEGKCAC
PpilMT2_IBOK01000603          MADVEK-K-CCAEPKDCCKEDPSCCQDGKGS-CHTDKDS-CADSCD-KK-ACGCKAGEDCKC-KDGKCAC
PpseMT2_GGRD01042272          MADVEK-TSCCTEPKECCKDE-SCCENGQGA-CHTGKEE-CKDSCH-KK-PCGCKEGEDCKC-SDGKCGC
PquaMT2_IALH01014478          MADVEK--PCCAEPKECCKDE-SCCQEGKGP-CHTEKEG-CGDSCH-KK-PCGCKAGEDCKC-KDGKCAC
PryuMT2a_IBPF01015651         --DVEK-TSCCTEPKECCKDE-SCCENGQGA-CHTGKEE-CKDSCH-KK-PCGCKEGEDCKC-SDGKCGC
PryuMT2b_ICDI01032605         MADVEK--PCCDEPKDCCKDE-SCCKEGKGP-CHTEKEG-CGDSCH-KK-ACGCKVGEDCKC-SGGKCEC
PryuMT2c_ICDI01004733         MADVEK-KPCCAEPKECCKDE-SCCQEGKGP-CHTEKEG-CADSCQ-KK-PCGCKAGEDCKC-QDGKCAC
PspiMT2_IBQV01000539          MADVEK-TSCCTETKECCKDE-SCCENGQGA-CHTGKEE-CKDSCH-KK-PCGCKAGEDCKC-SEGKCAC
PtepMT2_IBLE01032418          MADVEN-KSCCAEPKECCKDE-SCCQEGKGP-CHTDKDSSCADSCQ-KK-ACGCKSGEDCKC-KDGKCDC
PterMT2_IBAD01021535          MADVEN-KSCCAETKECCKDE-SCCQEGKGP-CHTEKEG-CADSCQ-KK-PCGCKAGEDCKC-KDGKCAC
SannMT2_IBPH01030142          MADVEK-KSCCTEGKECCKDE-SCCENGEGA-CHKGKEE-CKESCH-KK-PCGCKAGEDCKC-TDGKCGC
SdumMT2_XR_004757882+SRA      MADVEK-TGCCAEAKECAADN--CSTECKTN-CHDGKEE-CKDSCH-KK-PCGCKGTEDCKC-SEGKCGC
SforMT2_IAZL01034704          MADVEK-TSCCTESKECCKDE-SCCENGQGA-CHTGKEE-CKDSCH-KK-PCGCKAGEDCKC-SEGKCAC
SgroMT2_GBJQ01091123          MADVEK--PCCTEPKECCKDE-SCCDEGKGP-CHANEGG--DAACQ-QK-PCGCKSGEDCKC-KEGKCGC
SlinMT2_JT031711              MADVEK-TGCCAEAKECAADN--CSTECKTN-CHDGKEE-CKDSCH-KK-PCGCKGTEDCKC-SEGKCGC
SmimMT2_JT038942              MADVEK-TGCCAEAKECAADN--CSTECKTN-CHDGKEE-CKDSCH-KK-PCGCKGTEDCKC-SEGKCAC
SocrMT2_GIZV01010417          MADVEK-TSCCTESKECCKDE-SCCENDQGA-CHTGKEE-CKDTCH-KK-ACGCKAGEDCKC-TDGKCGC
SoriMT2_IBGW01039022          MADVEK-KSCCAEPKECCKDE-SCCQEGKGP-CHTEKEG-CADSCQ-KK-PCGCKAGEDCKC-QDGKCAC
StenMT2_JT042210              MADVEK-TGCCAEAKECAADN--CSTECKTN-CHDGKEE-CKDSCH-KK-PCGCKGTEDCKC-SEGKCGC
TangMT2_IBHB01033618          MADVEK-TSCCTEPKECCKDE-SCCQNGQGE-CHTGKAE-CKDTCQ-KK-ACGCKADEECKC-SEGKCAC
TaquMT2_IBZV01030133          MADVEK-TSCCTESKECCKDE-SCCENGQGA-CHTGKEE-CKDSCH-KK-ACGCKAGEDCKC-SDGKCGC
TbreMT2_GITM01027747          MADAEK--SCCAEPKECCKDE-SCCEEGKGP-CHTEGKG-CEDTCH-QK-PCGCKSGEECKCTSGSKCAC
TfenMT2_IAVL01019861          MADVEK-TTGCTEPKECCKDE-ACCENGQGA-CHDGKEE-CKDSCH-KK-ACGCKAGEDCKCSSDGKCAC
TlabMT2_IAWV01026404          MADVEK-TSCCTESKECCKDE-SCCENGQGE-CHTGKEE-CKDTCH-KK-ACGCKADEECKC-AEGKCAC
TokiMT2_ICDD01000078          MADVEK-TSCCTEPKECCKDE-SCCENGKGE-CHTGKEE-CKDTCH-KK-ACGCKADEECKC-SEGKCAC
TperMT2_GGOF01120677          MADVEK-KTCCAEPKECCKDE-SCCQDGKGT-CHTEKEG-CADSCQ-KK-PCGCKAGEDCKC-KDGKCAC
TrimMT2_IAWZ01003745          MADVEK-TSCCTEPKECCKDE-SCCQNGQGE-CHSGKEE-CKDTCH-KK-ACGCKADEECKC-SEGKCAC
TterMT2_IBIR01014579          MADVEK-TSCCTEPKECCKDE-SCCENGQGT-CHTGKEE-CKDSCH-KK-ACGCKAGEDCKC-SDGKCAC
UdivMT2_GJZJ01180553          MADVEK-KACCAEPKECCKDE-SCCQDGKGA-CHTEKEG-CADSCQ-KK-PCGCKAGEDCKC-KDGKCAC
VspMT2_ICOO01003872           MADVEK-TSCCTEAKECCKDE-SCCENGQGA-CHTGKEE-CKDTCH-KK-ACGCKAGEDCKC-SDGKCAC
ZasiMT2_IAGO01029205          MADVEK-TSCCTEQKECCKDE-SCCENGQGA-CHTGKEE-CKDSCH-KK-ACGCKGGEDCKC-TDGKCGC
Mesothelae
HhelMT2_IAMB01023112          MADTEK-TACCTAEKECCKDS-SCCEEGKGKECCDNNPDCCKDGAA-KK-PCGCKEGEECKC-KEGKCAC
LmurMT2_ICAH01033317          MADAEK-TACCTADKECCKDS-SCCEEGKGKECCDNNPDCCKEGAA-KK-PCGCKEGEECKC-QEGKCAC
RnisMT2a_IAQS01038821         MADTEK-TACCTAEKECCKDS-SCCEEGKGKECCDNNPDCCKDGAA-KK-PCGCKEGEECKC-QEGKCAC
RnisMT2b_IAIP01007631         MADTET-TACCTAEKQCCKDS-SCCEQGKGKECCDNHPDCCKDGAA-QK-PSGCKEGEECKS-QEGKCAC
Mygalomorphae
ArorMT2_IBPI01014909          MADVEK-KSCCSPEKECCKEN-ECTAEGTE--CCHKDSDGCKDGAVQKK-PCGCKVGEDCKC-KEGKCAC
CfraMT2_IAVG01026659          MADVEK-TSCCTTEEDCCKGG-DCTTEGKD--CS-KEHDCCKDGSA-TK-PCGCKQGEDCKC-KDGKCGC
CsigMT2_IBXS01010418          MADVEK-NTCCSPEKECCKEN-ECTAEGKE--CCHGDTECCKDGAAQKK-PCGCKQGEDCKC-QDGKCAC
MgigMT2_IAEV01010058          MADVEK-TACCTTEEDCCKGS-ECTTEGKD--CS-KEHDCCKDGSA-QK-PCGCKKGEDCKC-QEGKCAC
RatrMT2a_GIXJ01248059         MADVEK-TSCCTETKECCKDE-SCCENGQGA-CHTGKEE-CKDSCH-KK-PCGCKAGEDCKC-SDGKCGC
RatrMT2b_GIXJ01248060         MADVEK-TSCCTEPKECCKDE-SCCENGQGS-CHTGKEE-CKDTCH-KK-ACGCKAGEDCKC-SEGKCAC
```

CC $x_n$ CC $x_n$ CC $x_n$ CH     C $x_n$ C $x_n$ [CxC] $x_n$ [CxC] $x_n$ [CxC]
