## Supplementary material for "The Evolutionary History of Metallothioneins in Horseshoe Crabs, Scorpions, Spiders and other Chelicerate Species": Figure S3

**Figure S3A**

**A) Araneae MT3S**

(Position ruler: 1, 10, 20, 30, 40, 50, 60)

| Sequence | Alignment |
|---|---|
| **Araneomorphae** | |
| AaquMT3S_GIWR01171385 | MPGGPCGSNCTCER-CRCCSAA------TKCNSGCRCETCRCCQRNQ-KCECGDCKCCGAAKC |
| AargMT3S_GIWY01109359 | MPGGPCGANCNCEK-CRCCSAA------SKCTANCKCDTCRCCQRNQ-QCNCGDCKCCSSAKC |
| AdepMT3S_ICFY01031203 | MPCGTCGPSCTCDK-CRCCGAS----SGSKCGPNCKCADCKCCGKSSANCRCGDCKCCGAAKC |
| AfusMT3S_IBZC01022604 | MPCGTCGPNCSCEK-CRCCGAA----AAPKCSANCQCGDCKCCGRTSANCRCGDCKCCGAAKC |
| AipsMT3S_IBYV01029187 | MPGGPCGANCTCEK-CRCCSAA------SKCSSGCRCDTCRCCQRGQ-KCQCGDCKCCGAAKC |
| AlogMT3Sa_IBSH01031123 | MPCGTCGSSCTCDK-CRCCGAS----SDSKCSANCQCVDCKCCGRSSATCRCGDCKCCSASKC |
| AlogMT3Sb_IBSH01034144 | MPGGPCGSNCTCEK-CRCCSAA------SKCTASCKCDTCRCCQRNQ-KCECGDCKCCSASKC |
| AmelMT3S_IAPN01022659 | MPCGTCGSNCTCDK-CRCCGAA----SDSKCSANCKCVDCKCCGRSSANCRCGDCKCCSAAKC |
| AnaeMT3S_IBPA01019018 | MPG-PCGSNCNCDK-CRCCSAA------SKCGANCKCDNCRCCQRGGQKCECGDCKCCGAAKC |
| AnepMT3S_IBFL01003327 | MPCGTCGPNCTCDK-CRCCGAA----ASSKCSANCNCGDCKCCGRTSANCRCGACKCCGAAKC |
| ApotMT3S_IBWF01018135 | MPGGPCGSNCTCEK-CRCCSAA------SKCTASCKCDTCRCCQRNQ-KCECGDCKCCSASKC |
| AtanMT3S_IBAW01043010 | MPCGTCGPNCTCEK-CRCCSAA------AKCSANCKCDDCKCCGRGQ-NCRCGDCKCCGAAKC |
| AyagMT3S_IAYO01029315 | MPCGTCGPSCTCEK-CRCCGAS----AAPKCSANCQCGDCKCCGRTSANCRCGDCKCCGAAKC |
| BgraMT3S_IAMX01033138 | MPCGTCGPNCTCEK-CRCCGAA----AAPKCSANCNCGDCKCCGRTSANCRCGDCKCCGAAKC |
| BlonMT3S_GIWL01016652 | MPGGPCGTNCTCDK-CRCCSAA------TKCSSGCKCDTCRCCQRGQ-KCECGDCKCCGAAKC |
| CakiMT3S_ICFL01015310 | MPCGTCGASCTCEK-CRCCGAS----ASSKCSENCKCGDCRCCGRTSANCRCGDCKCCGAAKC |
| CalbMT3Sa_IAZE01015529 | MPCGTCGPSCTCDK-CRCCGAS----SGPKCSANCNCGDCKCCGRTQANCRCGDCKCCGAAKC |
| CalbMT3Sb_IAZE01014351 | MPGGPCGSNCNCEK-CRCCSAA------SKCTANCKCDTCRCCQRNQ-QCNCGDCKCCSSAKC |
| CbasMT3S_IBFW01014451 | MPGGPCGTNCTCDK-CRCCSAA------AKCSSGCKCDTCRCCQRSQ-KCQCGDCKCCGASKC |
| CcicMT3S_IBHY01004928 | MPCGTCGPSCSCEK-CRCCGAA----ASSKCSANCKCVDCKCCGRSSANCRCGDCKCCGAAKC |
| CdaMT3S_GIY23593 | MPCGTCGPSCSCDK-CRCCGAA--ASASAKCSANCKCDDCRCCGRSGANCRCGDCKCCSAAKC |
| CextMT3S_GIY05480 | MPCGTCGPSCSCDK-CRCCGAA--ASASAKCSANCKCDDCRCCGRSGANCRCGDCKCCSAAKC |
| CforMT3S_IBYP01019334 | MPCGTCGPNCTCDK-CRCCGAA----ASSKCSANCNCGDCKCCGRTSANCRCGDCKCCGAAKC |
| CfulMT3S_IBOC01006221 | MPGGPCGSNCTCEK-CRCCTAA------SKCTSSCKCDTCRCCQRSQ-KCNCGDCKCCGAAKC |
| ClasMT3S_IBOH01015086 | MPGGPCGSNCTCEK-CRCCSAA------TKCSSGCKCDNCRCCQRGQGQCQCGDCKCCGAAKC |
| DcanMT3S_ICHF01007191 | MPGGPCGSNCKCEK-CRCCSAA------SKCGANCKCDTCRCCQRGQ-KCECGDCKCCGAAKC |
| DfolMT3Sa_IAXI01008702 | MPGGPCGSNCTCEK-CRCCSAA------AKCGSSCKCDTCRCCQRGQ-RCECGDCKCCGAAKC |
| DfolMT3Sb_IAXI01012824 | MNCGTCGPNCTCEK-CRCCGAA----AAPKCSANCNCGDCKCCGRTSANCRCGDCKCCGAAKC |
| DmarMT3S_GIWQ01313974 | MPGGPCGTNCTCEK-CRCCSAA------AKCSPNCKCDTCRCCQRNQ-KCECGDCKCCGAAKC |
| DsubMT3S_IBBD01042702 | MPGGPCGSNCTCEK-CRCCSAA------SKCTASCKCDTCRCCQRNQ-KCECGDCKCCSASKC |
| DtriMT3S_GGRN01043194 | MPGGPCGSNCTCEK-CRCCSAA------AKCSSGCRCDTCRCCQRGQ-KCQCGDCKCCGAAKC |
| EabrMT3S_IBGU01031423 | MPCGTCGPNCTCEK-CRCCGAA----AAPKCSANCNCGDCKCCGRTSANCRCGDCKCCGAAKC |
| EkitMT3Sa_IBEG01030932 | MPCGTCGPNCTCEK-CRCCGAA----AAPKCSANCNCGDCKCCGRTSANCRCGDCKCCGAAKC |
| EkitMT3Sb_IBEG01022842 | MPGGPCGSNCTCEK-CRCCSAA------SKCTASCKCDTCRCCQRNQ-KCECGDCKCCSASKC |
| EkitMT3Sc_IBEG01030305 | MA-GPCGANCTCDK-CRCCSAA------SKCTASCKCDTCRCCQRSQ-NCNCGDCKCCGAAKC |
| EkitMT3Sd_IBEG01032665 | MP-GPCGENCKCEK-CRCCSAA------SKCTSNCKCDVCKCCERSQ-KCNCGDCKCCGEAKC |
| EkumMT3S_IANA01042734 | MPGGPCGSNCSCEK-CRCCSAA------SKCTASCKCDTCRCCQRGQ-KCNCGDCKCCGAAKC |
| EsakMT3S_IBEM01002195 | MPCGTCGPSCTCEK-CRCCGAA----AAPKCSANCQCGDCKCCGRSSANCRCGDCKCCGAAKC |
| FgraMT3S_IBPA01019018 | MPGGPCGPNCTCER-CRCCSAA------VRCGAGCKCDTCKCCQRGE-KCQCGDCKCCRAAKC |
| GbitMT3S_IBVP01005554 | MPCGTCGPNCTCEK-CRCCGAA----AAPKCSANCNCGDCKCCGRTSANCRCGDCKCCGAAKC |
| HadaMT3S_GJQJ01012079 | MPGGPCGSNCTCEK-CRCCSAA------SKCSSGCKCDTCRCCQRNQ-SCNCGDCKCCRAAKC |
| HmelMT3S_IAJG01036463 | MPGGPCGSNCTCEK-CRCCSAA------SKCTSSCKCDTCRCCQRSQ-KCNCGDCKCCGAAKC |
| HsanMT3S_IATX01035236 | MPCGTCGPSCTCEK-CRCCGAG--AASASKCSANCKCADCKCCGRTSANCRCGDCKCCDAAKC |
| HtanMT3Sa_IAQG01038721 | MPCGTCGPNCTCEK-CRCCGAA----AAPKCSANCNCGDCKCCGRTSANCRCGDCKCCGAAKC |
| HtanMT3Sb_IAQG01034859 | MPGGPCGSNCTCEK-CRCCSAA------AKCTSSCKCDTCRCCQRSQ-KCECGDCKCCSASKC |
| HvenMT3S_IAGF01017184 | MPGGPCGSNCTCEK-CRCCSAA------SKCGPGCRCDNCRCCQRGQ-NCQCGDCKCCGAAKC |
| LcoeMT3Sa_IBVM01020795 | MPGGPCGSNCTCEK-CRCCSAA------SKCSAGCRCDTCRCCQRGQ-KCQCGDCKCCGAAKC |
| LcoeMT3Sb_IBIG01009944 | MPGGPCGANCSCEK-CRCCSAA--ASSGGRCGPNCRCDVCRCCNRGQ-NCRCGDCRCCGASKC |
| LcorMT3Sa_IBDW01030729 | MPGGPCGSNCTCEK-CRCCTAA------SKCTASCKCDTCRCCQRSQ-KCNCGDCKCCGAAKC |
| LcorMT3Sb_IBDW01005672 | MPGGPCGSNCSCEK-CRCCSAA-ASASPSKCGPNCKCDVCRCCNRGQ-NCRCGDCKCCGASKC |
| LcornMT3S_IBUI01018852 | MPCGTCGPSCTCEK-CRCCGAA---ATSAKCSANCKCVDCKCCGKTGANCRCGDCKCCGAAKC |
| LgeoMT3S_GBJM01039547 | MP-GPCGPNCNCEK-CRCCSAA------SKCTSNCKCDVCRCCSRSQ-RCNCGDCKCCGEAKC |
| LhesMT3Sa_GBCS01027129 | MP-GPCGPNCNCEK-CRCCSAA------SKCTSSCKCDVCHCCSRSR-RCNCGDCKCCGEAKC |
| LhesMT3Sb_GBJN01094922 | MP-GPCGPNCNCEK-CRCCSAA------SKCTSNCKCDVCRCCSRNQ-KCNCGDCKCCGEAKC |
| LmaiMT3Sa_ICAK01036021 | MPGGPCGSNCTCEK-CRCCTAA------SKCTASCKCDTCRCCQSGQ-KCNCGDCKCCGAAKC |
| LmaiMT3Sb_ICAK01032991 | MPGGPCGSNCSCEK-CRCCSAA-ASASPSKCGPNCKCDVCRCCNRGQ-NCRCGDCKCCGASKC |
| LthoMT3S_ICGP01002053 | MPCGTCGPNCTCDK-CRCCGAA----ASSKCSANCNCGDCKCCGRTSANCRCGDCKCCGAAKC |
| MbelMT3Sa_IAGN01029399 | MPGGPCGSNCTCEK-CRCCSAA------SKCTASCRCDTCRCCQRSQ-KCNCGDCKCCGAAKC |
| MbelMT3Sb_IAGN01017054 | MPGGPCGANCSCEK-CRCCSAA-ASASASRCGPNCKCDVCRCCNKGQ-NCRCGDCKCCGASKC |
| MgraMT3S_IAWP01035947 | MPCGTCGPSCSCEK-CRCCGAA----ASSKCSPNCKCDNCQCCGRSSANCRCGDCKCCGASKC |
| MhasMT3S_IBKA01028729 | MPCGTCGPNCTCDK-CRCCGAA----ASSKCSANCNCGDCKCCGRTSANCRCGDCKCCGAAKC |
| MineMT3S_ICJC01041083 | MPCGTCGPNCTCEK-CRCCGAA----AAPKCSANCNCGDCKCCGRTSANCRCGDCKCCGAAKC |
| MlemMT3S_ICBE01013695 | MPCGTCGPSCSCEK-CRCCGAA---AAAPKCSANCKCVDCKCCGKTGANCRCGDCKCCGAAKC |
| MmusMT3S_GJIM01320864 | MPGGPCGSNCTCEK-CRCCSAA------TKCNSGCKCDPCRCCGKGQ-NCRCGDCKCCGAAKC |
| MplaMT3S_IBXG01043951 | MPCGTCGPNCTCEK-CRCCGAA---ASSAKCGPNCKCVDCKCCGKTSANCRCGDCKCCGAAKC |
| MpseMT3Sa_ICBP01045714 | MPGGPCGTNCTCEK-CRCCGAA------AKCSASCKCETCRCCQRGQ-NCRCGDCKCCGASKC |
| MpseMT3Sb_ICBP01037319 | MP-GPCGPNCTCDK-CRCCTAA------TKCSEGCKCDTCRCCERGQ-KCQCGDCKCCGAAKC |
| MshiMT3S_IAKA01005514 | MPGGPCGPNCSCEK-CRCCSAS----AAPKCSASCKCDKCRCCQRTQ-KCQCGDCKCCAGSTC |
| MyawMT3S_IBNG01051378 | MPGGPCGSNCTCEK-CRCCSAA------TKCNSGCKCDPCRCCGKGQ-NCRCGDCKCCGAAKC |
| NaraMT3S_IBED01005087 | MPCGTCGPSCTCDK-CRCCGAA----AAPKCSANCQCGDCKCCGRTSATCRCGDCKCCGAAKC |
| NclaMT3S_GFKT011191296 | MPGGPCGSNCNCEK-CRCCSAA------SKCTSGCKCDTCRCCQRNQ-KCECGDCKCCGAAKC |
| NmunMT3S_IAVV01005513 | MPCGTCGPNCTCEK-CRCCGAA----AAPKCSANCNCGDCKCCGRTSANCRCGDCKCCGAAKC |
| NumbMT3S_IAHU01040313 | MPCGTCGPSCTCEK-CRCCGAA--TASASKCSANCKCADCKCCGRTSANCRCGDCKCCGAAKC |
| NyesMT3S_IBBO01028050 | MPCGTCGPNCTCEK-CRCCGAA----AAPKCSANCNCGDCKCCGRTSANCRCGDCKCCGAAKC |
| OgraMT3Sa_ICKD01006318 | MHGGPCGANCTCEK-CRCCGAA------SRCSDTCKCDTCRCCQKGR-DCRCGDCKCCGASKC |
| OgraMT3Sb_ICDQ01018272 | -PGGPCGANCTCEK-CRCCGAA------SRCSDTCKCDTCRCCQKGR-DCRCGDCKCCGASKC |
| OgraMT3Sc_ICDQ01030917 | MPGGPCGANCTCEK-CRCCGAA------AKCSSSCKCETCRCCQRGQ-NCRCGDCKCCGASKC |
| OhosMT3Sa_IBIV01016281 | MPGGPCGPNCACEK-CRCCSAA------SKCTAACKCDTCRCCQRNQ-KCNCGDCKCCGASKC |
| OhosMT3Sb_ICJU01039914 | MPGGPCGANCTCEK-CRCCSSA-ATASASRCGPNCKCDVCRCCGRGQ-NCRCGDCKCCSASKC |
| OlunMT3S_ICGC01018723 | MPGGPCGSNCTCEK-CRCCSAA------SKCSAGCKCDTCRCCQRGE-KCQCGDCKCCGAAKC |
| OmatMT3S_IBFR01032300 | MPCGTCGPNCTCEK-CRCCGAA----AAPKCSANCNCGDCKCCGRTSANCRCGDCKCCGAAKC |
| OnavMT3S_IBWY01005004 | MPCGTCGPNCTCDK-CRCCGAA----ASSKCSANCNCGDCKCCGRTSANCRCGDCKCCGAAKC |
| OokiMT3Sa_IALW01006618 | MTGGTCGANCTCEK-CRCCGAA------SRCSDTCKCDTCRCCQKGR-DCRCGDCKCCGASKC |
| OokiMT3Sb_IALW01034307 | MPGGPCDPNCTCEK-CRCCGAA------AKCSSSCKCETCRCCQRGQ-NCRCGDCKCCGASKC |
| OokiMT3Sc_IALW01003257 | MPGGPCGANCTCEK-CRCCGAA------AKCSSSCKCETCRCCQRGQ-NCRCGDCKCCGASKC |
| OsybMT3Sa_IAUX01017549 | MPGGPCGANCTCEK-CRCCGAA------SRCSDTCKCDTCRCCQKGR-DCRCGDCKCCGASKC |
| OsybMT3Sb_IAUX01008602 | MPGGPCGANCTCEK-CRCCGAA------AKCSASCKCETCRCCQRGQ-NCRCGDCKCCGASKC |
| OvarMT3Sa_IAST01011549 | MPGGPCGANCTCEK-CRCCGAA------SRCSDTCKCDTCRCCQKGR-DCRCGDCKCCGASKC |
| OvarMT3Sb_IAST01017713 | MPGGPCGSNCTCEK-CRCCGAA------AKCSASCKCETCRCCQRGQ-NCRCGDCKCCGASKC |
| OyesMT3Sa_IAQA01022141 | MPGGPCGANCTCEK-CRCCGAA------SRCSDTCKCDTCRCCQKGR-DCRCGDCKCCGASKC |
| OyesMT3Sb_IAQA01008891 | MPGGPCGTNCTCEK-CRCCGAA------AKCSASCKCETCRCCQRGQ-NCRCGDCKCCGASKC |
| PameMT3S_GJIU01318898 | MPGGPCGANCSCEK-CRCCSAA--ASSGGRCGPNCRCEVCRCCNRGQ-NCRCGDCRCCSASKC |
| PastMT3S_IBUD01021694 | MPCGTCGPSCTCEK-CRCCGAA----AAPKCSSNCQCGDCKCCGKASASCRCGDCKCCSASKC |
| PausMT3S_ICKB01002344 | MPCGTCGPNCTCDK-CRCCGAA----ASSKCSANCNCGDCKCCGRTSANCRCGDCKCCGAAKC |
| PbicMT3S_ICDN01028901 | MPGGPCGSNCNCEK-CRCCSAA--SASDSRCGPNCRCDVCRCCKRGA-NCRCGDCRCCSASKC |
| PbreMT3S_IAWA01017274 | MPGGPCGANCSCEK-CRCCSAAGACGSSSRCSPNCKCDVCRCCNRGQ-NCRCGDCKCCSASKC |
| PceyMT3S_IAYL01023434 | MPGGPCGSNCSCEK-CRCCSAA-ASASASRCGSNCKCDVCRCCNRGQ-NCRCGDCKCCSASKC |
| PemaMT3S_IBPM01033659 | MPCGTCGPNCTCEK-CRCCGAA----AAPKCSANCNCGDCKCCGRTSANCRCGDCKCCGAAKC |
| PjapMT3S_IAZW01024263 | MPCGTCGPNCTCEK-CRCCGAA----AAPKCSANCNCGDCKCCGRTSANCRCGDCKCCGAAKC |
| PkatMT3S_IAOS01012376 | MPGGPCGSNCSCEK-CRCCSAA-ASASASRCGSNCKCDVCRCCNRGQ-NCRCGDCKCCSASKC |
| PlamMT3S_IAQF01007099 | MPCGACGPNCTCEK-CRCCGAA----AAPKCSANCNCSDCKCCGRTSANCRCGDCKCCSAAKC |
| PnigMT3S_GFUY01018717 | MPGGPCGANCTCER-CRCCSAA------TKCSSGCKCDSCRCCQRGQ-NCQCGDCKCCGAAKC |
| PpilMT3S_IBOK01019512 | MPGGPCGPNCTCEK-CRCCTAA------SKCTSNCKCDNCRCCQRSQ-KCNCGDCKCCGEAKC |
| PpseMT3S_GGRD01193535 | MPGGPCGSNCTCEK-CRCCSAA------SKCSAGCRCDTCRCCQRGQ-KCQCGDCKCCGAAKC |
| PquaMT3Sa_IALH01010477 | MPCGTCGPNCTCEK-CRCCGAA----AAPKCSANCNCGDCKCCGRTSANCRCGDCKCCGAAKC |
| PquaMT3Sb_IALH01010495 | MPGGPCGSNCTCEK-CRCCSAA------SKCTASCKCDTCRCCQRNQ-KCECGDCKCCSASKC |
| PryuMT3S_IAVY01004002 | MPCGTCGPNCTCEK-CRCCGAA----AAPKCSANCNCGDCKCCGRTSANCRCGDCKCCGAAKC |
| PspiMT3S_IBKY01029357 | MPGGPCGSNCTCEK-CRCCSAA------SKCSPSCKCDTCRCCQRGQ-KCQCGDCKCCGAAKC |
| PsteMT3S_IAGQ01013489 | MNCGTCGPNCTCEK-CRCCGAA----AAPKCSANCNCGDCKCCGRTSANCRCGDCKCCGAAKC |
| PtepMT3S_IACA01037330 | MPGGPCGPNCTCEK-CRCCSAA------SKCTSNCKCDNCRCCQRSQ-KCNCGDCKCCGEAKC |
| PterMT3S_IBAD01018297 | MPCGTCGPNCSCEK-CRCCGAA----SASKCSANCKCSDCKCCGRTSANCRCGDCKCCDAAKC |
| SannMT3S_IBPH01041663 | MPCGTCGPNCTCEK-CRCCGAA----AAPKCSANCNCGDCKCCGRTSANCRCGDCKCCGAAKC |
| SforMT3S_IAZL01014300 | MPCGTCGSNCTCEK-CRCCSAA-----ASKCGPGCKCDNCKCCQRSQ-KCQCGDCRCCGAAKC |
| SgroMT3S_GFDC01029402 | MPGGPCGPNCNCEK-CRCCSAA------SKCTSNCKCDVCRCCSRGQ-KCNCGDCKCCGEAKC |
| SlinMT3Sa_JT025713 | MPGGPCGSDCSCQN-CRCCSAA------AKCSANCKCDTCRCCQRSQ-TCRCGNCRCCSSAKC |
| SlinMT3Sb_JT027986 | MPGGPCGANCSCEK-CRCCSAA------AKCSAGCKCDVCRCCQRGQ-KCNCGDCRCCGAAKC |
| SmimMT3Sa_KFM81894 | MPGGPCGPNCTCRD-CRCSSSA------VKCNSGSKGETCRCSQNGQ-PCRCRDCKCRGGANR |
| SmimMT3Sb_JT036585 | MPGGPCGANCNCEK-CRCCSAA------AKCSAGCKCDVCHCCQRGQ-KCNCGDCRCCGAAKC |
| SocrMT3Sa_GIZU01029122 | MPGGPCGSNCTCEK-CRCCSAA------SKCSAGCRCDTCRCCQRGQ-KCQCGDCKCCGAAKC |
| SocrMT3Sb_GIZR01034889 | MPGGPCGANCTCEK-CRCCSAA--GSSGGRCGPNCRCDVCRCCNRGQ-NCRCGDCRCCSASKC |
| SoriMT3S_IBGW01035513 | MPGGPCGPNCNCEK-CRCCSAA------SKCTSNCKCDVCRCCQRGQ-RCNCGDCKCCGEAKC |
| SperMT3S_IBIS01040974 | MPCGTCGPNCTCDK-CRCCGAA----ASSKCSANCNCGDCKCCGRTSANCRCGDCKCCGAAKC |
| StenMT3S_JT042034 | MPGGPCGANCNCEK-CRCCSAA------AKCSAGCKCDVCRCCQRGQ-KCNCGDCRCCGAAKC |
| TangMT3S_IBHB01011973 | MPGGPCGSNCTCEK-CRCCTAA------SKCPASCKCDTCRCCQRSQ-KCHCGDCKCCGAAKC |
| TaquMT3Sa_IASZ01021367 | MPCGTCGPNCTCEK-CRCCGAA----AAPKCSANCNCGDCKCCGRTSANCRCGDCKCCGAAKC |
| TaquMT3Sb_IBZV01027233 | MPGGPCGPNCSCEK-CRCCSAA--ASSGGRCGPNCRCDVCRCCNRGQ-NCRCGDCKCCGASKC |
| TfenMT3S_IAVL01031865 | MPGGPCGANCTCEK-CRCCSAA------SKCGANCKCDNCRCCSRGQ-NCRCGDCKCCGAAKC |
| TlabMT3S_ICCK01008992 | MPGGPCGSNCSCEK-CRCCSAA-ASASASKCGPNCRCDVCRCCNRGQ-NCRCGDCKCCGASKC |
| TokiMT3S_ICDD01032490 | MPGGPCGANCSCEK-CRCCSAA-ASASASKCGPNCRCDVCRCCNRGQ-NCRCGDCKCCGASKC |
| TperMT3Sb_GGOF01118223 | MPGGPCGANCNCER-CRCCSAA------SKCSSGCKCDTCRCCQRNQ-KCQCGDCKCCGAAKC |
| TperMT3Sa_GGOF01037635 | MPGGPCGANCNCEK-CRCCSAA------SKCTANCKCDTCRCCQRNQ-QCNCGDCKCCSSAKC |
| TrimMT3S_IAWZ01037322 | MPGGPCGSNCSCEK-CRCCSAA-ASASASRCGPNCKCDVCRCCNRGQ-NCRCGDCKCCGASKC |
| TterMT3S_IBIR01028004 | MPGGPCGSNCTCEK-CRCCSAA------SKCSDGCKCDNCKCCQRGQAKCHCGDCKCCGAAKC |
| TwaiMT3S_GITR01027595 | MA-GPCGANCTCEK-CRCCSAA------SKCTSGCKCESCSCCQRSQ-KCNCGDCKCCGGAQC |
| UdivMT3S_GJZJ01032433 | MPGGPCGANCTCEK-CRCCGAA------AKCSASCKCETCRCCQRGQ-NCRCGDCKCCGAAKC |
| XtraMT3S_IAPE01005889 | MPCGTCGPNCTCEK-CRCCGAA----AAPKCSANCNCGDCKCCGRTSANCRCGDCKCCGAAKC |
| **Mesothelae** | |
| HhelMT3Sa_IAMB01031069 | MP-GPCGQNCTCER-CRCCAAA------TKCSSGCRCDTCRCCTRGQ-KCTCGDCKCCGAAKC |
| HhelMT3Sb_IAMB01033754 | MP-GPCGQNCTCER-CRCCSAA------TKCSSGCRCDTCRCCTRGQ-KCNCGDCKCCGAAKC |
| LmurMT3S_ICAH01016204 | MP-GPCGQNCTCER-CRCCSAA------TKCSAGCRCDTCRCCTRGQ-KCNCGDCKCCGAAKC |
| RnisMT3Sa_IAIP01026854 | MP-GPCGPNCTCDK-CRCCTAA------TKCSEGCKCDTCRCCERGQ-KCQCGDCKCCGAAKC |
| RnisMT3Sb_IAQS01029875 | MP-GPCGQNCTCDR-CRCCSAA------TKCSSGCRCDTCRCCTRSQ-KCNCGDCKCCGAAKC |
| RnisMT3Sc_IAQS01006286 | MP-GPCGQNCTCER-CRCCSAA------TKCSAGCRCDTCRCCTRGQ-KCNCGNCKCCGAAKC |
| **Mygalomorphae** | |
| ArorMT3Sa_IBPI01004899 | MP-GPCGQNCTFDR-CRCLAAA------TGCSGGCRCENCRCCTRGQ-KCNCVDCKCCGAAKC |
| ArorMT3Sb_IBPI01005671 | MP-GPCGQNCTCER-CRCCTAA------TRCSSGCRCETCRCCTRGQ-KCNCGDCKCCGAAKC |
| CsigMT3S_IBXS01007514 | MP-GPSGPNCSCER-CSCCAAA------TRCSSGCRCETCRCCSRGQ-RCNCGDCKCCGAAKC |
| CfraMT3Sa_IAVG01019714 | MP-GPCGQNCNCQQ-CRCCTAA------TKCSDGCRCETCRCCTRSQ-RCNCGDCKCCGAAKC |
| CfraMT3Sb_IAVG01034978 | MP-GPCGQNCNCER-CRCCTAA------AKCSEGCRCETCRCCTRSQ-KCNCGDCRCCGAAKC |
| MgigMT3S_IAEV010076969 | MP-GPCGQNCNCGQGCRCCTAA------VRCTDGCRCETCRCCSRSQ-KCNCGDCRCCGAAKC |
| MgigMT3Sb_IAEV01004369 | MP-GPCGQNCNCER-CRCCTAA------SKCTADCRCETCRCCSRSQ-KCECGDCKCCGAAKC |
| RatrMT3Sa_GIXJ01115636 | MP-GPCGQNCNCER-CRCCTAA------TRCSDGCRCETCRCCTRSQ-KCNCGDCRCCGAAKC |
| RatrMT3Sb_GIXJ01115623 | MP-GPCGPNCNCGDQCRCCTAA------TKCSAGCRCETCRCCTRSQ-RCNCGDCRCCGAAKC |

C xn [CxC] xn [C x CC] · C xn [CxC] xn [C x CC] · [CxC] xn [C x CC] xn C

### Figure S3B

#### B) Araneae MT3L

```
                                   1        10        20        30        40        50
Araneomorphae
AforMT3L_IAUW01044254              MDK--CGPNCKCTKCRCCDA-------AKSATSS-------------CCHA--
AipsMT3L_IBTU01015883              MDK--CGANCKCTKCRCCDA-------AKESAPA-------------CCQAAG
AjovMT3L_IATN01033941              MEKK-CGPNCRCAKCRCCDA-------AKQTAPP-------------CCTASA
AnaeMT3L_IBOW01008935              MERT-CGPNCKCTKCRCCDA-------AKSG----------------------
ApotMT3L_IBWF01014765              MERT-CGPNCKCTKCRCCDA-------AKSG----------------------
AspMT3L_IBEZ01030571               MDK--CGPNC---KCRCCDA-------AKSATPG-------------CCQA--
AtanMT3L_IBAW01038778              MEKT-CGPNCTCTKCRCCDS-------AKQAAPP-------------CCTAAA
BdecMT3L_IAQQ01008592              MDK--CGPNCKCTRCRCCDA-------AKSVTPTC------------CCQDSA
BtexMT3L_ICNG01037212              MERT-CGPNCKCTKCRCCDA-------AKSG----------------------
BtubMT3L_IBHZ01007440              MDK--CGPNCKCTKCRCCDA-------AKSATPT-------------CCQDAA
CbasMT3L_IBNI01016319              MDKR-CGPNCKCTKCRCCDG-------GKGA----------------------
CforMT3L_IBYP01009086              MDK--CGPNCKCTKCRCCDA-------AKSATPA-------------CCQTAA
CfulMT3L_IBOC01030735              MDK--CGPNCKCTKCRCCDA-------AKSVTPAC------------CCQDSA
CjapMT3L_IBPR01039541              MDNK-CGPNCKCTKCRCCDS-------TGGAAPC-------------CTGGAA
ClasMT3L_IAVB01025709              MDKK-CGPNCKCTKCRCCDS-------AKP-----------------CCTGAV
DcanMT3L_ICHF01001373              MERK-CSPNCKCTKCQCCPA------SVKQTS----------------------
DmarMT3L_GIWQ01151592              MEKAPCGSNCKCTKCRCCDG-------A-------------------QCGM--
DspMT3L_IBQW01022981               MDK--CGANCKCTKCRCCDA-------AKPATLS-------------CCQSAA
DsubMT3L_ICGF01021847              MDK--CGPNCKCTKCRCCDA-------AKSATPA-------------CCQA--
DtriMT3L_GGRN01035980              MDK--CGANCKCTKCRCCDA-------AKPATLS-------------CCQSAA
EkumMT3L_IANA01024013              MDR--CGPNCKCTKCRCCDA-------AKSATPA-------------CCQAAA
EtriMT3L_IBTQ01004588              MDK--CGPNCKCTKCRCCDA-------AKSATPA-------------CCQA--
FgraMT3L_IBPA01036519              MEKK-CGANCKCTKCRCCDS-------ARQAAPP-------------CCTAAA
HadaMT3L_IBMC01008950              MEKK-CGPNCKCTKCRCCNA-------ARSAAPS-------------CCTAAS
HmelMT3L_IAJG01015975              MDK--CGPNCKCTKCRCCDA-------AKSATPT-------------CCQDAA
HvenMT3L_IBFQ01012557              MERK-CGASCKCTKCQCCPE------SVKQAS----------------------
LcoeMT3L_IBIG01003521              MDK--CGPNCKCTKCRCCDA-------AKDSTPA-------------CCQAAA
LcorMT3L_IBDW01040765              MEKN-CGPNCKCTKCRCCNA-------AKG-----------------------
LevaMT3L_IBFM01000596              MDKS-CGPNCKCTKCRCCDA-------AKSASPA-------------CCAA--
LmaiMT3L_ICAK01042855              MEK--CGPNCKCTKCRCCDA-------AKSAT---------------CCQDAA
MbelMT3L_IAGN01039957              MDK--CGPNCKCTKCRCCDA-------AKSATPS-------------CCQDAA
MineMT3L_ICJC01003799              MEKK-CGPNCKCTKCRCCDS-------AKQAAPP-------------CCTAAA
MmusMT3L_GJIM01217943              MERK-CGPNCRCTKCRCCDA-------AKQSAPP-------------CCTAAA
MpseMT3L_ICBP01040719              MEKN-CGPNCKCTKCRCCNA-------AKG-----------------------
MshiMT3L_IAKA01013492              MDKR-CGANCKCTKCRCCDA------QKPKQAA---------------------
MspMT3L_IAGI01044335               MEKK-CGPNCKCTKCRCCDA-------AKTAP---------------CCTAAA
MyawMT3L_IBNG01023303              MERK-CGPNCRCTKCRCCDA-------AKQSAPP-------------CCTAAA
NminMT3L_IAIC01046021              MENK-CGPNCKCTKCRCCDA-------AKQATPS-------------CCAAAA
OhosMT3L_IBFD01027433              MDK--CGPNCKCTKCRCCDA-------AKSATPA-------------CCQA--
OlunMT3L_ICGC01023166              MER--CGPNCKCTKCRCCDA-------GKSGAA--------------------
OmatMT3L_IBFR01009325              MDK--CGPNCKCTKCRCCDA-------AKSASPC-------------CCQSAA
PameMT3L_GJIU01357226              MDK--CGANCKCTKCRCCDA-------VKESTPA-------------CCQAAA
PbicMT3L_ICDN01027384              MDK--CGPNCKCTKCRCCDA-------AKPATPA-------------CCQEAA
PbreMT3L_IAWA01009108              MDK--CGPNCKCTKCRCCDA-------AKSAKPS-------------CCQT--
PceyMT3L_IAYL01009101              MDK--CGPNCKCTKCRCCEA-------AKSATSC-------------CCEDAA
PclaMT3L_ICBG01041784              MEKK-CGPNCRCTKCRCCDA-------AKQAAPP-------------CCTAAA
PemaMT3L_IBPM01010691              MDS--CGPNCKCSKCRCCDA-------ATQSALT-------------CCCAAS
PkatMT3L_IAOS01005111              MDK--CGPNCKCTKCRCCDA-------AKSATTC-------------CCQDAA
PlamMT3L_IASI01033194              MEGKTCGSNCKCTKCRCCDA-------AKTG----------------------
PlamMT3L_IBMA01021694              MDK--CGPNCKCTKCRCCDA-------AKPATPA-------------CCQEAA
PnigMT3Lpart_GFUY01014770          -----------------------------------------------------
PpilMT3L_IBOK01002666              MEKN-CGPNCKCTKCRCCNA-------AKG-----------------------
PpseMT3L_GGRD01246346              MDK--CGANCKCTKCRCCDA-------AKESTPA-------------CCQAAA
PspMT3L_IBWH01021393               MEKK-CGSSCKCTKCQCCPA------SVKQAS----------------------
PspiMT3L_IBKY01007427              MERK-CGSNCKCTKCQCCPA------SVKQAS----------------------
PundMT3L_IBZD01006492              MDK--CGPNCKCTKCRCCDA-------AKSATPT-------------CCQAAA
RacuMT3L_IAYK01009383              MDK--CGPNCKCTKCRCCDA-------AKSATPA-------------CCQAAA
RhatrMT3L_IAUA01039453             MERK-CGPNCRCVKCRCCDA-------SKQAAPP-------------CCTAAA
SannMT3La_IBPH01038578             MEK--CGANCKCTKCRCCGA-------AKSAAPS-------------CCTAAS
SannMT3Lb_IAOG01004562             MET--CGANCRCTKCRCCDA-------GKAAAPS-------------CCTAAA
ScamMT3L_ICHM01000361              MDK--CGPNCKCTKCRCCDA-------AKTATPS-------------CCQAAA
SchiMT3L_ICKA01016492              MDK--CGPNCKCTKCRCCDA-------AKSATPA-------------CCQA--
ScyaMT3L_IBGG01005528              MERR-CGANCKCTKCRCCEA-------GKTAAAR---------------QA--
SduMT3L_XP_035211109               METK-CGVGCQCNQCKCCDV-------SKTRDS--------------CCDKT-
SforMT3L_ICBI01018703              MERK-CGSNCKCTKCQCCPA------SVKQAS----------------------
SlinMT3L_JT033686                  METK-CGAECQCKECQCCDA-------AKTGDS--------------CCDKT-
SmimMT3L_JT040526                  METK-CGAGCQCKQCQCCDA-------AKAGDS--------------CCDKT-
SocrMT3L_GIZR01017868              MDK--CGANCKCTKCRCCDA-------AKESTPA-------------CCQAAA
SoriMT3L_IBGW01024166              MDK--CGANCKCTKCRCCDA-------AKGAAPS-------------CCPSAA
SrubMT3L_IBXR01041414              MDK--CGPNCKCTKCRCCDA-------AKTATPS-------------CCQAAA
TangMT3L_IBHB01010593              MDK--CGPNCKCTKCRCCDA-------AKSPG---------------CCQA--
TaquMT3L_IASZ01035477              MDK--CGANCKCTKCRCCDA-------AKDTTPA-------------CCQAAA
TbunMT3L_IALY01018183              MDS--CGPNCKCSKCRCCDA-------AKQSAPS-------------CCCGAS
TfenMT3L_IAVL01011395              MDR--CGPNCKCSKCRCCDA-------ATQSAPS-------------CCCGAS
TindMT3L_IBFA01024184              MDK--CGPNCKCTKCRCCDA-------AKSATPA-------------CCQV--
TlabMT3L_IAWV01002859              MDK--CGPNCKCTKCRCCDA-------AKSATPA-------------CCQAAA
TokiMT3L_ICDD01018390              MDK--CGPNCKCTKCRCCDA-------AKSATPA-------------CCQAAA
TperMT3L_GGOF01029183              MDK--CGTNCKCTKCRCCDA-------GKSAGPA-------------CCQAAA
TrimMT3La_IAVJ01015074             MDK--CGPNCKCTKCRCCDA-------AKS--PA-------------CCQA--
TrimMT3Lb_IAWZ01003419             MEKN-CGPNCKCTKCRCCDA-------A--------------------------
TterMT3La_IBIR01022592             MDK--CGPNCKCTKCRCCDS-------AKQASP--------------CCTGG-
TterMT3Lb_IBIR01015641             MDK--CGANCKCTKCRCCDA-------AKESTPA-------------CCQAAA
VspMT3L_ICOO01039941               MERK-CGPNCKCTKCRCCDS-------GKQAAPP-------------CCTAAA
XinsMT3L_IAFS01028859              MDK--CGPNCKCTKCRCCDA-------AKSVTPA-------------CCQASA
ZasiMT3L_IAGO01012009              MEK--CGPNCTCANCRCCEA-------A-GAAPS-------------CCTAA-
Mesothelae
HhelMT3L_IAMB01018332              MADK-CGPNCKCQKCRCCDA------SVKAQAAG-------------CCG---
LmurMT3L_ICAH01020478              MAGK-CGPNCKCEKCRCCDASGPASASAKAQGAG-------------CCG---
RnisMT3L_IAIP01021207              MAGK-CGPNCKCEKCRCCDASAPASASAKAQAAG-------------CCG---
Mygalomorphae
ArorMT3L_IBPI01012923              MESK-CGPNCECTECGCCAALSSATKDCCGSQCTCEAGKCECCDTTKCCG---
CsigMT3L_IBXS01029286              MEGK-CGSQCQCTECRCCGALSSASKGCCGDQCCCEAGKCQCCDTAQSCG---
```

C $x_n$ [CxC] $x_n$ [C x CC]    CC $x_n$ [CxC] $x_n$ [C x CC]    CC $x_n$....

| | |
|---|---|
| | 60 · 70 · 80 · 90 · 100 |
| Araneomorphae | |
| AforMT3L_IAUW01044254 | --KKTRCECDDCQCCKSAG---------CGK-SCQC-NCKCCSGSK--Q--KE |
| AipsMT3L_IBTU01015883 | --KKATCGCDDCTCCSSSGCCE-KTA--SGK-GCQC-NCKCCSPAAR-Q--KS |
| AjovMT3L_IATN01033941 | A-KKAACSCDDCTCCQASSCGESQKT--CGK-GCQCGDCKCCSSSAC-QTGKS |
| AnaeMT3L_IBOW01008935 | --KKADCGCEDCKCCSSSSC-GSGKS--CGS-GCTCGDCKCCSSSAC--KTAS |
| ApotMT3L_IBWF01014765 | --KKADCGCEDCKCCSSSSC-GSGKS--CGN-GCTCGDCKCCSSSAC--KTAS |
| AspMT3L_IBEZ01030571 | --KKTRCGCDDCLCCKSSGC-GAQKSD-CGK-GCQC-NCKCCSGPS--Q--KD |
| AtanMT3L_IBAW01038778 | ASKKADCGCDNCTCCQSSSC-GSQKS--CGK-GCQCGDCKCCSASPC-QTNKS |
| BdecMT3L_IAQQ01008592 | --KKANCVCDDCQCCKSSSC-ETQKSD-CGK-GCQC-SCKCCSGGT--Q--KD |
| BtexMT3L_ICNG01037212 | --KKADCGCEDCKCCSSSSC-GSGKS--CGS-GCTCGDCKCCSSSAC--KTAS |
| BtubMT3L_IBHZ01007440 | --KKTNCECDDCQCCKSSDC-GAQKSG-CGK-GCQC-SCKCCSGAT--Q--KD |
| CbasMT3L_IBNI01016319 | --AKADCGCEECRCCSSSGC-GSQKS--CGK-GCQCGDCKCCSSSAC-Q--TS |
| CforMT3L_IBYP01009086 | --KKTNCGCDDCQCCKSSGC-GTQKSD-CGE-GCQC-NCKCCSGAT--Q--KD |
| CfulMT3L_IBOC01030735 | --KKANCACDDCQCCKSSSC-ATQKSD-CGK-GCQC-NCKCCSGGT--Q--KD |
| CjapMT3L_IBPR01039541 | G-KKADCGCDDCRCCSSAPCSATQKS--CGK-GCQCGDCKCCSSSPC-QTAKS |
| ClasMT3L_IAVB01025709 | --GKADCGCEDCKCCSNSSC-KSQKS--CGQ-GCQCGDCKCCSSSPC-QAAKS |
| DcanMT3L_ICHF01001373 | ----KDCGCEECRCCKTSAC-ATGKT--CGK-GCQCGDCKCCSSSGC-QTSRA |
| DmarMT3L_GIWQ01151592 | --KAGNCGCAECQCCSSSACQSGGKC--CGK-GCRCEDCKCCSAASG-Q-TKA |
| DspMT3L_IBQW01022981 | --TKAACGCDDCLCCNSSGC-GAQKA--SDK-GCQC-ECKCCSSSN--Q--KN |
| DsubMT3L_ICGF01021847 | --KQTKCGCDDCLCCKSSGC-GAQKSD-CGK-GCQC-NCKCCSSGA--Q--KD |
| DtriMT3L_GGRN01035980 | --TKAACGCDDCLCCNSSGC-GAQKA--SDK-GCQC-ECKCCSSSS--Q--KN |
| EkumMT3L_IANA01024013 | --KKTDCGCDDCLCCKSSSC-GAQKSD-CGK-GCQC-SCKCCAGAA--Q--KD |
| EtriMT3L_IBTQ01004588 | --KKTSCKCDDCLCC--------TQKSD-CGK-ECQC-SCKCCSGPA--Q--KD |
| FgraMT3L_IBPA01036519 | A-KKAGCSCDDCTCCKSSSC-TSQKT--CGK-GCKCGDCQCCSSSGC-Q-GDK |
| HadaMT3L_IBMC01008950 | A-KTADCGCDDCTCCQKASCCGSQKT--CGK-GCDCGDCKCCSSSSC-QAGKS |
| HmelMT3L_IAJG01015975 | --KKTDCRCEDCQCCKSSSC-GTQKSD-CGK-GCQC-NCKCCSGDT--Q--KD |
| HvenMT3L_IBFQ01012557 | ----KDCGCEECRCCKSSPC-ASGKT--CGK-GCQCGDCKCCSSSAC--KTKS |
| LcoeMT3L_IBIG01003521 | --KKTSCGCDDCSCCSSSGCCE-QKA--SGK-GCQC-SCKCCSPAAS-Q--KN |
| LcorMT3L_IBDW01040765 | ----KDCGCETCRCCDASSC-KSAKS--CGS-GCDCRDCKCCSATSCCQ--TS |
| LevaMT3L_IBFM01000596 | --KKTDCGCDDCQCCKSSSC-GTQKSD-CGK-ECQC-NCKCCSGAA--Q--RD |
| LmaiMT3L_ICAK01042855 | --KKTNCGCDDCQCCKSAGS-GAQKSD-CGK-GCQC-NCKCCSGGK--Q--KD |
| MbelMT3L_IAGN01039957 | --KKANCGCDDCQCCKSSSC-GTQKSD-CGE-GCQC-SCKCCAGAT--Q--KD |
| MineMT3L_ICJC01003799 | A-KKADCGCNDCSCCQNSSC--------CGSKGCQCSDCKCCSSSSC-QTSKS |
| MmusMT3L_GJIM01217943 | ASKKADCGCDDCTCCQKSACCGSEKT--CGK-GCQCGDCRCCSSSSC-QAGKS |
| MpseMT3L_ICBP01040719 | ----KDCGCETCRCCDASSC-KSAKS--CGS-GCDCRDCKCCSATSCCQ--TS |
| MshiMT3L_IAKA01013492 | ---GTDCGCEECRCCKSAPC-SSGKT--CGK-GCQCGDCKCCSSSAC-Q--TS |
| MspMT3L_IAGI01044335 | A-KKADCGCDDCHCCKSSSC-ASQKT--CGK-GCQCGDCKCCSSSAC-QTSKS |
| MyawMT3L_IBNG01023303 | ASKKAACGCDDCTCCQKSACCGSEKT--CGK-GCQCGDCKCCSASSC-QAGKS |
| NminMT3L_IAIC01046021 | A-KKADCGCDDCTCCQKSSC-GSQKS--CGK-GCQCSDCKCCSSAAC--HASK |
| OhosMT3L_IBFD01027433 | --KKTSCKCDDCLCC--------TQKSD-CGK-ECQC-SCKCCSVPA--Q--KD |
| OlunMT3L_ICGC01023166 | ----SGCGCKDCSCCGSSSC-GSQKT--CGK-GCQCKDCKCCSSSGC-QTGKS |
| OmatMT3L_IBFR01009325 | --KKANCRCDDCQCCKSSSC-GTQKSD-CGK-GCQC-NCKCCSGGT--Q--KD |
| PameMT3L_GJIU01357226 | --RKTTCGCDDCSCCSSSGCCE-PKA--VGK-GCQC-NCKCCSPASS-Q--KN |
| PbicMT3L_ICDN01027384 | A-KKAACGCDDCQCCSSSGC-GTQKT--CGK-GCQCGDCKCCSAAAS-Q--KK |
| PbreMT3L_IAWA01009108 | --AKKSCECDDCRCCNSSSCQ-------TGK-ECQC-NCKCCSASSE-Q--KD |
| PceyMT3L_IAYL01009101 | --KKANCRCDDCQCCKSSDC-GTQKSD-CGE-GCQC-SCKCCSGAM--Q--KD |
| PclaMT3L_ICBG01041784 | A-KKADCGCDDCTCCQKSSCCGSEKA--CGK-GCQCGDCRCCSSSAC-QTGKS |
| PemaMT3L_IBPM01010691 | --GKKSCSCTDCTCCKSSSCCEKQKS--CGE-G-KCDNCKCCEASPC-QTSKS |
| PkatMT3L_IAOS01005111 | --KKANCRCDDCQCCKSSDC-ETQKSD-CGK-GCQC-NCKCCS-GAT-Q--KD |
| PlamMT3L_IASI01033194 | ----KECGCEECRCCKSASC-GSQQST-CGK-GCTCGDCKCCSSADC-Q--QK |
| PlamMT3L_IBMA01021694 | A-KKAACGCDDCQCCSSSGC-GTQKT--CGK-GCQCGDCKCCSAATS-Q--KK |
| PnigMT3Lpart_GFUY01014770 | ------CDCDDCTCCNSSGC-GAQKT--CGK-GCQCSDCKCCSSASC---HKA |
| PpilMT3L_IBOK01002666 | ----KDCGCETCRCCDASSC-KSAKS--CGS-GCDCRDCKCCSATSCCQ--TS |
| PpseMT3L_GGRD01246346 | --KKTSCGCDDCSCCSSSGCCE-QTA--SGK-GCQC-SCKCCSPASS-Q--KK |
| PspMT3L_IBWH01021393 | ----KDCGCEECRCCKSSPC-ASGKT--CGK-GCQCGDCKCCSSSAC-Q--TS |
| PspiMT3L_IBKY01007427 | ----KECGCEECRCCKSSPC-ASGKT--CGK-GCQCGDCKCCSSSAC---KTS |
| PundMT3L_IBZD01006492 | --MKVNCRCDDCQCCKSS---GTQKSD-CGK-GCQC-SCKCCSGAT--Q--KD |
| RacuMT3L_IAYK01009383 | --KKADCGCDDCQCCKSSGC-GTQKSD-CGK-GCQC-SCKCCSGAA--Q--KE |
| RhatrMT3L_IAUA01039453 | A-KTADCGCDNCTCCQKSSCCGSEKA--CGK-GCVCGDCKCCSSSPC-QTSKS |
| SannMT3La_IBPH01038578 | A-RKADCGCDDCTCCQKSPCCASQKT--CGK-GCDCGDCKCCSSSAC-QKGKS |
| SannMT3Lb_IAOG01004562 | S-KKADCSCDDCTCCQKSSC-GSQKS--CGK-GCDCSDCKCCSSSACRTAGKT |
| ScamMT3L_ICHM01000361 | --KKSTCGCDDCSCCKSSGC-GSGKTD-CGK-GCQC-NCKCCSSASK-Q--KD |
| SchiMT3L_ICKA01016492 | --KKTSCGCDDCQCCKSSGC-GTQKSD-CGK-GCQC-NCKCCSGTS--Q--KD |
| ScyaMT3L_IBGG01005528 | ---GKDCGCDECRCCKDSSC-GTGKS--CGN-GCECGDCKCCSSAAC-Q--TS |
| SduMT3L_XP_035211109 | --GSKDCCCINSSCCKESGC-NIAKQG-TG--DCNCADCKCCSTATC-Q--TK |
| SforMT3L_ICBI01018703 | ----KDCGCEECRCCKSSPC-ASGKS--CGK-GCQCGDCKCCSSSAC-Q--TS |
| SlinMT3L_JT033686 | --GSKDCCCTNSSCCKESGC-NVSKQG-TG--NCNCTDCKCCSTATC-Q--MK |
| SmimMT3L_JT040526 | --GSKDCCCINSPCCKASGC-NISKEG-TG--NCNCTDCKCCSTATC-Q--TK |
| SocrMT3L_GIZR01017868 | --KKTSCGCDDCSCCSSSGCCE-QKA--SGK-GCQC-SCKCCSPASR-Q--KN |
| SoriMT3L_IBGW01024166 | --KKSNCGCDDCRCCSSSGC-EAQKA--SGK-GCQC-ECKCCSS----Q--RN |
| SrubMT3L_IBXR01041414 | --KKSTCGCDDCQCCKSSSC-GSQKTD-CAK-GCQC-NCKCCSPASK-Q--KD |
| TangMT3L_IBHB01010593 | --AKTKCGCDDCQCCKS----GAQKSD-CSK-GCQC-NCKCCSK----Q--KD |
| TaquMT3L_IASZ01035477 | --KKTTCGCDDCSCCSSSGCC-EQKT--SGK-GCQC-SCKCCSS----Q--KN |
| TbunMT3L_IALY01018183 | --GKKSCDCTDCTCCKSSSCCEKEKS--CGE-G-KCDNCKCCAASPC-QTSNS |
| TfenMT3L_IAVL01011395 | --GKKSCDCTDCACCKSSSCCEKQKS--CGE-G-KCDNCKCCSASAC-QTSKS |
| TindMT3L_IBFA01024184 | --KKTKCGCDDCQCCKTSGA--SD----CGK-GCQC-NCKCCSGAT--Q--KD |
| TlabMT3L_IAWV01002859 | --QKTDCGCDDCQCCKSSSC-GTQKAD-CGK-GCQC-SCKCCSA----Q--KD |
| TokiMT3L_ICDD01018390 | --QKTDCGCEDCQCCKS--C-GTQKAD-CGE-GCPC-SCKCCST----Q--KD |
| TperMT3L_GGOF01029183 | --KKAACKCDDCRCCSSAAC-GKQKT--CGK-GCQCSDCKCCSSSSC-Q--KD |
| TrimMT3La_IAVJ01015074 | --AKKKCGCDDCQCCKS----GAQKVD-CGK-GCEC-NCKCCST----Q--KD |
| TrimMT3Lb_IAWZ01003419 | --KGKDCGCENCRCCDASSC-KSGKS--CGS-GCDCRDCKCCSASSCCQ--TS |
| TterMT3La_IBIR01022592 | --KKADCGCDDCRCCSSSPC-ATQKS--CGK-GCQCGDCKCCSSSPC-QSAKS |
| TterMT3Lb_IBIR01015641 | --KKTTCGCDDCSCCSSSGCCE-QKA--SGK-GCQC-SCKCCSPASS-Q--KN |
| VspMT3L_ICOO01039941 | A-KKAACGCDDCTCCKSSSC-GAQKS--CGK-GCQCGDCKCCSSSAC-QTSKG |
| XinsMT3L_IAFS01028859 | --KKTNCGCDDCQCCKSSSC-GTQKSD-CGK-GCQC-NCKCCSGGT--Q--KD |
| ZasiMT3L_IAGO01012009 | ----SKCSCADCSCCRSAPCCASKKS--CGK-GCQCGDCKCCSSSAC-QTGKS |
| Mesothelae | |
| HhelMT3L_IAMB01018332 | ---GKSCTCEKCQCCSSAGCGAAAKS--CGS-NCTCDDCRCCSEAQCAQ--K- |
| LmurMT3L_ICAH01020478 | ---GSSCSCAQCQCCTSAGCGAATAS--CGP-SCTCDACKCCSEAQCGQ--K- |
| RnisMT3L_IAIP01021207 | ---GGSCTCERCQCCSSAGCGAATKG--CGP-NCTCDDCRCCSEAQCAQ--K- |
| Mygalomorphae | |
| ArorMT3L_IBPI01012923 | --EGVQCSCEDCQCCSSAGC--GKKQ--SGP-CCACDDCNCCSSAAC-IS--S |
| CsigMT3L_IBXS01029286 | --GAEKCACSDCQCCSSSGCHKQEAAARCGP-SCDCDDCNCCGTAAC-QT--S |

...[CxC]$x_n$[C x CC] $x_n$ C    C $x_n$ [CxC] $x_n$ [C x CC] $x_n$ C

110 120 130 140

| | |
|---|---|
| **Araneomorphae** | |
| AforMT3L_IAUW01044254 | G--CGSNCQC-DVCTCCSAAKDSGACHCGDCKCCSAASC |
| AipsMT3L_IBTU01015883 | T--CGSSCKC-DVCSCCSAAKGSGDCHCGNCKCCSTACC |
| AjovMT3L_IATN01033941 | R--CGSSCRC-DVCTCCSAAKGSGDCHCGDCRCCSAASC |
| AnaeMT3L_IBOW01008935 | S-CCGTSCKC-DVCTCCSAAKTGGNCHCGDCKCCSTATC |
| ApotMT3L_IBWF01014765 | S-CCGTSCKC-DVCTCCSAAKTGGNCHCGDCKCCSTATC |
| AspMT3L_IBEZ01030571 | R--CGSNCQC-DVCTCCSATKDSVTCHCGNCKCCSAATC |
| AtanMT3L_IBAW01038778 | S--CGSNCKC-DVCTCCSAAKGGGVCHCGNCKCCPSSGC |
| BdecMT3L_IAQQ01008592 | S--CGSNCKC-DVCTCCSAAKDSGACHCGDCRCCSAASC |
| BtexMT3L_ICNG01037212 | S-CCGTSCKC-DVCTCCSAAKTGGDCHCGDCKCCSTAAC |
| BtubMT3L_IBHZ01007440 | R--CGSNCQC-DVCTCCSAAKDSGACHCGDCRCCSAASC |
| CbasMT3L_IBNI01016319 | A-CCGSNCKC-DVCTCCSAAKDGGDCHCGDCKCCSQAKC |
| CforMT3L_IBYP01009086 | S--CGSNCKC-DVCTCCSAAKDSGACHCGDCRCCSAASC |
| CfulMT3L_IBOC01030735 | S--CGSNCKC-DVCTCCSAAKDSGACHCGDCRCCSAASC |
| CjapMT3L_IBPR01039541 | S--CGSSCKC-DVCTCCSAAKGGGDCHCGDCKCCSTAAC |
| ClasMT3L_IAVB01025709 | S--CGSSCKC-DVCTCCSAAKDGGACHCGDCKCCTAAVC |
| DcanMT3L_ICHF01001373 | DKSCGASCKC-DVCTCCSAATGGGICHCGDCKCCSTAVC |
| DmarMT3L_GIWQ01151592 | S-CCCTNCKC-DVCSCCSAAKGSGTCPCGDCKCCPTAPC |
| DspMT3L_IBQW01022981 | R--CGSNCKC-DVCTCCSSATGSGNCHCGDCKCCSAAVC |
| DsubMT3L_ICGF01021847 | S--CGSNCKC-DVCTCCSATKDSGACHCGNCKCCSAASC |
| DtriMT3L_GGRN01035980 | R--CGSNCKC-DVCTCCSSATGSGNCHCGDCKCCFAAVC |
| EkumMT3L_IANA01024013 | S--CGSNCKC-DVCTCCSAATDSGACHCGDCRCCSAATC |
| EtriMT3L_IBTQ01004588 | S--CGSNCKC-DVCTCCSDAKESGASHCGNCRCCSAASC |
| FgraMT3L_IBPA01036519 | S-CCGSGCKC-DVCTCCSAAKGGGDCHCGDCRCCPSATC |
| HadaMT3L_IBMC01008950 | --CCGSNCRC-DDCNCCSAAKGSGDCHCGNCRCCSAASC |
| HmelMT3L_IAJG01015975 | S--CGSNCKC-DVCTCCSAAKDSGACHCGDCRCCSAASC |
| HvenMT3L_IBFQ01012557 | ---CGTGCKC-DVCTCCSAATGGGNCHCGDCKCCSTAKC |
| LcoeMT3L_IBIG01003521 | T--CGSSCKC-DVCSCCSAAKGSGDCHCGNCKCCSTACC |
| LcorMT3L_IBDW01040765 | S-CCGSGCKC-DKCTCCAAAKGKGTCNCGDCKCCSTAKC |
| LevaMT3L_IBFM01000596 | S--CGSNCKC-DVCTCCSAAKDSGNCHCGDCRCCSAASC |
| LmaiMT3L_ICAK01042855 | G--CESNCKC-DRCTCCPVTKDSGVCHCGFCRCCDLASF |
| MbelMT3L_IAGN01039957 | S--CGSNCKC-DVCTCCSAAKDSGACHCGDCRCCSTASC |
| MineMT3L_ICJC01003799 | S--CGSSCKC-DVCTCCSAAKGSGDCHCGNCKCCSAASC |
| MmusMT3L_GJIM01217943 | S--CGSNCKC-DVCTCCSAAKSGGDCHCGDCRCCSAASC |
| MpseMT3L_ICBP01040719 | S-CCGSGCKC-DKCTCCAAAKGKGTCNCGDCKCCSTAKC |
| MshiMT3L_IAKA01013492 | S--CGTSCAC-DVCTCCSASKGDGDCHCGDCKCCSSAKC |
| MspMT3L_IAGI01044335 | --CCGSSCKC-DVCTCCSAAKDGGVCHCGNCRCCETAAC |
| MyawMT3L_IBNG01023303 | S--CGSNCKC-DVCTCCSAAKSGGDCHCGDCRCCSAASC |
| NminMT3L_IAIC01046021 | S-CCGSSCKC-DVCTCCSAAKGSGDCHCGDCRCCSAASC |
| OhosMT3L_IBFD01027433 | S--CGSNCKC-DVCTCCSAAKDSGASHCGSCRCCSAASC |
| OlunMT3L_ICGC01023166 | --CCGSGCKC-DVCTCCSAAQGGGACHCGDCKCCSSAKC |
| OmatMT3L_IBFR01009325 | S--CGSNCKC-DVCTCCSAAKDSGACHCGDCRCCSAASC |
| PameMT3L_GJIU01357226 | T--CGSSCKC-DVCSCCSAAKGSGDCHCGNCKCCSTACC |
| PbicMT3L_ICDN01027384 | T--CGSNCRC-DVCTCCSSAKGSGNCHCGDCKCCSAAVC |
| PbreMT3L_IAWA01009108 | S--CGSNCKC-DVCTCCSASKESGDCHCGDCKCCSGAAC |
| PceyMT3L_IAYL01009101 | S--CGSNCKC-DVCTCCSAAKDSGACHCGDCRCCSTASC |
| PclaMT3L_ICBG01041784 | S--CGSNCKC-DVCTCCSAAKSGGDCHCGDCRCCSAASC |
| PemaMT3L_IBPM01010691 | S--CGSSCKC-DVCTCCSAAKGGADCHCGNCKCCSAATC |
| PkatMT3L_IAOS01005111 | S--CGSNCKC-DVCTCCSAAKDSGACHCGDCRCCSAASC |
| PlamMT3L_IASI01033194 | S--CGSGCKC-DVCSCCTAATGGGVCHCGNCKCCPSSGC |
| PlamMT3L_IBMA01021694 | T--CGSNCRC-DVCTCCSSAKGSGNCHCGDCKCCSAAVC |
| PnigMT3Lpart_GFUY01014770 | S--CGSDCKC-DVCTCCSAAKSSGDCHCGNCRCCSAAVC |
| PpilMT3L_IBOK01002666 | S-CCGSGCKC-DKCTCCAAAKGKGTCNCGDCKCCSTAKC |
| PpseMT3L_GGRD01246346 | A--CGSSCKC-DVCTCCSAAKGSGDCHCGNCKCCSTACC |
| PspMT3L_IBWH01021393 | S--CGAGCKC-DVCTCCSAATGGGNCHCGDCKCCSAAVC |
| PspiMT3L_IBKY01007427 | S--CGTGCKC-DVCTCCSAATGGGNCHCGDCKCCSSAVC |
| PundMT3L_IBZD01006492 | S--CGSNCKC-DVCTCCSAAKDSGACHCGDCRCCSAASC |
| RacuMT3L_IAYK01009383 | S--CGSNCKC-DVCTCCSAAKDSGACHCGDCRCCSAASC |
| RhatrMT3L_IAUA01039453 | --CCGSNCKC-DVCTCCSAAKSGGDCQCGNCRCCSAASC |
| SannMT3La_IBPH01038578 | --CCGSGCKC-DDCSCCSAAKGSGDCHCGDCKCCSSASC |
| SannMT3Lb_IAOG01004562 | --CCGSNCKC-DVCTCCPAAKGTTCSH------------ |
| ScamMT3L_ICHM01000361 | S--CGSSCQC-DVCTCCSAAKESGSCHCGDCKCCSAAVC |
| SchiMT3L_ICKA01016492 | S--CGSNCKC-DVCTCCSAAKDSGSCHCGDCKCCSAASC |
| ScyaMT3L_IBGG01005528 | S--CGSSCPC-DVCTCCSASKGSGNCCCGDCKCCPTAKC |
| SduMT3L_XP_035211109 | S----PSCKC-DVCTCCQKS-GGTDCHCGKCKCCPSAKS |
| SforMT3L_ICBI01018703 | S--CGTGCKC-DVCTCCSAATGGGNCHCGDCKCCSAAVC |
| SlinMT3L_JT033686 | S----PSCKC-DVCTCCQKS-GGTDCHCGKCKCCPSAKS |
| SmimMT3L_JT040526 | S----PSCKC-DVCTCCQKS-GGTDCHCGKCKCCPSAKS |
| SocrMT3L_GIZR01017868 | T--CGSSCKC-DVCSCCSAAKGSGDCHCGNCKCCSTACC |
| SoriMT3L_IBGW01024166 | T--CGSNCKC-DVCTCCSSAKGSGDCHCGNCKCCSAAGC |
| SrubMT3L_IBXR01041414 | S--CGSNCQC-DVCTCCSAAKESGSCHCGDCRCCSAAAC |
| TangMT3L_IBHB01010593 | S--CGSNCKC-DVCTCCSAAKDSGACHCGDCRCCSAASC |
| TaquMT3L_IASZ01035477 | T--CGSSCKC-DVCSCCSAAKGSGDCHCGNCKCCSTACC |
| TbunMT3L_IALY01018183 | S--CGSSCKC-DVCTCCSAAKGGADCHCGNCKCCSAASC |
| TfenMT3L_IAVL01011395 | S--CGPSCKC-DVCTCCSAAKGEADCHCGNCKCCSAASC |
| TindMT3L_IBFA01024184 | S--CGSNCKC-DVCTCCSAAKDSGACHCGDCRCCSAASC |
| TlabMT3L_IAWV01002859 | S--CGSNCKC-DVCTCCSAAKDSGACHCGDCRCCSAASC |
| TokiMT3L_ICDD01018390 | S--CGSNCKC-DVCTCCSAAKDSGACHCGDCRCCSAASC |
| TperMT3L_GGOF01029183 | C--CGSNCQC-DDCRCCSAAKGSGDCKCGDCKCCSAATC |
| TrimMT3La_IAVJ01015074 | S--CGSNCKC-DVCTCCSAAKDSGACHCGDCRCCSAASC |
| TrimMT3Lb_IAWZ01003419 | S-CCGSGCKC-DKCSCCAAAKGKGTCNCGDCKCCSAAKC |
| TterMT3La_IBIR01022592 | S--CGSNCKC-DVCTCCSAAKGGGDCHCGDCKCCGAAVC |
| TterMT3Lb_IBIR01015641 | T--CGSSCKC-DVCSCCSAAKGSGDCHCGNCKCCSTACC |
| VspMT3L_ICOO01039941 | S--CGSSCQC-DVCTCCSAAKGGGDCHCGDCKCCSSAVC |
| XinsMT3L_IAFS01028859 | S--CGSNCKC-DVCTCCSAAKDSGACHCGDCRCCSAASC |
| ZasiMT3L_IAGO01012009 | --CCGSDCNC-TVCRCCSAAKSGGDCRCGDCKCCSSASC |
| **Mesothelae** | |
| HhelMT3L_IAMB01018332 | ---CGSGCTC-DVCRCCSASRGQGRCQCGDCRCCSSASC |
| LmurMT3L_ICAH01020478 | ---CGSGCTC-DVCRCCSASRGQGRCQCGDCRCCSSASC |
| RnisMT3L_IAIP01021207 | ---CGSGCTC-DVCRCCSASRGQGRCQCGDCRCCSSASC |
| **Mygalomorphae** | |
| ArorMT3L_IBPI01012923 | SR-CGSGCQC-DDCSCCPAAHGQGTCECGDCKCCSTASC |
| CsigMT3L_IBXS01029286 | SR-CGSGCQCAGDCSCCPAAHGQGACTCGDCKCCSSATC |

C $x_n$ [CxC] $x_n$ [C x CC] [CxC]$x_n$ [C x CC] $x_n$ C
