## Supplementary material for "The Evolutionary History of Metallothioneins in Horseshoe Crabs, Scorpions, Spiders and other Chelicerate Species": Figure S5

```
                      1        10        20        30        40
ARTHROPODA
Chelicerata MT1
AinsMT1      MPEI-------ECKC-KEGE-CIC-GEECKCEG-GCDCG-TTCGK-----
NgraMT1      MPDT-------ECKC-KDGE-CGC-GEECKCT--NCEC--TKCGPKKS--
NmolMT1      MPET-------ECKC-KDGE-CVC-GENCPCE--NCEC--KACSK-----
PfemMT1      MPET-------ECKC-KEGE-CGCVEQECKCE--NCGC--ATCGK-----
PlitMT1_     MPGP-------ECKC-TEGQ-CAC-GEKCECT--ACGC--AKCAK-----
ScheMT1      MPET-------ECKC-KDGE-CGC-GESCQCE--NCEC--DSCPAKI---
TorbMT1      MPET-------ECKC-KDGE-CGC-AEACTCE--NCEC--KSCNAK----
Myriapoda MT1
LforMT1S     MPDP-------CCQC--TGEDCKC-GDSCQCGP-DCKC---KCAK-----
GminMT1Sa    MP----------CGC-EEGT-CVC-TDKCPCGD-DCKC--PGCKKA----
GminMT1Sb    MP----------CGC-EGGK-CVC-KDNCNCGA-DCKC--VGCKK-----
MOLLUSCA MT β2/β1
AcraMT1      MSDP--------CNCADTGD-CKC-AADCDCKAKGCKCG-AGCK------
AspMT1       MSDP--------CKCAETGD-CKC-AADCDCTAKGCKCG-AGC-------
MfoMT1       MPDK--------CNCAQGGE-CQC-GSDCNCS--DCKCDPDKCKAAQN--
NcaMT1       MSDP--------CKCAETGD-CKC-AADCDCTAKGCKCG-AGCKSTQH--
NmegMT1      MSDP--------CNCAETGD-CKC-EAGCDCS--KCNCDPAKCKVK----
NspMT1       MSDP--------CNCAESGE-CQC-KEXCDCT--XCKCDPAKXKVKA---
NspMT2       MSDP--------CNCAETGD-CKC-EAGCDCS--KCNCDPAKCKVK----
LhyMT1       MPDP--------CTCSETGESCKC-DSPCTCT--DCKCACAKCKT-----
AenMT1       MSDP--------CNCATGGN-CTC-GESCDCSKSNCNCDPSKCNSKCSSK
GebMT1       MSDP--------CNCATTGD-CKC-DESCACT--DCKCDASKCKSKSAAH
CHORDATA MT β/α
BfloMT1      MPDPCNCAQSGTCSC--GGP-CQC-GDDCQCGD-GCKC--VGCKLHSNVT
BlanMT1      MPDPCNCAQSGACSC--NGL-CQC-GDDCQCGD-GCKC--VGCKLHGNVD
BbelMT1      MPDPCNCAQSGACSC-AGGT-CQC-GDDCRCGD-GCKC--VGCKLHGNV-
HsapMT1A     M-DP-------NCSCATGGS-CTC-TGSCKCK--ECKC--TSCKKS----
HsapMT2A     M-DP-------NCSCAAGDS-CTC-AGSCKCK--ECKC--TSCKKS----
HsapMT3      M-DPE------TCPCPSGGS-CTC-ADSCKCE--GCKC--TSCKKS----
HsapMT4      M-DPR------ECVCMSGGI-CMC-GDNCKCT--TCNC--KTYWKS----
```

[CxC] $x_n$ [CxC] $x_n$ [CxC] $x_n$ [CxC] $x_n$ C

```
         60        70        80        90        100       110
--KCKCSTLDACDCGVGCSKDKACEC---------ADECGCKK-----------------
--TCKCSELEACDCGVGCSKEVKCEC---------KDDCKCKQ-----------------
--KCKCKELEACDCGVGCSKEAKCEC---------KEDCKCKQ-----------------
--KCKCSTLDACDCGAGCAKDKPCEC---------AEECGCKK-----------------
--KCSCSELDQCACGAGCCKEKACAC----------EECDCRK-----------------
--KCKCTTLDDCGCGEGCSKAEACKC---------GDDCSCKK-----------------
--KCKCSESETCTCGVGCNKDTACKC---------EDACGCKGE----------------
                                                  ----------
--SCKCTG------SSGCV--EGCAC---------AEGCACKNKV---------------
--HCKCFEAKSCSCGANCTT-GDCSC---------PKNCSCFS-----------------
--HCKCVQAKSCTCGADCA--NGCAC---------AKDCTCFS-----------------
                                                  ----------
                                                  ----------
                                                  ----------
--FTVCKHN---------------------------------------------------
--KCKCT-DS-CACGKGCKSGEACKC---------GTDCSCR------------------
--NAKCCTDSACKCADGCKGGDGCKC---------QSDCGCCK-----------------
--AKKCC--GKCECTDGCKGADGCKC---------AAGCDCCK-----------------
--NAKCCTDSACKCADGCKGGDGXKC---------Q------------------------
--VCKCP--GTCKCGNDCTGPENCKC---------KPDCSCK------------------
GGVCNCPADG-CKCGGECTGPSSCSC---------GPSCSCK------------------
--ACKCT-DSSCQCGKGCTGAASCSC---------APDCSCK------------------
                                                  ----------
D-IVTCCVD--CK---GIG--KNCAC-----------GCSCCQPDTPAVAILTTPPAAHL
V-TLTCCGT--CT---GIG--KNCAC-----------GCSCCQPDVPAVTVLTTPPAAHL
--SVTCCAM--CT---GKV--KNCAC-----------GCSCCQPAAPAVVNMTTPPAAHL
--CCSCCPMS-CA---KCA--QGCICKGA------SEKCSCCA-----------------
--CCSCCPVG-CA---KCA--QGCICKGA------SDKCSCCA-----------------
--CCSCCPAE-CE---KCA--KDCVCKGGEAAEAEAEKCSCCQ-----------------
--CCPCCPPG-CA---KCA--RGCICKGG------SDKCSCCP-----------------
```

[CxC] $x_n$ [CxC] $x_n$ C $x_n$ [CxC] $x_n$ [CxC]
