## Supplementary material for "The Evolutionary History of Metallothioneins in Horseshoe Crabs, Scorpions, Spiders and other Chelicerate Species": Table S1

Table S1. Putative chelicerate MT sequences identified in this study.

| Clade | Species | MT name | Accession number* | Sequence |
|---|---|---|---|---|
| Pycnogonida class | | | | |
| | Anoplodactylus insignis | AinsMT1 | SRA | MPEIECKCKEGECICGEECKCEGGCDCGTTCGKKCKCSTLDACDCGVGCSKDKACECADECGCKK |
| | Nymphon gracile | NgraMT1 | GCID01034565 | MPDTECKCKDGECGCGEECKCTNCECTKCGPKKSTCKCSELEACDCGVGCSKEVKCECKDDCKCKQ |
| | Nymphon molleri | NmolMT1 | SRA | MPETECKCKDGECVCGENCPCENCECKACSKKCKCKELEACDCGVGCSKEAKCECKEDCKCKQ |
| | Phoxichilidium cf.femoratum | PfemMT1 | SRA | MPETECKCKEGECGCVEQECKCENCGCATCGKKCKCSTLDACDCGAGCAKDKPCECAEECGCKK |
| | Pycnogonum litorale | PlitMT1 | SRA | MPGPECKCTEGQCACGEKCECTACGCAKCAKKCSCSELDQCACGAGCCKEKACACEECDCRK |
| | Stylopallene cheilorhynchus | ScheMT1 | SRA | MPETECKCKDGECGCGESCQCENCECDSCPAKIKCKCTTLDDCGCGEGCSKAEACKCGDDCSCKK |
| | Tanystylum orbiculare | TorbMT1 | SRA | MPETECKCKDGECGCAEACTCENCECKSCNAKKCKCSESETCTCGVGCNKDTACKCEDACGCKGE |
| Merostomata class | | | | |
| | Carcinoscorpius rotundicauda | CrotMT1a | SRA | MSDLICKCGDPLCTCTKECTCGKEGGCKCTTCNKKDEXDXSCKCKDPXCTCSVKCTCGQKDGCKCTSCT<br>RKDDTEGSPCKCTEGEPCSCTSCTCSKKE |
| | | CrotMT1b | SRA | MTDAVCKCGDQMCTCIKECTCGKEDGCKCTTCNKKLETERSCQCKDPTCTXIEKCTCGKEDGCKCTTCSK<br>KEIAGQCKCTEGEACSCTSCTCRKTE |
| | Limulus polyphemus | LpolMT1a | XR_002609474+SRA | MSDSACKCGDSLCTCVKECTCGKLDGCKCATCKKKEETEKSCKCKDPTCTCDKECTCGKEDGCKCTTCN<br>KKDETEKSXCKCIEGEPCSCTSCTCSKKQ |
| | | LpolMT1b | XR_002609207 | MSDSACKCGDPTCTCGENCTCGKEDGCKCTTCNKKDETEKSRCKCTEGEPCSCTSCTCSKKE |
| | | LpolMT1c | XR_001292027 | MPDPACKCGDELCTCGEECTCGKEDGCKCTTCNKKGEAETEASPCKCSKGESCTCASCTCSKKE |
| | Tachypleus gigas | TgigMT1a | SRA | MSETVCKCGDPLCTCTHECTCSKEGGCKCTTCNKKDETATSCKCKDPTCTCSDKCTCGQKDGCKCTSCM<br>RKDDTEGSPCKCTEGEPCSCTSCTCNKKE |
| | | TgigMT1b | SRA | MTDVVCKCGDPMCTCIKECTCXKEDGCKCTTCNKKDETERSCKCKDPTCTCIEKCTCGKEDGCKCTTCNK<br>KXETEDRCKCTEGETCSCTSCTCSKTD |
| | | TgigMT1c | SRA | MSDPACTCGDEMCTCGEECTCGKEDGCKCTTCNKKCEKEGEASPCKCSEGEPCSCESCSCSKKE |
| | Tachypleus tridentatus | TtriMT1a | GGKL01028814 | MSDLPCKCGDSLCTCTQECTCGKEGGCKCTTCNKKDETDWSCKCKDPTCTCSDKCTCGQKDGCKCTSC<br>MKKDDTEGSPCKCTEGEPCSCTSCTCSKK |
| | | TtriMT1b | GGKL01032357+SRA | MSDPACMCGDEMCTCGEECTCGKEDGCKCTTCNKKGEKEGEASPCKCSEGEPCSCESCTCSKKE |

### Arachnida class

#### Amblypygi order

| | | | | |
|---|---|---|---|---|
| | Acanthophrynus coronatus | AcorMT1 | SRA | MPDPACKCGDSKCNCQGSCECTSGKCPCSKCKTATSCNCTGGGNCSCSSCACKK |
| | Charinus ioanniticus | CioaMT1 | SRA | MPDPACKCGDEKCNCKGSCECASGACPCSKCKKACNCTGGESCTCSSCACKK |
| | Charinus israelensis | CisrMT1 | SRA | MPDPACKCGDEKCNCKGSCECASGACPCSKCKKACNCTGGESCTCSSCTCKK |
| | Damon diadema | DdiaMT1 | GERU01028960 | MPDPACKCGDSKCTCQGSCECASGGCPCSKCKKSSACKCTGGGDCTCSACACKK |
| | Euphrynichus bacillifer | EbacMT1 | GDAJ01000060 | MPDPACKCGDSKCTCQGSCECSSGGCPCSKCKKTSACKCTGGGDCTCSSCACKK |
| | Phrynus marginemaculatus | PmarMT1 | SRA | MPDPACKCGDSKCNCQGSCECSSGKCPCSKCKTASSCKCTGGGDCTCSSCACKK |

#### Araneae order

| | | | | |
|---|---|---|---|---|
| Araneomorphae suborder | Agelenopsis naevia | AnaeMT2 | IBOW01043046 | MADVEKKSSCTEPRECCKDASCCDEGKGECHTGKEECKDTCHEKSCGCKEDCKCSENKCKC |
| | | AnaeMT3L | IBOW01008935 | MERTCGPNCKCTKCRCCDAAKSGKKADCGCEDCKCCSSSSCGSGKSCGSGCTCGDCKCCSSSACKTAS SCCGTSCKCDVCTCCSAAKTGGNCHCGDCKCCSTATC |
| | | AnaeMT3S | IBPA01019018 | MPGPCGSNCNCDKCRCCSAASKCGANCKCDNCRCCQRGGQKCECGDCKCCGAAKC |
| | Agelenopsis potteri | ApotMT2 | IBWF01015665 | MADVEKKSSCTEPRECCKDASCCDEGKGECHTGKEECKDTCHEKSCGCKEDCKCSENKCKC |
| | | ApotMT3L | IBWF01014765 | MERTCGPNCKCTKCRCCDAAKSGKKADCGCEDCKCCSSSSCGSGKSCGNGCTCGDCKCCSSSACKTA SSCCGTSCKCDVCTCCSAAKTGGNCHCGDCKCCSTATC |
| | | ApotMT3S | IBWF01018135 | MPGGPCGSNCTCEKCRCCSAASKCTASCKCDTCRCCQRNQKCECGDCKCCSASKC |
| | Alenatea fuscocolorata | AfusMT3S | IBZC01022604 | MPCGTCGPNCSCEKCRCCGAAAAPKCSANCQCGDCKCCGRTSANCRCGDCKCCGAAKC |
| | Amyciaea forticeps | AforMT3L | IAUW01044254 | MDKCGPNCKCTKCRCCDAAKSATSSCCHAKKTRCECDDCQCCKSAGCGKSCQCNCKCCSGSKQKEGC GSNCQCDVCTCCSAAKDSGACHCGDCKCCSAASC |
| | Anepsion depressum | AdepMT3S | ICFY01031203 | MPCGTCGPSCTCDKCRCCGASSGSKCGPNCKCADCKCCGKSSANCRCGDCKCCGAAKC |
| | Apricia jovialis | AjovMT3L | IATN01033941 | MEKKCGPNCRCAKCRCCDAAKQTAPPCCTASAAKKAACSCDDCTCCQASSCGESQKTCGKGCQCGDC KCCSSSACQTGKSRCGSSCRCDVCTCCSAAKGSGDCHCGDCRCCSAASC |
| | Arachnura logio | AlogMT2 | IBSH01016593 | MADVEKKPCCAEPQECCKDESCCQEGKGPCHTDKDGCADSCQKKLCGCKSGEDCKCTDGKCAC |

| | | | |
|---|---|---|---|
| | AlogMT3Sa | IBSH01031123 | MPCGTCGSSCTCDKCRCCGASSDSKCSANCQCVDCKCCGRSSATCRCGDCKCCSASKC |
| | AlogMT3Sb | IBSH01034144 | MPGGPCGSNCTCEKCRCCSAASKCTASCKCDTCRCCQRNQKCECGDCKCCSASKC |
| Arachnura melanura | AmelMT2 | IAPN01004235 | MADVEKKPCCDEPKECCKDESCCQEGKGPCHDEKDGCADSCQKKPCGCKAGEDCKCSEGKCAC |
| | AmelMT3S | IAPN01022659 | MPCGTCGSNCTCDKCRCCGAASDSKCSANCKCVDCKCCGRSSANCRCGDCKCCSAAKC |
| Araneus diadematus | AdiaMT2 | GERP01021930 | MADVEKKSCCAEPKECCKDESCCQEGKGPCHNEKEVCADSCQKKPCGCKAGEDCKCKDGKCAC |
| Araniella yaginumai | AyagMT3S | IAYO01029315 | MPCGTCGPSCTCEKCRCCGASAAPKCSANCQCGDCKCCGRTSANCRCGDCKCCGAAKC |
| Arctosa ipsa | AipsMT2 | IBYV01021947 | MADVEKTSCCTEPKECCKDESCCENGQGACHTGKEECKDSCHKKACGCKAGEECKCTDGKCGC |
| | AipsMT3L | IBTU01015883 | MDKCGANCKCTKCRCCDAAKESAPACCQAAGKKATCGCDDCTCCSSSGCCEKTASGKGCQCNCKCCS PAARQKSTCGSSCKCDVCSCCSAAKGSGDCHCGNCKCCSTACC |
| | AipsMT3S | IBYV01029187 | MPGGPCGANCTCEKCRCCSAASKCSSGCRCDTCRCCQRGQKCQCGDCKCCGAAKC |
| Argiope argentata | AargMT2 | GIWY01029574 | MADVEKKTCCAEPKECCKDESCCQDGKGTCHTEKEGCADSCQKKPCGCKAGEDCKCKDGKCAC |
| | AargMT3S | GIWY01109359 | MPGGPCGANCNCEKCRCCSAASKCTANCKCDTCRCCQRNQQCNCGDCKCCSSAKC |
| Argiope bruennichi | AbruMT2 | KAF8785867 | MADVEKKACCAEPKECCKDESCCQEGKGPCHTEKEGCADSCQKKPCGCKAGEDCKCKDGKCAC |
| Argyrodes nephilae | AnepMT3S | IBFL01003327 | MPCGTCGPNCTCDKCRCCGAAASSKCSANCNCGDCKCCGRTSANCRCGACKCCGAAKC |
| Argyroneta aquatica | AaquMT2 | GIWR01226499 | MADVEKTTSCCTDPKECCKDDACCEQGQGPCHTGKEECKDTCHKKACGCKAGEDCKCSGGKCGC |
| | AaquMT3S | GIWR01171385 | MPGGPCGSNCTCERCRCCSAATKCNSGCRCETCRCCQRNQKCECGDCKCCGAAKC |
| Asemonea tanikawai | AtanMT2 | IBAW01019844 | MADVEKTTCCTESKECCKDESCCENGQGACHTGKEECKDSCHKKPCGCKTGEECKCSDGKCGC |
| | AtanMT3L | IBAW01038778 | MEKTCGPNCTCTKCRCCDSAKQAAPPCCTAAAASKKADCGCDNCTCCQSSSCGSQKSCGKGCQCGDC KCCSASPCQTNKSSCGSNCKCDVCTCCSAAKGGGVCHCGNCKCCPSSGC |
| | AtanMT3S | IBAW01043010 | MPCGTCGPNCTCEKCRCCSAAAKCSANCKCDDCKCCGRGQNCRCGDCKCCGAAKC |
| Australomisidia sp. | AspMT3L | IBEZ01030571 | MDKCGPNCKCRCCDAAKSATPGCCQAKKTRCGCDDCLCCKSSGCGAQKSDCGKGCQCNCKCCSGPS QKDRCGSNCQCDVCTCCSATKDSVTCHCGNCKCCSAATC |
| Badumna longinqua | BlonMT3 | GIWL01016652 | MPGGPCGTNCTCDKCRCCSAATKCSSGCKCDTCRCCQRGQKCECGDCKCCGAAKC |
| Barronopsis texana | BtexMT3L | ICNG01037212 | MERTCGPNCKCTKCRCCDAAKSGKKADCGCEDCKCCSSSSCGSGKSCGSGCTCGDCKCCSSSACKTAS SCCGTSCKCDVCTCCSAAKTGGDCHCGDCKCCSTAAC |

| | | | |
|---|---|---|---|
| Bassaniana decorata | BdecMT3L | IAQQ01008592 | MDKCGPNCKCTRCRCCDAAKSVTPTCCCQDSAKKANCVCDDCQCCKSSSCETQKSDCGKGCQCSCKC CSGGTQKDSCGSNCKCDVCTCCSAAKDSGACHCGDCRCCSAASC |
| Bathyphantes gracilis | BgraMT3S | IAMX01033138 | MPCGTCGPNCTCEKCRCCGAAAAPKCSANCNCGDCKCCGRTSANCRCGDCKCCGAAKC |
| Boliscus tuberculatus | BtubMT3L | IBHZ01007440 | MDKCGPNCKCTKCRCCDAAKSATPTCCQDAAKKTNCECDDCQCCKSSDCGAQKSGCGKGCQCSCKC CSGATQKDRCGSNCQCDVCTCCSAAKDSGACHCGDCRCCSAASC |
| Caerostris darwini | CdarMT2 | GGTX01063157 | MADVEKPCCAEPKECCKDESCCDEGKGPCHTDGKEDCADSCHKKPCGCKAGEDCKCKDGKCAC |
| | CdarMT3S | GIY23593 | MPCGTCGPSCSCDKCRCCGAAASASAKCSANCKCDDCRCCGRSGANCRCGDCKCCSAAKC |
| Caerostris extrusa | CextMT3S | GIY05480 | MPCGTCGPSCSCDKCRCCGAAASASAKCSANCKCDDCRCCGRSGANCRCGDCKCCSAAKC |
| Camaricus formosus | CforMT2 | IBYP01014520 | MADVENKSCCAEPKECCKDESCCQEGKGPCHTEKEGCADSCQKKPCGCKAGEDCKCQDGKCAC |
| | CforMT3L | IBYP01009086 | MDKCGPNCKCTKCRCCDAAKSATPACCQTAAKKTNCGCDDCQCCKSSGCGTQKSDCGEGCQCNCKC CSGATQKDSCGSNCKCDVCTCCSAAKDSGACHCGDCRCCSAASC |
| | CforMT3S | IBYP01019334 | MPCGTCGPNCTCDKCRCCGAAASSKCSANCNCGDCKCCGRTSANCRCGDCKCCGAAKC |
| Cheiracanthium japonicum | CjapMT3L | IBPR01039541 | MDNKCGPNCKCTKCRCCDSTGGAAPCCTGGAAGKKADCGCDDCRCCSSAPCSATQKSCGKGCQCGD CKCCSSSPCQTAKSSCGSSCKCDVCTCCSAAKGGGDCHCGDCKCCSTAAC |
| Cheiracanthium lascivum | ClasMT2 | IBQA01031776 | MADVEKKSCCAEPKECCKDESCCQEGKGPCHTEKEGCADSCQKKPCGCKAGEDCKCTDGKCAC |
| | ClasMT3L | IAVB01025709 | MDKKCGPNCKCTKCRCCDSAKPCCTGAVGKADCGCEDCKCCSNSSCKSQKSCGQGCQCGDCKCCSS SPCQAAKSSCGSSCKCDVCTCCSAAKDGGACHCGDCKCCTAAVC |
| | ClasMT3S | IBOH01015086 | MPGGPCGSNCTCEKCRCCSAATKCSSGCKCDNCRCCQRGQGQCQCGDCKCCGAAKC |
| Clubiona basarukini | CbasMT2 | IBFW01007189 | MADVEKTSCCTEPKECCKDESCCENGQGACHTGKEECKDTCHKKACGCKGGEDCKCSEGKCGC |
| | CbasMT3L | IBNI01016319 | MDKRCGPNCKCTKCRCCDGGKGAAKADCGCEECRCCSSSGCGSQKSCGKGCQCGDCKCCSSSACQT SACCGSNCKCDVCTCCSAAKDGGDCHCGDCKCCSQAKC |
| | CbasMT3S | IBFW01014451 | MPGGPCGTNCTCDKCRCCSAAAKCSSGCKCDTCRCCQRSQKCQCGDCKCCGASKC |
| Coriarachne fulvipes | CfulMT2 | IBOC01038815 | MADVEKTSCCTEPKECCKDESCCENGQGECHTGKEGCKDTCHKKACGCKADEECKCTEGKCAC |
| | CfulMT3L | IBOC01030735 | MDKCGPNCKCTKCRCCDAAKSVTPACCCQDSAKKANCACDDCQCCKSSSCATQKSDCGKGCQCNCK CCSGGTQKDSCGSNCKCDVCTCCSAAKDSGACHCGDCRCCSAASC |
| | CfulMT3S | IBOC01006221 | MPGGPCGSNCTCEKCRCCTAASKCTSSCKCDTCRCCQRSQKCNCGDCKCCGAAKC |
| Cyclosa alba | CalbMT2 | IAZE01025958 | MADVEKKTCCAEPKECCKDESCCQEGKGPCHTEKEGCADSCQKKPCGCKAGEDCKCKDGKCAC |
| | CalbMT3Sa | IAZE01015529 | MPCGTCGPSCTCDKCRCCGASSGPKCSANCNCGDCKCCGRTQANCRCGDCKCCGAAKC |

| | | | |
|---|---|---|---|
| | CalbMT3Sb | IAZE01014351 | MPGGPCGSNCNCEKCRCCSAASKCTANCKCDTCRCCQRNQQCNCGDCKCCSSAKC |
| Cyrtarachne akirai | CakiMT3S | ICFL01015310 | MPCGTCGASCTCEKCRCCGASASSKCSENCKCGDCRCCGRTSANCRCGDCKCCGAAKC |
| Cyrtophora cicatrosa | CcicMT3S | IBHY01004928 | MPCGTCGPSCSCEKCRCCGAAASSKCSANCKCVDCKCCGRSSANCRCGDCKCCGAAKC |
| Delena cancerides | DcanMT2 | ICHF01017029 | MADVEKTSCCTEPKECCKDESCCENGQGACHTGKEECKDTCHKKPCGCKAGEDCKCSEGKCAC |
| | DcanMT3L | ICHF01001373 | MERKCSPNCKCTKCQCCPASVKQTSKDCGCEECRCCKTSACATGKTCGKGCQCGDCKCCSSSGCQTSR ADKSCGASCKCDVCTCCSAATGGGICHCGDCKCCSTAVC |
| | DcanMT3S | ICHF01007191 | MPGGPCGSNCKCEKCRCCSAASKCGANCKCDTCRCCQRGQKCECGDCKCCGAAKC |
| Desis marina | DmarMT2 | GIWQ01214392 | MADVEKTTSCCTDPKECCKDDACCEQGQGPCHTGKEECKDTCHKKACGCKAGEDCKCSGGKCGC |
| | DmarMT3L | GIWQ01151592 | MEKAPCGSNCKCTKCRCCDGAQCGMKAGNCGCAECQCCSSSACQSGGKCCGKGCRCEDCKCCSAAS GQTKASCCCTNCKCDVCSCCSAAKGSGTCPCGDCKCCPTAPC |
| | DmarMT3S | GIWQ01313974 | MPGGPCGTNCTCEKCRCCSAAAKCSPNCKCDTCRCCQRNQKCECGDCKCCGAAKC |
| Diaea subdola | DsubMT2 | ICGF01004055 | MADVEKKSCCAEPKECCKDESCCQEGKGPCHTEKEGCADSCQKKPCGCKAGEDCKCTDGKCAC |
| | DsubMT3L | ICGF01021847 | MDKCGPNCKCTKCRCCDAAKSATPACCQAKQTKCGCDDCLCCKSSGCGAQKSDCGKGCQCNCKCCS SGAQKDSCGSNCKCDVCTCCSATKDSGACHCGNCKCCSAASC |
| | DsubMT3S | IBBD01042702 | MPGGPCGSNCTCEKCRCCSAASKCTASCKCDTCRCCQRNQKCECGDCKCCSASKC |
| Dictyna foliicola | DfolMT2 | IAXI01039432 | MADVENKSCCAEPKECCKDESCCQEGKGPCHTDKDSSCADSCQKKACGCKSGEDCKCKDGKCDC |
| | DfolMT3Sa | IAXI01008702 | MPGGPCGSNCTCEKCRCCSAAAKCGSSCKCDTCRCCQRGQRCECGDCKCCGAAKC |
| | DfolMT3Sb | IAXI01012824 | MNCGTCGPNCTCEKCRCCGAAAAPKCSANCNCGDCKCCGRTSANCRCGDCKCCGAAKC |
| Dolomedes sp. | DspMT3L | IBQW01022981 | MDKCGANCKCTKCRCCDAAKPATLSCCQSAATKAACGCDDCLCCNSSGCGAQKASDKGCQCECKCCS SSNQKNRCGSNCKCDVCTCCSSATGSGNCHCGDCKCCSAAVC |
| Dolomedes triton | DtriMT2 | GGRN01026674 | MADVEKTSCCTEAKECCKDESCCENGQGACHTGKEECKDSCHKKACGCKAGEDCKCSEGKCAC |
| | DtriMT3L | GGRN01035980 | MDKCGANCKCTKCRCCDAAKPATLSCCQSAATKAACGCDDCLCCNSSGCGAQKASDKGCQCECKCCS SSSQKNRCGSNCKCDVCTCCSSATGSGNCHCGDCKCCFAAVC |
| | DtriMT3S | GGRN01043194 | MPGGPCGSNCTCEKCRCCSAAAKCSSGCRCDTCRCCQRGQKCQCGDCKCCGAAKC |
| Ebelingia kumadai | EkumMT2 | IANA01004259 | MADVEKTSCCTEPKECCKDESCCENGQGECHTGKEECKDSCHKKACGCKADEECKCSEGKCAC |
| | EkumMT3L | IANA01024013 | MDRCGPNCKCTKCRCCDAAKSATPACCQAAAKKTDCGCDDCLCCKSSSCGAQKSDCGKGCQCSCKCC AGAAQKDSCGSNCKCDVCTCCSAATDSGACHCGDCRCCSAATC |

| | | | |
|---|---|---|---|
| | EkumMT3S | IANA01042734 | MPGGPCGSNCSCEKCRCCSAASKCTASCKCDTCRCCQRGQKCNCGDCKCCGAAKC |
| Ebrechtella tricuspidata | EtriMT3L | IBTQ01004588 | MDKCGPNCKCTKCRCCDAAKSATPACCQAKKTSCKCDDCLCCTQKSDCGKECQCSCKCCSGPAQKDS CGSNCKCDVCTCCSDAKESGASHCGNCRCCSAASC |
| Enoplognatha abrupta | EabrMT3S | IBGU01031423 | MPCGTCGPNCTCEKCRCCGAAAAPKCSANCNCGDCKCCGRTSANCRCGDCKCCGAAKC |
| Episinus kitazawai | EkitMT2 | IBEG01009253 | MADVENKSCCAEPKECCKDESCCQEGKGPCHTEKEGCADSCQKKPCGCKAGEDCKCKDGKCAC |
| | EkitMT3Sa | IBEG01030932 | MPCGTCGPNCTCEKCRCCGAAAAPKCSANCNCGDCKCCGRTSANCRCGDCKCCGAAKC |
| | EkitMT3Sb | IBEG01022842 | MPGGPCGSNCTCEKCRCCSAASKCTASCKCDTCRCCQRNQKCECGDCKCCSASKC |
| | EkitMT3Sc | IBEG01030305 | MAGPCGANCTCDKCRCCSAASKCTASCKCDTCRCCQRSQNCNCGDCKCCGAAKC |
| | EkitMT3Sd | IBEG01032665 | MPGPCGENCKCEKCRCCSAASKCTSNCKCDVCKCCERSQKCNCGDCKCCGEAKC |
| Eriovixia sakiedaorum | EsakMT2 | IBEM01035248 | MADVEKKSCCAEPKECCKDESCCQEGKGPCHSEKEGCADSCQKPCGCKAGEDCKCKDGKCSC |
| | EsakMT3S | IBEM01002195 | MPCGTCGPSCTCEKCRCCGAAAAPKCSANCQCGDCKCCGRSSANCRCGDCKCCGAAKC |
| Falconina gracilis | FgraMT2 | IBPA01038432 | MADVEKTSCCTEPKECCKDESCCENGQGACHTGKEECKETCHKKPCGCKAGEDCKCSEGKCAC |
| | FgraMT3L | IBPA01036519 | MEKKCGANCKCTKCRCCDSARQAAPPCCTAAAAKKAGCSCDDCTCCKSSSCTSQKTCGKGCKCGDCQ CCSSSGCQGDKSCCGSGCKCDVCTCCSAAKGGGDCHCGDCRCCPSATC |
| | FgraMT3S | IBPA01019018 | MPGGPCGPNCTCERCRCCSAAVRCGAGCKCDTCKCCQRGEKCQCGDCKCCRAAKC |
| Gibbaranea bituberculata | GbitMT2 | IBVP01006635 | MADVEKKSCCAEPKECCKDESCCQEGKGPCHTEKEGCADSCQKKPCGCKAGEDCKCKDGKCAC |
| | GbitMT3S | IBVP01005554 | MPCGTCGPNCTCEKCRCCGAAAAPKCSANCNCGDCKCCGRTSANCRCGDCKCCGAAKC |
| Hasarius adansoni | HadaMT2 | GJQJ01025370 | MADVEKKTCCTEPKECCKDESCCQNGQGECHTGKEECKDSCQKKACGCKAGEDCKCSDGKCGC |
| | HadaMT3L | IBMC01008950 | MEKKCGPNCKCTKCRCCNAARSAAPSCCTAASAKTADCGCDDCTCCQKASCCGSQKTCGKGCDCGDC KCCSSSSCQAGKSCCGSNCRCDDCNCCSAAKGSGDCHCGNCRCCSAASC |
| | HadaMT3S | GJQJ01012079 | MPGGPCGSNCTCEKCRCCSAASKCSSGCKCDTCRCCQRNQSCNCGDCKCCRAAKC |
| Heriaeus mellotteei | HmelMT2 | IAJG01014351 | MADVEKTSCCTEPKECCKDESCCENGQGACHTGKEECKDSCHKKPCGCKEGEDCKCSDGKCGC |
| | HmelMT3L | IAJG01015975 | MDKCGPNCKCTKCRCCDAAKSATPTCCQDAAKKTDCRCEDCQCCKSSSCGTQKSDCGKGCQCNCKCC SGDTQKDSCGSNCKCDVCTCCSAAKDSGACHCGDCRCCSAASC |
| | HmelMT3S | IAJG01036463 | MPGGPCGSNCTCEKCRCCSAASKCTSSCKCDTCRCCQRSQKCNCGDCKCCGAAKC |

| | | | |
|---|---|---|---|
| Heteropoda venatoria | HvenMT2 | IBFQ01010827 | MADVEKTSCCTEPKECCKDESCCENGQGACHTGKEECKDTCHKKPCGCKAGEDCKCSEGKCAC |
| | HvenMT3L | IBFQ01012557 | MERKCGASCKCTKCQCCPESVKQASKDCGCEECRCCKSSPCASGKTCGKGCQCGDCKCCSSSACKTKSCGTGCKCDVCTCCSAATGGGNCHCGDCKCCSTAKC |
| | HvenMT3S | IAGF01017184 | MPGGPCGSNCTCEKCRCCSAASKCGPGCRCDNCRCCQRGQNCQCGDCKCCGAAKC |
| Hylyphantes tanikawai | HtanMT2 | IAQG01046202 | MADVEKKSCAESKDCCKDDPECCQEGKGPCHTDKDSSCADSCQKKACGCKSGEDCKCKDGKCEC |
| | HtanMT3Sa | IAQG01038721 | MPCGTCGPNCTCEKCRCCGAAAAPKCSANCNCGDCKCCGRTSANCRCGDCKCCGAAKC |
| | HtanMT3Sb | IAQG01034859 | MPGGPCGSNCTCEKCRCCSAAAKCTSSCKCDTCRCCQRSQKCECGDCKCCSASKC |
| Hypsosinga sanguinea | HsanMT2 | IATX01006580 | MADVENKSCCAEPKECCKDESCCQDGKGPCHNEKEGCADSCQKKACGCKAGEDCKCKDGKCGC |
| | HsanMT3S | IATX01035236 | MPCGTCGPSCTCEKCRCCGAGAASASKCSANCKCADCKCCGRTSANCRCGDCKCCDAAKC |
| Labulla thoracica | LthoMT2 | ICGP01036048 | MADVENKSCCAEPKECCKDESCCQEGKGPCHTEKEGCADSCQKKPCGCKAGEDCKCQDGKCAC |
| | LthoMT3S | ICGP01002053 | MPCGTCGPNCTCDKCRCCGAAASSKCSANCNCGDCKCCGRTSANCRCGDCKCCGAAKC |
| Larinioides cornutus | LcornMT3S | IBUI01018852 | MPCGTCGPSCTCEKCRCCGAAATSAKCSANCKCVDCKCCGKTGANCRCGDCKCCGAAKC |
| Latrodectus geometricus | LgeoMT2 | GFCZ01010332 | MADVEKPCCPEPKECCKDESCCDKGKGPCHAGEGGGEACQQKPCGCKGGEECKCKEGKCGC |
| | LgeoMT3S | GBJM01039547 | MPGPCGPNCNCEKCRCCSAASKCTSNCKCDVCRCCSRSQRCNCGDCKCCGEAKC |
| Latrodectus hesperus | LhesMT2 | GBJN01124720 | MADVEKPCCPEPKECCKDESCCDQGKGPCHAGEGGGDACQQKPCGCKAGEDCKCKEGKCGC |
| | LhesMT3Sa | GBCS01027129 | MPGPCGPNCNCEKCRCCSAASKCTSSCKCDVCHCCSRSRRCNCGDCKCCGEAKC |
| | LhesMT3Sb | GBJN01094922 | MPGPCGPNCNCEKCRCCSAASKCTSNCKCDVCRCCSRNQKCNCGDCKCCGEAKC |
| Lehtinelagia evanida | LevaMT3L | IBFM01000596 | MDKSCGPNCKCTKCRCCDAAKSASPACCAAKKTDCGCDDCQCCKSSSCGTQKSDCGKECQCNCKCCSGAAQRDSCGSNCKCDVCTCCSAAKDSGNCHCGDCRCCSAASC |
| Loxosceles laeta | LlaeMT2 | EY188643 | MADVEKAPCCDSPKECCKDATCCDEGKGPCHGPDEKCSDSASKKKCGCKAGEDCKCSEGKCAC |
| Lycosa coelestis | LcoeMT2 | IBIG01002848 | MADVEKTSCCTEPKECCKDESCCENGQGACHTGKEECKDSCHKKACGCKAGEDCKCSDGKCGC |
| | LcoeMT3L | IBIG01003521 | MDKCGPNCKCTKCRCCDAAKDSTPACCQAAAKKTSCGCDDCSCCSSSGCCEQKASGKGCQCSCKCCSPAASQKNTCGSSCKCDVCSCCSAAKGSGDCHCGNCKCCSTACC |
| | LcoeMT3Sa | IBVM01020795 | MPGGPCGSNCTCEKCRCCSAASKCSAGCRCDTCRCCQRGQKCQCGDCKCCGAAKC |

| | | | |
|---|---|---|---|
| | LcoeMT3Sb | IBIG01009944 | MPGGPCGANCSCEKCRCCSAAASSGGRCGPNCRCDVCRCCNRGQNCRCGDCRCCGASKC |
| Lysiteles coronatus | LcorMT2 | IBDW01045585 | MADVEKTSCCTESKECSKDESCCENGQGECHTGEEKCKDTCHKKACGCKADEECKCSEGKCAC |
| | LcorMT3L | IBDW01040765 | MEKNCGPNCKCTKCRCCNAAKGKDCGCETCRCCDASSCKSAKSCGSGCDCRDCKCCSATSCCQTSSC CGSGCKCDKCTCCAAAKGKGTCNCGDCKCCSTAKC |
| | LcorMT3Sa | IBDW01030729 | MPGGPCGSNCTCEKCRCCTAASKCTASCKCDTCRCCQRSQKCNCGDCKCCGAAKC |
| | LcorMT3Sb | IBDW01005672 | MPGGPCGSNCSCEKCRCCSAAASASPSKCGPNCKCDVCRCCNRGQNCRCGDCKCCGASKC |
| Lysiteles maior | LmaiMT2 | ICAK01041707 | MADVEKTSCCTEPKECCKDESCCENGQGECHTGEEKCKDTCHKKACGCKADEECKCAEGKCAC |
| | LmaiMT3L | ICAK01042855 | MEKCGPNCKCTKCRCCDAAKSATCCQDAAKKTNCGCDDCQCCKSAGSGAQKSDCGKGCQCNCKCCS GGKQKDGCESNCKCDRCTCCPVTKDSGVCHCGFCRCCDLASF |
| | LmaiMT3Sa | ICAK01036021 | MPGGPCGSNCTCEKCRCCTAASKCTASCKCDTCRCCQSGQKCNCGDCKCCGAAKC |
| | LmaiMT3Sb | ICAK01032991 | MPGGPCGSNCSCEKCRCCSAAASASPSKCGPNCKCDVCRCCNRGQNCRCGDCKCCGASKC |
| Macracantha hasselti | MhasMT2 | IBKA01027009 | MADVENKSCCAEPKECCKDESCCQEGKGPCHTEKEGCADSCQKKPCGCKAGEDCKCQDGKCAC |
| | MhasMT3S | IBKA01028729 | MPCGTCGPNCTCDKCRCCGAAASSKCSANCNCGDCKCCGRTSANCRCGDCKCCGAAKC |
| Mallinella shimojanai | MshiMT2 | IAKA01018968 | MADVEKTSCCTEAKECCKDESCCQNGQGSCHTGKEECKDSCHKKPCGCKASEECKCSDGKCPC |
| | MshiMT3L | IAKA01013492 | MDKRCGANCKCTKCRCCDAQKPKQAAGTDCGCEECRCCKSAPCSSGKTCGKGCQCGDCKCCSSSACQ TSSCGTSCACDVCTCCSASKGDGDCHCGDCKCCSSAKC |
| | MshiMT3S | IAKA01005514 | MPGGPCGPNCSCEKCRCCSASAAPKCSASCKCDKCRCCQRTQKCQCGDCKCCAGSTC |
| Mangora placida | MplaMT2 | IBXG01004564 | MADVEKKSCCAEPKECCKDETCCQEGKGPCHTEKESCQKKPCGCSTGEECKCKDGKCAC |
| | MplaMT3S | IBXG01043951 | MPCGTCGPNCTCEKCRCCGAAASSAKCGPNCKCVDCKCCGKTSANCRCGDCKCCGAAKC |
| Marpissa muscosa | MmusMT2 | GJIM01328652 | MADVEKKSCCTEAKECCKDESCCQNGQGTCHTGKEECKDSCKTKACGCKAGEDCKCSDGKCAC |
| | MmusMT3L | GJIM01217943 | MERKCGPNCRCTKCRCCDAAKQSAPPCCTAAAASKKADCGCDDCTCCQKSACCGSEKTCGKGCQCGD CRCCSSSSCQAGKSSCGSNCKCDVCTCCSAAKSGGDCHCGDCRCCSAASC |
| | MmusMT3S | GJIM01320864 | MPGGPCGSNCTCEKCRCCSAATKCNSGCKCDPCRCCGKGQNCRCGDCKCCGAAKC |
| Marpissa yawatai | MyawMT2 | IBNG01005425 | MADVEKKSDCTESKECCKNEMPHEPCHTDKEDCKDCCKTKECGCKAGEDCKCSDGKCGC |
| | MyawMT3L | IBNG01023303 | MERKCGPNCRCTKCRCCDAAKQSAPPCCTAAAASKKAACGCDDCTCCQKSACCGSEKTCGKGCQCGD CKCCSASSCQAGKSSCGSNCKCDVCTCCSAAKSGGDCHCGDCRCCSAASC |

| | | | |
|---|---|---|---|
| | MyawMT3S | IBNG01051378 | MPGGPCGSNCTCEKCRCCSAATKCNSGCKCDPCRCCGKGQNCRCGDCKCCGAAKC |
| Mecynogea lemniscata | MlemMT2 | ICBE01019609 | MADVEKKSCCAEPKECCKDESCCQEGKGPCHTEKDGCADSCQKKPCGCKAGEDCKCKDGKCAC |
| | MlemMT3S | ICBE01013695 | MPCGTCGPSCSCEKCRCCGAAAAAPKCSANCKCVDCKCCGKTGANCRCGDCKCCGAAKC |
| Micrathena gracilis | MgraMT2 | IAWP01018183 | MADVEKKPCCAEPKECCKDESCCQEGKGPCHTEKDGCADSCQTKPCGCKAGEDCKCTDGKCSC |
| | MgraMT3S | IAWP01035947 | MPCGTCGPSCSCEKCRCCGAAASSKCSPNCKCDNCQCCGRSSANCRCGDCKCCGASKC |
| Microdipoena pseudojobi | MpseMT2 | ICBP01032588 | MADVEKKCCAEPKDCCKEDPSCCQDGKGSCHTDKDSCADSCDKKACGCKAGEDCKCKDGKCAC |
| | MpseMT3L | ICBP01040719 | MEKNCGPNCKCTKCRCCNAAKGKDCGCETCRCCDASSCKSAKSCGSGCDCRDCKCCSATSCCQTSSC CGSGCKCDKCTCCAAAKGKGTCNCGDCKCCSTAKC |
| | MpseMT3Sa | ICBP01045714 | MPGGPCGTNCTCEKCRCCGAAAKCSASCKCETCRCCQRGQNCRCGDCKCCGASKC |
| | MpseMT3Sb | ICBP01037319 | MPGPCGPNCTCDKCRCCTAATKCSEGCKCDTCRCCERGQKCQCGDCKCCGAAKC |
| Misumenops bellulus | MbelMT2 | IAGN01028376 | MADVEKTPCCTEAKECCKDQKGQGECHTGKEECKDTCHKKACGCKADEECKCSEGKCAC |
| | MbelMT3L | IAGN01039957 | MDKCGPNCKCTKCRCCDAAKSATPSCCQDAAKKANCGCDDCQCCKSSSCGTQKSDCGEGCQCSCKC CAGATQKDSCGSNCKCDVCTCCSAAKDSGACHCGDCRCCSTASC |
| | MbelMT3Sa | IAGN01029399 | MPGGPCGSNCTCEKCRCCSAASKCTASCRCDTCRCCQRSQKCNCGDCKCCGAAKC |
| | MbelMT3Sb | IAGN01017054 | MPGGPCGANCSCEKCRCCSAAASASASRCGPNCKCDVCRCCNKGQNCRCGDCKCCGASKC |
| Morebilus sp. | MspMT3L | IAGI01044335 | MEKKCGPNCKCTKCRCCDAAKTAPCCTAAAAKKADCGCDDCHCCKSSSCASQKTCGKGCQCGDCKCC SSSACQTSKSCCGSSCKCDVCTCCSAAKDGGVCHCGNCRCCETAAC |
| Myrmarachne inermichelis | MineMT2 | ICJC01044708 | MADVEKPCCAEPKECCKDESCCQEGKGPCHTEKEGCGDSCHKKPCGCKAGEDCKCKDGKCAC |
| | MineMT3L | ICJC01003799 | MEKKCGPNCKCTKCRCCDSAKQAAPPCCTAAAAKKADCGCNDCSCCQNSSCCGSKGCQCSDCKCCSS SSCQTSKSSCGSSCKCDVCTCCSAAKGSGDCHCGNCKCCSAASC |
| | MineMT3S | ICJC01041083 | MPCGTCGPNCTCEKCRCCGAAAAPKCSANCNCGDCKCCGRTSANCRCGDCKCCGAAKC |
| Neon minutus | NminMT3L | IAIC01046021 | MENKCGPNCKCTKCRCCDAAKQATPSCCAAAAAKKADCGCDDCTCCQKSSCGSQKSCGKGCQCSDCK CCSSAACHASKSCCGSSCKCDVCTCCSAAKGSGDCHCGDCRCCSAASC |
| Neoscona arabesca | NaraMT2 | IBED01024418 | MADVEKKSCCAEPKECCKDESCCQEGKGPCHTEKEGCADSCQKKPCGCKAGEDCKCTDGKCAC |
| | NaraMT3S | IBED01005087 | MPCGTCGPSCTCDKCRCCGAAAAPKCSANCQCGDCKCCGRTSATCRCGDCKCCGAAKC |
| Nephila clavipes | NclaMT2a | GFKT010927754 | MADVEKPCCAEPKECCKDESCCDEGKGPCHTDGKEDCADSCHKKPCGCKAGEDCKCKDGKCAC |

| | | | |
|---|---|---|---|
| | NclaMT2b | GFKT012135641 | MADVEKTACCAEPKECCKDESCCQEGKGPCHTEKESCGDSCHKKPCGCKAGEDCKCKDGKCAC |
| | NclaMT3S | GFKT011191296 | MPGGPCGSNCNCEKCRCCSAASKCTSGCKCDTCRCCQRNQKCECGDCKCCGAAKC |
| Nephilengys cruentata | NcruMT2 | GEWZ01004248 | MADVEKTACCAEPKECCKDESCCQEGKGPCHTEKESCGDSCHKKPCGCKAGEDCKCKDGKCAC |
| Nesticus yesoensis | NyesMT2 | IBBO01018781 | MADVEKKCCDEPKECCKDESCCQEGKGSCHTEKDGSCADKCHAKKACGCTAGEECKCKEGKCGC |
| | NyesMT3S | IBBO01028050 | MPCGTCGPNCTCEKCRCCGAAAAPKCSANCNCGDCKCCGRTSANCRCGDCKCCGAAKC |
| Nihonhimea mundula | NmunMT2 | IAVV01016525 | MADVEKPCCAEPKECCKDESCCQEGKGPCHTEKEGCADSCQKKPCGCKAGEDCKCKDGKCAC |
| | NmunMT3S | IAVV01005513 | MPCGTCGPNCTCEKCRCCGAAAAPKCSANCNCGDCKCCGRTSANCRCGDCKCCGAAKC |
| Nuctenea umbratica | NumbMT3S | IAHU01040313 | MPCGTCGPSCTCEKCRCCGAATASASKCSANCKCADCKCCGRTSANCRCGDCKCCGAAKC |
| Octonoba grandiconcava | OgraMT2 | IBKN01001004 | MADVEKSCCAETKDCCDVDANCCENGKGTCHEKDADCCKDGCHTKKPCGCKANEACKCSEGKCAC |
| | OgraMT3Sa | ICDQ01018272 | PGGPCGANCTCEKCRCCGAASRCSDTCKCDTCRCCQKGRDCRCGDCKCCGASKC |
| | OgraMT3Sb | ICDQ01030917 | MPGGPCGANCTCEKCRCCGAAAKCSSSCKCETCRCCQRGQNCRCGDCKCCGASKC |
| Octonoba grandiprojecta | OgrpMT2 | ICKD01013852 | MADVEKTSCCTEAKECCKDESCCENGQGACHTGKEECKDSCHKKACGCKAGEDCKCSEGKCAC |
| | OgrpMT3S | ICKD01006318 | MHGGPCGANCTCEKCRCCGAASRCSDTCKCDTCRCCQKGRDCRCGDCKCCGASKC |
| Octonoba okinawensis | OokiMT2 | IALW01017951 | MADVEKSCCAETKDCCDVDANCCENGKGTCHEKDADCCKDGCHTKKPCGCKANEACKCSEGKCAC |
| | OokiMT3Sa | IALW01003257 | MPGGPCGANCTCEKCRCCGAAAKCSSSCKCETCRCCQRGQNCRCGDCKCCGASKC |
| | OokiMT3Sb | IALW01034307 | MPGGPCDPNCTCEKCRCCGAAAKCSSSCKCETCRCCQRGQNCRCGDCKCCGASKC |
| | OokiMT3Sc | IALW01006618 | MTGGTCGANCTCEKCRCCGAASRCSDTCKCDTCRCCQKGRDCRCGDCKCCGASKC |
| Octonoba sybotides | OsybMT2 | IAZZ01001752 | MADVEKSCCAETKDCCDVDANCCENGKGTCHEKDADCCKDGCHTKKPCGCKANEACKCSEGKCAC |
| | OsybMT3Sa | IAUX01017549 | MPGGPCGANCTCEKCRCCGAASRCSDTCKCDTCRCCQKGRDCRCGDCKCCGASKC |
| | OsybMT3Sb | IAUX01008602 | MPGGPCGANCTCEKCRCCGAAAKCSASCKCETCRCCQRGQNCRCGDCKCCGASKC |
| Octonoba varians | OvarMT2 | IAST01008113 | MADVEKSCCAETKDCCDVDANCCENGKGTCHEKDADCCKDGCHTKKPCGCKANEACKCSEGKCAC |

| | | | | |
|---|---|---|---|---|
| | | OvarMT3Sa | IAST01011549 | MPGGPCGANCTCEKCRCCGAASRCSDTCKCDTCRCCQKGRDCRCGDCKCCGASKC |
| | | OvarMT3Sb | IAST01017713 | MPGGPCGSNCTCEKCRCCGAAAKCSASCKCETCRCCQRGQNCRCGDCKCCGASKC |
| | Octonoba yesoensis | OyesMT2 | IAQA01003517 | MADVEKSCCAETKDCCDVDANCCENGKGTCHEKDADCCKDGCHTKKPCGCKANEACKCSEGKCAC |
| | | OyesMT3Sa | IAQA01022141 | MPGGPCGANCTCEKCRCCGAASRCSDTCKCDTCRCCQKGRDCRCGDCKCCGASKC |
| | | OyesMT3Sb | IAQA01008891 | MPGGPCGTNCTCEKCRCCGAAAKCSASCKCETCRCCQRGQNCRCGDCKCCGASKC |
| | Oecobius navus | OnavMT2 | IBWY01033478 | MADVENKSCCAEPKECCKDESCCQEGKGPCHTEKEGCADSCQKKPCGCKAGEDCKCQDGKCAC |
| | | OnavMT3S | IBWY01005004 | MPCGTCGPNCTCDKCRCCGAAASSKCSANCNCGDCKCCGRTSANCRCGDCKCCGAAKC |
| | Otacilia luna | OlunMT2 | ICGC01035958 | MADVEKTSCCTDAKECKDDSCCDNGQAACHTGKEECAETCHKKPCGCKAGEDCKCSDGKCAC |
| | | OlunMT3L | ICGC01023166 | MERCGPNCKCTKCRCCDAGKSGAASGCGCKDCSCCGSSSCGSQKTCGKGCQCKDCKCCSSSGCQTGKSCCGSGCKCDVCTCCSAAQGGGACHCGDCKCCSSAKC |
| | | OlunMT3S | ICGC01018723 | MPGGPCGSNCTCEKCRCCSAASKCSAGCKCDTCRCCQRGEKCQCGDCKCCGAAKC |
| | Oxytate hoshizuna | OhosMT2 | IBYW01036832 | MADVEKTSCCTESKECCKDESCCENGQGECHTGKEECKDTCHKKACGCKADEECKCSEGKCAC |
| | | OhosMT3L | IBFD01027433 | MDKCGPNCKCTKCRCCDAAKSATPACCQAKKTSCKCDDCLCCTQKSDCGKECQCSCKCCSVPAQKDSCGSNCKCDVCTCCSAAKDSGASHCGSCRCCSAASC |
| | | OhosMT3Sa | IBIV01016281 | MPGGPCGPNCACEKCRCCSAASKCTAACKCDTCRCCQRNQKCNCGDCKCCGASKC |
| | | OhosMT3Sb | ICJU01039914 | MPGGPCGANCTCEKCRCCSSAATASASRCGPNCKCDVCRCCGRGQNCRCGDCKCCSASKC |
| | Ozyptila matsumotoi | OmatMT2 | IBFR01041374 | MADVEKTSCCTETKECCKDESCCENGQGECHTGKEGCKDTCHKKACGCKAGEECKCTEGKCAC |
| | | OmatMT3L | IBFR01009325 | MDKCGPNCKCTKCRCCDAAKSASPCCCQSAAKKANCRCDDCQCCKSSSCGTQKSDCGKGCQCNCKCCSGGTQKDSCGSNCKCDVCTCCSAAKDSGACHCGDCRCCSAASC |
| | | OmatMT3S | IBFR01032300 | MPCGTCGPNCTCEKCRCCGAAAAPKCSANCNCGDCKCCGRTSANCRCGDCKCCGAAKC |
| | Pachygnatha quadrimaculata | PquaMT2 | IALH01014478 | MADVEKPCCAEPKECCKDESCCQEGKGPCHTEKEGCGDSCHKKPCGCKAGEDCKCKDGKCAC |
| | | PquaMT3Sa | IALH01010477 | MPCGTCGPNCTCEKCRCCGAAAAPKCSANCNCGDCKCCGRTSANCRCGDCKCCGAAKC |
| | | PquaMT3Sb | IALH01010495 | MPGGPCGSNCTCEKCRCCSAASKCTASCKCDTCRCCQRNQKCECGDCKCCSASKC |
| | Pandava laminata | PlamMT3L | IASI01033194 | MEGKTCGSNCKCTKCRCCDAAKTGKECGCEECRCCKSASCGSQQSTCGKGCTCGDCKCCSSADCQQKSCGSGCKCDVCSCCTAATGGGVCHCGNCKCCPSSGC |

| | | | |
|---|---|---|---|
| Pandercetes sp. | PspMT3L | IBWH01021393 | MEKKCGSSCKCTKCQCCPASVKQASKDCGCEECRCCKSSPCASGKTCGKGCQCGDCKCCSSSACQTSS CGAGCKCDVCTCCSAATGGGNCHCGDCKCCSAAVC |
| Parasteatoda ryukyu | PryuMT2a | IBPF01015651 | DVEKTSCCTEPKECCKDESCCENGQGACHTGKEECKDSCHKKPCGCKEGEDCKCSDGKCGC |
| | PryuMT2b | ICDI01032605 | MADVEKPCCDEPKDCCKDESCCKEGKGPCHTEKEGCGDSCHKKACGCKVGEDCKCSGGKCEC |
| | PryuMT2c | ICDI01004733 | MADVEKKPCCAEPKECCKDESCCQEGKGPCHTEKEGCADSCQKKPCGCKAGEDCKCQDGKCAC |
| | PryuMT3S | IAVY01004002 | MPCGTCGPNCTCEKCRCCGAAAAPKCSANCNCGDCKCCGRTSANCRCGDCKCCGAAKC |
| Parasteatoda tepidariorum | PtepMT2 | IBLE01032418 | MADVENKSCCAEPKECCKDESCCQEGKGPCHTDKDSSCADSCQKKACGCKSGEDCKCKDGKCDC |
| | PtepMT3S | IACA01037330 | MPGGPCGPNCTCEKCRCCSAASKCTSNCKCDNCRCCQRSQKCNCGDCKCCGEAKC |
| Pardosa amentata | PameMT2 | GJIU01181368 | MADVEKTSCCTESKECCKDESCCENGQGACHTGKEECKDSCHKKACGCKAGEDCKCSDGKCGC |
| | PameMT3L | GJIU01357226 | MDKCGANCKCTKCRCCDAVKESTPACCQAAARKTTCGCDDCSCCSSSGCCEPKAVGKGCQCNCKCCS PASSQKNTCGSSCKCDVCSCCSAAKGSGDCHCGNCKCCSTACC |
| | PameMT3S | GJIU01318898 | MPGGPCGANCSCEKCRCCSAAASSGGRCGPNCRCEVCRCCNRGQNCRCGDCRCCSASKC |
| Pardosa pseudoannulata | PpseMT2 | GGRD01042272 | MADVEKTSCCTEPKECCKDESCCENGQGACHTGKEECKDSCHKKPCGCKEGEDCKCSDGKCGC |
| | PpseMT3L | GGRD01246346 | MDKCGANCKCTKCRCCDAAKESTPACCQAAAKKTSCGCDDCSCCSSSGCCEQTASGKGCQCSCKCCS PASSQKKACGSSCKCDVCTCCSAAKGSGDCHCGNCKCCSTACC |
| | PpseMT3S | GGRD01193535 | MPGGPCGSNCTCEKCRCCSAASKCSAGCRCDTCRCCQRGQKCQCGDCKCCGAAKC |
| Perilla teres | PterMT2 | IBAD01021535 | MADVENKSCCAETKECCKDESCCQEGKGPCHTEKEGCADSCQKKPCGCKAGEDCKCKDGKCAC |
| | PterMT3S | IBAD01018297 | MPCGTCGPNCSCEKCRCCGAASASKCSANCKCSDCKCCGRTSANCRCGDCKCCDAAKC |
| Pharta brevipalpus | PbreMT2 | IAWA01034742 | MADVEKTSCCTESKECCKDESCCENGQGACHTGKEECKDSCHKKACGCKADEDCKCAEGKCAC |
| | PbreMT3L | IAWA01009108 | MDKCGPNCKCTKCRCCDAAKSAKPSCCQTAKKSCECDDCRCCNSSSCQTGKECQCNCKCCSASSEQK DSCGSNCKCDVCTCCSASKESGDCHCGDCKCCSGAAC |
| | PbreMT3S | IAWA01017274 | MPGGPCGANCSCEKCRCCSAAGACGSSSRCSPNCKCDVCRCCNRGQNCRCGDCKCCSASKC |
| Phidippus clarus | PclaMT2 | ICBG01047316 | MADVEKKSCCTEPKECCKDESCCQNGQGSCHTGKEECKDSCQTKACGCKAGEDCKCSDGKCAC |
| | PclaMT3L | ICBG01041784 | MEKKCGPNCRCTKCRCCDAAKQAAPPCCTAAAAKKADCGCDDCTCCQKSSCCGSEKACGKGCQCGDC RCCSSSACQTGKSSCGSNCKCDVCTCCSAAKSGGDCHCGDCRCCSAASC |
| Philodromus emarginatus | PemaMT2 | IBPM01002942 | MADVEKTSCCTEPKECCKDESCCQNGQGACHTDKEECKESCHKKPCGCKTGEECKCSSDGKCAC |

| | | | |
|---|---|---|---|
| | PemaMT3L | IBPM01010691 | MDSCGPNCKCSKCRCCDAATQSALTCCCAASGKKSCSCTDCTCCKSSSCCEKQKSCGEGKCDNCKCCEASPCQTSKSSCGSSCKCDVCTCCSAAKGGADCHCGNCKCCSAATC |
| | PemaMT3S | IBPM01033659 | MPCGTCGPNCTCEKCRCCGAAAAPKCSANCNCGDCKCCGRTSANCRCGDCKCCGAAKC |
| Phoneutria nigriventer | PnigMT2 | GFUY01017528 | MADVEKTSCCTEQKECCKDESCCENGQGACHTGKEECKDTCHKKACGCKAGEDCKCSEGKCAC |
| | PnigMT3Lpart | GFUY01014770 | CDCDDCTCCNSSGCGAQKTCGKGCQCSDCKCCSSASCHKASCGSDCKCDVCTCCSAAKSSGDCHCGNCRCCSAAVC |
| | PnigMT3S | GFUY01018717 | MPGGPCGANCTCERCRCCSAATKCSSGCKCDSCRCCQRGQNCQCGDCKCCGAAKC |
| Phoroncidia pilula | PpilMT2 | IBOK01000603 | MADVEKKCCAEPKDCCKEDPSCCQDGKGSCHTDKDSCADSCDKKACGCKAGEDCKCKDGKCAC |
| | PpilMT3L | IBOK01002666 | MEKNCGPNCKCTKCRCCNAAKGKDCGCETCRCCDASSCKSAKSCGSGCDCRDCKCCSATSCCQTSSCCGSGCKCDKCTCCAAAKGKGTCNCGDCKCCSTAKC |
| | PpilMT3S | IBOK01019512 | MPGGPCGPNCTCEKCRCCTAASKCTSNCKCDNCRCCQRSQKCNCGDCKCCGEAKC |
| Phrynarachne ceylonica | PceyMT2 | IAYL01007552 | MADVEKTSCCTEPKECCKDESCCENGQGKCHTGKEECKDTCHKKSCGCKADEECKCSEGKCAC |
| | PceyMT3L | IAYL01009101 | MDKCGPNCKCTKCRCCEAAKSATSCCCEDAAKKANCRCDDCQCCKSSDCGTQKSDCGEGCQCSCKCCSGAMQKDSCGSNCKCDVCTCCSAAKDSGACHCGDCRCCSTASC |
| | PceyMT3S | IAYL01023434 | MPGGPCGSNCSCEKCRCCSAAASASASRCGSNCKCDVCRCCNRGQNCRCGDCKCCSASKC |
| Phrynarachne katoi | PkatMT2 | IAOS01018527 | MADVEKTSCCTEPKECCKDESCCENGQGECHTGKEECKDTCHKKACGCKADEECKCSEGKCAC |
| | PkatMT3L | IAOS01005111 | MDKCGPNCKCTKCRCCDAAKSATTCCCQDAAKKANCRCDDCQCCKSSDCETQKSDCGKGCQCNCKCCSGATQKDSCGSNCKCDVCTCCSAAKDSGACHCGDCRCCSAASC |
| | PkatMT3S | IAOS01012376 | MPGGPCGSNCSCEKCRCCSAAASASASRCGSNCKCDVCRCCNRGQNCRCGDCKCCSASKC |
| Phycosoma japonicum | PjapMT2 | IAZW01000630 | MADVENKSCCAEPKECCKDESCCQEGKGPCHTDKDSSCADSCQKKACGCKSGEDCKCKDGKCDC |
| | PjapMT3Sa | IAZW01024263 | MPCGTCGPNCTCEKCRCCGAAAAPKCSANCNCGDCKCCGRTSANCRCGDCKCCGAAKC |
| | PjapMT3Sb | IAZW010044059 | MPCGNCGPNCTCEKCRCCGASSPKCSSNCKCGDCKCCNKKTNCRCGDCKCCGASPC |
| Pisaura bicornis | PbicMT2 | ICDN01011823 | MADVEKTSCCTESKECCKDDSCCENGKAACHAGKEECKESCHQKDSPCKCKSGEECKKEGCSC |
| | PbicMT3L | ICDN01027384 | MDKCGPNCKCTKCRCCDAAKPATPACCQEAAAKKAACGCDDCQCCSSSGCGTQKTCGKGCQCGDCKCCSAAASQKKTCGSNCRCDVCTCCSSAKGSGNCHCGDCKCCSAAVC |
| | PbicMT3S | ICDN01028901 | MPGGPCGSNCNCEKCRCCSAASASDSRCGPNCRCDVCRCCKRGANCRCGDCRCCSASKC |
| Pisaura lama | PlamaMT2 | IAQF01020027 | MADVEKKACCAEPKECCKDESCCQEGKGPCHTEKEGCADSCQKKPCGCKAGEDCKCKDGKCAC |

| | | | |
|---|---|---|---|
| | PlamaMT3L | IBMA01021694 | MDKCGPNCKCTKCRCCDAAKPATPACCQEAAAKKAACGCDDCQCCSSSGCGTQKTCGKGCQCGDCKCCSAATSQKKTCGSNCRCDVCTCCSSAKGSGNCHCGDCKCCSAAVC |
| | PlamaMT3S | IAQF01007099 | MPCGACGPNCTCEKCRCCGAAAAPKCSANCNCSDCKCCGRTSANCRCGDCKCCSAAKC |
| Pistius undulatus | PundMT3L | IBZD01006492 | MDKCGPNCKCTKCRCCDAAKSATPTCCQAAAMKVNCRCDDCQCCKSSGTQKSDCGKGCQCSCKCCSGATQKDSCGSNCKCDVCTCCSAAKDSGACHCGDCRCCSAASC |
| Platnickina sterninotata | PsteMT3S | IAGQ01013489 | MNCGTCGPNCTCEKCRCCGAAAAPKCSANCNCGDCKCCGRTSANCRCGDCKCCGAAKC |
| Plebs astridae | PastMT3S | IBUD01021694 | MPCGTCGPSCTCEKCRCCGAAAAPKCSSNCQCGDCKCCGKASASCRCGDCKCCSASKC |
| Poecilopachys australasia | PausMT3S | ICKB01002344 | MPCGTCGPNCTCDKCRCCGAAASSKCSANCNCGDCKCCGRTSANCRCGDCKCCGAAKC |
| Pseudopoda spirembolus | PspiMT2 | IBQV01000539 | MADVEKTSCCTETKECCKDESCCENGQGACHTGKEECKDSCHKKPCGCKAGEDCKCSEGKCAC |
| | PspiMT3L | IBKY01007427 | MERKCGSNCKCTKCQCCPASVKQASKECGCEECRCCKSSPCASGKTCGKGCQCGDCKCCSSSACKTSSCGTGCKCDVCTCCSAATGGGNCHCGDCKCCSSAVC |
| | PspiMT3S | IBKY01029357 | MPGGPCGSNCTCEKCRCCSAASKCSPSCKCDTCRCCQRGQKCQCGDCKCCGAAKC |
| | RhatrMT3L | IAUA01039453 | MERKCGPNCRCVKCRCCDASKQAAPPCCTAAAAKTADCGCDNCTCCQKSSCCGSEKACGKGCVCGDCKCCSSSPCQTSKSCCGSNCKCDVCTCCSAAKSGGDCQCGNCRCCSAASC |
| Runcinia acuminata | RacuMT3L | IAYK01009383 | MDKCGPNCKCTKCRCCDAAKSATPACCQAAAKKADCGCDDCQCCKSSGCGTQKSDCGKGCQCSCKCCSGAAQKESCGSNCKCDVCTCCSAAKDSGACHCGDCRCCSAASC |
| Schizocosa ocreata | SocrMT2 | GIZV01010417 | MADVEKTSCCTESKECCKDESCCENDQGACHTGKEECKDTCHKKACGCKAGEDCKCTDGKCGC |
| | SocrMT3L | GIZR01017868 | MDKCGANCKCTKCRCCDAAKESTPACCQAAAKKTSCGCDDCSCCSSSGCCEQKASGKGCQCSCKCCSPASRQKNTCGSSCKCDVCSCCSAAKGSGDCHCGNCKCCSTACC |
| | SocrMT3Sa | GIZU01029122 | MPGGPCGSNCTCEKCRCCSAASKCSAGCRCDTCRCCQRGQKCQCGDCKCCGAAKC |
| | SocrMT3Sb | GIZR01034889 | MPGGPCGANCTCEKCRCCSAAGSSGGRCGPNCRCDVCRCCNRGQNCRCGDCRCCSASKC |
| Shinobius orientalis | SoriMT2 | IBGW01039022 | MADVEKKSCCAEPKECCKDESCCQEGKGPCHTEKEGCADSCQKKPCGCKAGEDCKCQDGKCAC |
| | SoriMT3L | IBGW01024166 | MDKCGANCKCTKCRCCDAAKGAAPSCCPSAAKKSNCGCDDCRCCSSSGCEAQKASGKGCQCECKCCSSQRNTCGSNCKCDVCTCCSSAKGSGDCHCGNCKCCSAAGC |
| | SoriMT3S | IBGW01035513 | MPGGPCGPNCNCEKCRCCSAASKCTSNCKCDVCRCCQRGQRCNCGDCKCCGEAKC |
| | SrubMT3L | IBXR01041414 | MDKCGPNCKCTKCRCCDAAKTATPSCCQAAAKKSTCGCDDCQCCKSSSCGSQKTDCAKGCQCNCKCCSPASKQKDSCGSNCQCDVCTCCSAAKESGSCHCGDCRCCSAAAC |
| Singa perpolita | SperMT3S | IBIS01040974 | MPCGTCGPNCTCDKCRCCGAAASSKCSANCNCGDCKCCGRTSANCRCGDCKCCGAAKC |
| Sinopoda forcipata | SforMT2 | IAZL01034704 | MADVEKTSCCTESKECCKDESCCENGQGACHTGKEECKDSCHKKPCGCKAGEDCKCSEGKCAC |

| | | | | |
|---|---|---|---|---|
| | | SforMT3L | ICBI01018703 | MERKCGSNCKCTKCQCCPASVKQASKDCGCEECRCCKSSPCASGKSCGKGCQCGDCKCCSSSACQTS SCGTGCKCDVCTCCSAATGGGNCHCGDCKCCSAAVC |
| | | SforMT3S | IAZL01014300 | MPCGTCGSNCTCEKCRCCSAAASKCGPGCKCDNCKCCQRSQKCQCGDCRCCGAAKC |
| | Steatoda grossa | SgroMT2 | GBJQ01091123 | MADVEKPCCTEPKECCKDESCCDEGKGPCHANEGGDAACQQKPCGCKSGEDCKCKEGKCGC |
| | | SgroMT3S | GFDC01029402 | MPGGPCGPNCNCEKCRCCSAASKCTSNCKCDVCRCCSRGQKCNCGDCKCCGEAKC |
| | Stegodyphus dumicola | SdumMT2 | XR_004757882+SRA | MADVEKTGCCAEAKECAADNCSTECKTNCHDGKEECKDSCHKKPCGCKGTEDCKCSEGKCGC |
| | | SdumMT3L | XP_035211109 | METKCGVGCQCNQCKCCDVSKTRDSCCDKTGSKDCCCINSSCCKESGCNIAKQGTGDCNCADCKCCST ATCQTKSPSCKCDVCTCCQKSGGTDCHCGKCKCCPSAKS |
| | Stegodyphus lineatus | SlinMT2 | JT031711 | MADVEKTGCCAEAKECAADNCSTECKTNCHDGKEECKDSCHKKPCGCKGTEDCKCSEGKCGC |
| | | SlinMT3L | JT033686 | METKCGAECQCKECQCCDAAKTGDSCCDKTGSKDCCCTNSSCCKESGCNVSKQGTGNCNCTDCKCCS TATCQMKSPSCKCDVCTCCQKSGGTDCHCGKCKCCPSAKS |
| | | SlinMT3Sa | JT025713 | MPGGPCGSDCSCQNCRCCSAAAKCSANCKCDTCRCCQRSQTCRCGNCRCCSSAKC |
| | | SlinMT3Sb | JT027986 | MPGGPCGANCSCEKCRCCSAAAKCSAGCKCDVCRCCQRGQKCNCGDCRCCGAAKC |
| | Stegodyphus mimosarum | SmimMT2 | JT38942) | MADVEKTGCCAEAKECAADNCSTECKTNCHDGKEECKDSCHKKPCGCKGTEDCKCSEGKCAC |
| | | SmimMT3L | JT040526 | METKCGAGCQCKQCQCCDAAKAGDSCCDKTGSKDCCCINSPCCKASGCNISKEGTGNCNCTDCKCCST ATCQTKSPSCKCDVCTCCQKSGGTDCHCGKCKCCPSAKS |
| | | SmimMT3Sa | KFM81894 | MPGGPCGPNCTCRDCRCSSSAVKCNSGSKGETCRCSQNGQPCRCRDCKCRGGANR |
| | | SmimMT3Sb | JT036585 | MPGGPCGANCNCEKCRCCSAAAKCSAGCKCDVCHCCQRGQKCNCGDCRCCGAAKC |
| | Stegodyphus tentoriicola | StenMT2 | JT042210 | MADVEKTGCCAEAKECAADNCSTECKTNCHDGKEECKDSCHKKPCGCKGTEDCKCSEGKCGC |
| | | StenMT3S | JT042034 | MPGGPCGANCNCEKCRCCSAAAKCSAGCKCDVCRCCQRGQKCNCGDCRCCGAAKC |
| | Stephanopis cambridgei | ScamMT3L | ICHM01000361 | MDKCGPNCKCTKCRCCDAAKTATPSCCQAAAKKSTCGCDDCSCCKSSGCGSGKTDCGKGCQCNCKCC SSASKQKDSCGSSCQCDVCTCCSAAKESGSCHCGDCKCCSAAVC |
| | Storena cyanea | ScyaMT3L | IBGG01005528 | MERRCGANCKCTKCRCCEAGKTAAARQAGKDCGCDECRCCKDSSCGTGKSCGNGCECGDCKCCSSAA CQTSSCGSSCPCDVCTCCSASKGSGNCCCGDCKCCPTAKC |
| | Synagelides annae | SannMT2 | IBPH01030142 | MADVEKKSCCTEGKECCKDESCCENGEGACHKGKEECKESCHKKPCGCKAGEDCKCTDGKCGC |
| | | SannMT3La | IBPH01038578 | MEKCGANCKCTKCRCCGAAKSAAPSCCTAASARKADCGCDDCTCCQKSPCCASQKTCGKGCDCGDCK CCSSSACQKGKSCCGSGCKCDDCSCCSAAKGSGDCHCGDCKCCSSASC |
| | | SannMT3Lb | IAOG01004562 | METCGANCRCTKCRCCDAGKAAAPSCCTAAASKKADCSCDDCTCCQKSSCGSQKSCGKGCDCSDCKC CSSSACRTAGKTCCGSNCKCDVCTCCPAAKGTTCSH |

| | | | |
|---|---|---|---|
| | SannMT3S | IBPH01041663 | MPCGTCGPNCTCEKCRCCGAAAAPKCSANCNCGDCKCCGRTSANCRCGDCKCCGAAKC |
| Synema chikunii | SchiMT3L | ICKA01016492 | MDKCGPNCKCTKCRCCDAAKSATPACCQAKKTSCGCDDCQCCKSSGCGTQKSDCGKGCQCNCKCCSGTSQKDSCGSNCKCDVCTCCSAAKDSGSCHCGDCKCCSAASC |
| Tengella perfuga | TperMT2 | GGOF01120677 | MADVEKKTCCAEPKECCKDESCCQDGKGTCHTEKEGCADSCQKKPCGCKAGEDCKCKDGKCAC |
| | TperMT3L | GGOF01029183 | MDKCGTNCKCTKCRCCDAGKSAGPACCQAAAKKAACKCDDCRCCSSAACGKQKTCGKGCQCSDCKCCSSSSCQKDCCGSNCQCDDCRCCSAAKGSGDCKCGDCKCCSAATC |
| | TperMT3Sa | GGOF01037635 | MPGGPCGANCNCEKCRCCSAASKCTANCKCDTCRCCQRNQQCNCGDCKCCSSAKC |
| | TperMT3Sb | GGOF01118223 | MPGGPCGANCNCERCRCCSAASKCSSGCKCDTCRCCQRNQKCQCGDCKCCGAAKC |
| Tetragnatha brevignatha | TbreMT2 | GITM01027747 | MADAEKSCCAEPKECCKDESCCEEGKGPCHTEGKGCEDTCHQKPCGCKSGEECKCTSGSKCAC |
| Tetragnatha waikamoi | TwaiMT3S | GITR01027595 | MAGPCGANCTCEKCRCCSAASKCTSGCKCESCSCCQRSQKCNCGDCKCCGGAQC |
| Thanatus bungei | TbunMT3L | IALY01018183 | MDSCGPNCKCSKCRCCDAAKQSAPSCCCGASGKKSCDCTDCTCCKSSSCCEKEKSCGEGKCDNCKCCAASPCQTSNSSCGSSCKCDVCTCCSAAKGGADCHCGNCKCCSAASC |
| Tharpyna indica | TindMT3L | IBFA01024184 | MDKCGPNCKCTKCRCCDAAKSATPACCQVKKTKCGCDDCQCCKTSGASDCGKGCQCNCKCCSGATQKDSCGSNCKCDVCTCCSAAKDSGACHCGDCRCCSAASC |
| Thomisus labefactus | TlabMT2 | IAWV01026404 | MADVEKTSCCTESKECCKDESCCENGQGECHTGKEECKDTCHKKACGCKADEECKCAEGKCAC |
| | TlabMT3L | IAWV01002859 | MDKCGPNCKCTKCRCCDAAKSATPACCQAAAQKTDCGCDDCQCCKSSSCGTQKADCGKGCQCSCKCCSAQKDSCGSNCKCDVCTCCSAAKDSGACHCGDCRCCSAASC |
| | TlabMT3S | ICCK01008992 | MPGGPCGSNCSCEKCRCCSAAASASASKCGPNCRCDVCRCCNRGQNCRCGDCKCCGASKC |
| Thomisus okinawensis | TokiMT2 | ICDD01000078 | MADVEKTSCCTEPKECCKDESCCENGKGECHTGKEECKDTCHKKACGCKADEECKCSEGKCAC |
| | TokiMT3L | ICDD01018390 | MDKCGPNCKCTKCRCCDAAKSATPACCQAAAQKTDCGCEDCQCCKSCGTQKADCGEGCPCSCKCCSTQKDSCGSNCKCDVCTCCSAAKDSGACHCGDCRCCSAASC |
| | TokiMT3S | ICDD01032490 | MPGGPCGANCSCEKCRCCSAAASASASKCGPNCRCDVCRCCNRGQNCRCGDCKCCGASKC |
| Tibellus fengi | TfenMT2 | IAVL01019861 | MADVEKTTGCTEPKECCKDEACCENGQGACHDGKEECKDSCHKKACGCKAGEDCKCSSDGKCAC |
| | TfenMT3L | IAVL01011395 | MDRCGPNCKCSKCRCCDAATQSAPSCCCGASGKKSCDCTDCACCKSSSCCEKQKSCGEGKCDNCKCCSASACQTSKSSCGPSCKCDVCTCCSAAKGEADCHCGNCKCCSAASC |
| | TfenMT3S | IAVL01031865 | MPGGPCGANCTCEKCRCCSAASKCGANCKCDNCRCCSRGQNCRCGDCKCCGAAKC |
| Tmarus angulatus | TangMT2 | IBHB01033618 | MADVEKTSCCTEPKECCKDESCCQNGQGECHTGKAECKDTCQKKACGCKADEECKCSEGKCAC |
| | TangMT3L | IBHB01010593 | MDKCGPNCKCTKCRCCDAAKSPGCCQAAKTKCGCDDCQCCKSGAQKSDCSKGCQCNCKCCSKQKDSCGSNCKCDVCTCCSAAKDSGACHCGDCRCCSAASC |

| | | | |
|---|---|---|---|
| | TangMT3S | IBHB01011973 | MPGGPCGSNCTCEKCRCCTAASKCPASCKCDTCRCCQRSQKCHCGDCKCCGAAKC |
| Tmarus rimosus | TrimMT2 | IAWZ01003745 | MADVEKTSCCTEPKECCKDESCCQNGQGECHSGKEECKDTCHKKACGCKADEECKCSEGKCAC |
| | TrimMT3La | IAVJ01015074 | MDKCGPNCKCTKCRCCDAAKSPACCQAAKKKCGCDDCQCCKSGAQKVDCGKGCECNCKCCSTQKDS CGSNCKCDVCTCCSAAKDSGACHCGDCRCCSAASC |
| | TrimMT3Lb | IAWZ01003419 | MEKNCGPNCKCTKCRCCDAAKGKDCGCENCRCCDASSCKSGKSCGSGCDCRDCKCCSASSCCQTSSC CGSGCKCDKCSCCAAAKGKGTCNCGDCKCCSAAKC |
| | TrimMT3S | IAWZ01037322 | MPGGPCGSNCSCEKCRCCSAAASASASRCGPNCKCDVCRCCNRGQNCRCGDCKCCGASKC |
| Trochosa aquatica | TaquMT2 | IBZV01030133 | MADVEKTSCCTESKECCKDESCCENGQGACHTGKEECKDSCHKKACGCKAGEDCKCSDGKCGC |
| | TaquMT3L | IASZ01035477 | MDKCGANCKCTKCRCCDAAKDTTPACCQAAAKKTTCGCDDCSCCSSSGCCEQKTSGKGCQCSCKCCS SQKNTCGSSCKCDVCSCCSAAKGSGDCHCGNCKCCSTACC |
| | TaquMT3Sa | IASZ01021367 | MPCGTCGPNCTCEKCRCCGAAAAPKCSANCNCGDCKCCGRTSANCRCGDCKCCGAAKC |
| | TaquMT3Sb | IBZV01027233 | MPGGPCGPNCSCEKCRCCSAAASSGGRCGPNCRCDVCRCCNRGQNCRCGDCKCCGASKC |
| Trochosa terricola | TterMT2 | IBIR01014579 | MADVEKTSCCTEPKECCKDESCCENGQGTCHTGKEECKDSCHKKACGCKAGEDCKCSDGKCAC |
| | TterMT3La | IBIR01022592 | MDKCGPNCKCTKCRCCDSAKQASPCCTGGKKADCGCDDCRCCSSSPCATQKSCGKGCQCGDCKCCS SSPCQSAKSSCGSNCKCDVCTCCSAAKGGGDCHCGDCKCCGAAVC |
| | TterMT3Lb | IBIR01015641 | MDKCGANCKCTKCRCCDAAKESTPACCQAAAKKTTCGCDDCSCCSSSGCCEQKASGKGCQCSCKCCS PASSQKNTCGSSCKCDVCSCCSAAKGSGDCHCGNCKCCSTACC |
| | TterMT3S | IBIR01028004 | MPGGPCGSNCTCEKCRCCSAASKCSDGCKCDNCKCCQRGQAKCHCGDCKCCGAAKC |
| Uloborus diversus | UdivMT2 | GJZJ01180553 | MADVEKKACCAEPKECCKDESCCQDGKGACHTEKEGCADSCQKKPCGCKAGEDCKCKDGKCAC |
| | UdivMT3S | GJZJ01032433 | MPGGPCGANCTCEKCRCCGAAAKCSASCKCETCRCCQRGQNCRCGDCKCCGAAKC |
| Vulsor sp. | VspMT2 | ICOO01003872 | MADVEKTSCCTEAKECCKDESCCENGQGACHTGKEECKDTCHKKACGCKAGEDCKCSDGKCAC |
| | VspMT3L | ICOO01039941 | MERKCGPNCKCTKCRCCDSGKQAAPPCCTAAAAKKAACGCDDCTCCKSSSCGAQKSCGKGCQCGDCK CCSSSACQTSKGSCGSSCQCDVCTCCSAAKGGGDCHCGDCKCCSSAVC |
| Xysticus insulicola | XinsMT3L | IAFS01028859 | MDKCGPNCKCTKCRCCDAAKSVTPACCQASAKKTNCGCDDCQCCKSSSCGTQKSDCGKGCQCNCKC CSGGTQKDSCGSNCKCDVCTCCSAAKDSGACHCGDCRCCSAASC |
| Xysticus transversomaculatus | XtraMT3S | IAPE01005889 | MPCGTCGPNCTCEKCRCCGAAAAPKCSANCNCGDCKCCGRTSANCRCGDCKCCGAAKC |
| Zelotes asiaticus | ZasiMT2 | IAGO01029205 | MADVEKTSCCTEQKECCKDESCCENGQGACHTGKEECKDSCHKKACGCKGGEDCKCTDGKCGC |
| | ZasiMT3L | IAGO01012009 | MEKCGPNCTCANCRCCEAAGAAPSCCTAASKCSCADCSCCRSAPCCASKKSCGKGCQCGDCKCCSSS ACQTGKSCCGSDCNCTVCRCCSAAKSGGDCRCGDCKCCSSASC |

| | | | | |
|---|---|---|---|---|
| Mesothelae suborder | | | | |
| | Heptathela helios | HhelMT2 | IAMB01023112 | MADTEKTACCTAEKECCKDSSCCEEGKGKECCDNNPDCCKDGAAKKPCGCKEGEECKCKEGKCAC |
| | | HhelMT3L | IAMB01018332 | MADKCGPNCKCQKCRCCDASVKAQAAGCCGGKSCTCEKCQCCSSAGCGAAAKSCGSNCTCDDCRCC SEAQCAQKCGSGCTCDVCRCCSASRGQGRCQCGDCRCCSSASC |
| | | HhelMT3Sa | IAMB01031069 | MPGPCGQNCTCERCRCCAAATKCSSGCRCDTCRCCTRGQKCTCGDCKCCGAAKC |
| | | HhelMT3Sb | IAMB01033754 | MPGPCGQNCTCERCRCCSAATKCSSGCRCDTCRCCTRGQKCNCGDCKCCGAAKC |
| | Liphistius murphyorum | LmurMT2 | ICAH01033317 | MADAEKTACCTADKECCKDSSCCEEGKGKECCDNNPDCCKEGAAKKPCGCKEGEECKCQEGKCAC |
| | | LmurMT3L | ICAH01020478 | MAGKCGPNCKCEKCRCCDASGPASASAKAQGAGCCGGSSCSCAQCQCCTSAGCGAATASCGPSCTCD ACKCCSEAQCGQKCGSGCTCDVCRCCSASRGQGRCQCGDCRCCSSASC |
| | | LmurMT3S | ICAH01016204 | MPGPCGQNCTCERCRCCSAATKCSAGCRCDTCRCCTRGQKCNCGDCKCCGAAKC |
| | Ryuthela nishihirai | RnisMT2a | IAQS01038821 | MADTEKTACCTAEKECCKDSSCCEEGKGKECCDNNPDCCKDGAAKKPCGCKEGEECKCQEGKCAC |
| | | RnisMT2b | IAIP01007631 | MADTETTACCTAEKQCCKDSSCCEQGKGKECCDNHPDCCKDGAAQKPSGCKEGEECKSQEGKCAC |
| | | RnisMT3L | IAIP01006156 | MAGKCGPNCKCEKCRCCDASGKAQAAGCCRGESCTCEKCQCCSSAGCGAATKNCGPNCTCDDCRCC SEAQCAQKCGAGCTCDVCRCCSASKGQGRCNCGDCRCCSSASC |
| | | RnisMT3Sa | IAIP01026854 | MPGPCGPNCTCDKCRCCTAATKCSEGCKCDTCRCCERGQKCQCGDCKCCGAAKC |
| | | RnisMT3Sb | IAQS01029875 | MPGPCGQNCTCDRCRCCSAATKCSSGCRCDTCRCCTRSQKCNCGDCKCCGAAKC |
| | | RnisMT3Sc | IAQS01006286 | MPGPCGQNCTCERCRCCSAATKCSAGCRCDTCRCCTRGQKCNCGNCKCCGAAKC |
| Mygalomorphae suborder | | | | |
| | Antrodiaetus roretzi | ArorMT2 | IBPI01014909 | MADVEKKSCCSPEKECCKENECTAEGTECCHKDSDGCKDGAVQKKPCGCKVGEDCKCKEGKCAC |
| | | ArorMT3L | IBPI01012923 | MESKCGPNCECTECGCCAALSSATKDCCGSQCTCEAGKCECCDTTKCCGEGVQCSCEDCQCCSSAGC GKKQSGPCCACDDCNCCSSAACISSSRCGSGCQCDDCSCCPAAHGQGTCECGDCKCCSTASC |
| | | ArorMT3Sa | IBPI01004899 | MPGPCGQNCTFDRCRCLAAATGCSGGCRCENCRCCTRGQKCNCVDCKCCGAAKC |
| | | ArorMT3Sb | IBPI01005671 | MPGPCGQNCTCERCRCCTAATRCSSGCRCETCRCCTRGQKCNCGDCKCCGAAKC |
| | Calommata signata | CsigMT2 | IBXS01010418 | MADVEKNTCCSPEKECCKENECTAEGKECCHGDTECCKDGAAQKKPCGCKQGEDCKCQDGKCAC |
| | | CsigMT3L | IBXS01029286 | MEGKCGSQCQCTECRCCGALSSASKGCCGDQCCCEAGKCQCCDTAQSCGGAEKCACSDCQCCSSSG CHKQEAAARCGPSCDCDDCNCCGTAACQTSSRCGSGCQCAGDCSCCPAAHGQGACTCGDCKCCSSA |

| | | | | |
|---|---|---|---|---|
| | | CsigMT3S | IBXS01007514 | MPGPSGPNCSCERCSCCAAATRCSSGCRCETCRCCSRGQRCNCGDCKCCGAAKC |
| | Conothele fragaria | CfraMT2 | IAVG01026659 | MADVEKTSCCTTEEDCCKGGDCTTEGKDCSKEHDCCKDGSATKPCGCKQGEDCKCKDGKCGC |
| | | CfraMT3Sa | IAVG01019714 | MPGPCGQNCNCQQCRCCTAATKCSDGCRCETCRCCTRSQRCNCGDCKCCGAAKC |
| | | CfraMT3Sb | IAVG01034978 | MPGPCGQNCNCERCRCCTAAAKCSEGCRCETCRCCTRSQKCNCGDCRCCGAAKC |
| | Macrothele gigas | MgigMT2 | IAEV01010058 | MADVEKTACCTTEEDCCKGSECTTEGKDCSKEHDCCKDGSAQKPCGCKKGEDCKCQEGKCAC |
| | | MgigMT3Sa | IAEV010076969 | MPGPCGQNCNCGQGCRCCTAAVRCTDGCRCETCRCCSRSQKCNCGDCRCCGAAKC |
| | | MgigMT3Sb | IAEV01004369 | MPGPCGQNCNCERCRCCTAASKCTADCRCETCRCCSRSQKCECGDCKCCGAAKC |
| | Rhianodes atratus | RatrMT2a | GIXJ01248059 | MADVEKTSCCTETKECCKDESCCENGQGACHTGKEECKDSCHKKPCGCKAGEDCKCSDGKCGC |
| | | RatrMT2b | GIXJ01248060 | MADVEKTSCCTEPKECCKDESCCENGQGSCHTGKEECKDTCHKKACGCKAGEDCKCSEGKCAC |
| | | RatrMT3Sa | GIXJ01115636 | MPGPCGQNCNCERCRCCTAATRCSDGCRCETCRCCTRSQKCNCGDCRCCGAAKC |
| | | RatrMT3Sb | GIXJ01115623 | MPGPCGPNCNCGDQCRCCTAATKCSAGCRCETCRCCTRSQRCNCGDCRCCGAAKC |
| Ixodida order | | | | |
| | Amblyomma americanum | AameMT1 | GAGD01006548 | MDRACKCGDPKCTCTTECKCGTGGCPCDKCKK |
| | Dermacentor variabilis | DvarMT1a | GGQS01008255 | MDKPACKCGDPKCVCTTQCKCATGGCPCDRCKK |
| | | DvarMT1b | GGQS01015897 | MERACKCGDPKCVCTTQCKCASGGCPCGKCKK |
| | Hyalomma dromedarii | HdroMT1 | GFGI01011981 | MDRSCKCGDPKCVCTTQCKCASGGCPCGKCKK |
| | Ixodes holocyclus | IholMT1a | GIBQ01032995 | MDRACKCGDPKCTCTTSCKCASGECPCDKCKPQSPKRACCK |
| | | IholMT1b | GIBQ01041329 | MDRACKCGDPKCICTTTCKCASGECPCDKCKHKSPKRACCK |
| | Ixodes persulcatus | IperMT1a | GBXQ01027128 | MDRACKCGDPKCTCTTSCKCPDGDCPCDKCKAKSPSK |
| | | IperMT1b | GBXQ01027126 | MDRACKCGDPKCTCTTSCKCPNGDCPCDKCKTKSPAK |
| | Ixodes ricinus | IricMT1a | GIDG01028662 | MDRACKCGDPKCTCTTSCKCADGDCPCDKCKAKSPSK |

| | | | | |
|---|---|---|---|---|
| | | IricMT1b | GIDG01001147 | MDRDCKCGDPKCICTDSCKCAAGECPCDKCKPKSAEKKCCC |
| | Ixodes scapularis | IscaMT1a | GGIX01090004 | MDPVCKCGDPKCTCTTSCKCPGGDCPCDKCKTKSPAK |
| | | IscaMT1b | GGIX01470521 | MDRVCKCGDPKCTCTTSCKCPDGDCPCDKCKAKSPSK |
| | Ornithodoros erraticus | OerrMT1 | GFWV01004713 | MDRACKCGDPKCTCTTECKCGTGGCPCTKCKK |
| | Rhipicephalus annulatus | RannMT1a | GBJT01020093 | MNKPPCKCGDPKCVCTTQCKCATGGCPCDRCKK |
| | | RannMT1b | GBJS01007861 | MDRACKCGDPKCICTTQCKCSTGGCPCDKCKK |
| | Rhipicephalus appendiculatus | RappMT1a | GEFI01043136 | MDKPPCKCGDPKCVCTTQCKCASGGCPCGRCKK |
| | | RappMT1b | GEFJ01061002 | MDRACKCGDPKCSCTTQCKCSTGGCPCDKCKK |
| | Rhipicephalus bursa | RburMT1 | GFZJ01015271 | MDRACKCGDPKCICTTQCKCSTGGCPCDKCKK |
| | Rhipicephalus haemaphysaloides | RhaeMT1 | GIJA01013269 | MDRACKCGDPKCVCTTQCKCSTGGCPCGKCKK |
| | Rhipicephalus microplus | RmicMT1a | GIZL01038324 | MDKPPCKCGDPKCVCTTQCKCATGGCPCDRCKK |
| | | RmicMT1b | GIZL01194700 | MDRACKCGDPKCICTTQCKCSTGGCPCDKCKK |
| | Rhipicephalus pulchellus | RpulMT1 | GACK01009326 | MDRACKCGDPRCVCTTQCKCSTGGCPCDKCKK |
| | Rhipicephalus sanguineus | RsanMT1 | GINV01000976 | MDKPACKCGDPKCVCTTQCKCATGGCPCDRCKK |
| | Rhipicephalus zambeziensis | RzamMT1a | GFPF01021474 | MDKPPCKCGDPKCVCTTQCKCASGGCPCGRCKK |
| | | RzamMT1b | GFPF01017025 | MDRACKCGDPKCTCTTQCKCSTGGCPCDKCKK |
| Opiliones order | | | | |
| | Aoraki longitarsa | AlonMT1 | SRA | MPDPACSCDDECTCGTDCKCGAECKCEKCKGSAKCNCKNGGECTCTTCTCK |
| | Avima matintaperera | AmatMT1 | SRA | MSDPCNCGDECTCGTECTCGEECKCKTCKKPDSQNEATETS |
| | Brasilogovea microphaga | BmicMT1 | SRA | MPGPGCNCGDECKCGSDCKCGSDCKCDKCKGTDKCQCTGNADCQCXKCTCK |
| | Egaenus convexus | EconMT1 | GERV01018384 | MPDECKCKKDGKCTCQGECKCGEGCACDCCKTTAECKCEKCTCDPCSCK |

| | | | | |
|---|---|---|---|---|
| | Hesperonemastoma modestum | HmodMT1 | SRA | MSDPCSCLEGECTCGTDCKCGADCACEKCKSAAQATENLTCAKCSAEPCSC |
| | Karos barbarikos | KbarMT1 | SRA | MSDPCKCDDECKCGSECTCGDECKCEHCKKGKTDSTSEAD |
| | Larifuga capensis | LcapMT1 | SRA | MDPCKCDEECTCGSDCKCGADCKCETCKKGDDQGETATSEDS |
| | Neopurcellia salmoni | NsalMT1 | SRA | MPDPACSCGDECTCGSDCKCGADCKCETCKGSTPKCNCTSGGECTCTNCDCK |
| | Pachylicus acutus | PacuMT1 | SRA | MSDPCKCDEECKCGSECTCGEECKCEKCKKGSETMNKLH |
| | Phalangium opilio | PopiMT1a | SRA | MPGPGCSCGEECKCGSDCKCGSDCKCDKCKGSAKCQCTGAADCTCAKCSCK |
| | | PopiMT1b | GDAO01000195 | MPDSCKCKQDGKCTCEGECKCGESCKCDCCKTTAECKCEKCTCDPCSCK |
| | | PopiMT1c | SRA | MPDSCKCKEAGKCTCQPECKCGPECKCDCCKTTAACKCETCTCDPCTCK |
| | Rakaia magna australis | RmagMT1 | SRA | MPDPACNCGDECSCGQDCKCGADCKCEKCKKGSAAKCDCTNGGECTCTNCTCK |
| | Synthetonychia proxima | SproMT1 | SRA | MADPCACEDDCQCGSDCKCGADCKCEKCKKGQADQSEDQASETSQ |
| | Vonones ornata | VornMT1 | SRA | MSDPCKCDEECKCGGECKCGEECKCEKCKKGQADSQSDSAPSETS |
| Scorpiones order | | | | |
| | Centruroides noxius | CnoxMT1 | SRA | MPDPACKCGDSKCNCQTKCTCSSGGCKCDKCHCKCKEGGSCSCSSCSCK |
| | Chiromachus ochropus | CochMT1 | SRA | MPDPACNCGDSGCTCTTSCQCASKGCKCSKCTCQCKEGGSCSCSPCTCKIGRAHV |
| | Hadogenes paucidens | HpauMT1 | SRA | MPDPACKCGDSNCTCTSSCECASGGCKCSKCACKCKEGGSCACNPCNCK |
| | Hormiops davidovi | HdavMT1 | SRA | MPDPACKCGDSNCSCSSSCGCASGECKCSKCACKCKEGGSCSCSPCNCK |
| | Opisthacanthus asper | OaspMT1 | SRA | MPDPACGCSDSACTCTSSCKCASNGCKCSKCTCQCKEGGSCSCNPCTCK |
| | Palaeocheloctonus pauliani | PpauMT1 | SRA | MPDPACNCGDSSCTCTTSCQCASGGCKCSKCTCKCKEGGSCSCSPCTCK |
| | Tityus serrulatus | TserMT1 | GBZU01013840 | MPDPACKCGDSKCTCQTKCTCSSGGCKCDKCHCKCKEGGSCSCTSCNCK |
| Solifugae order | | | | |
| | Eremobates sp. | EspMT1 | SRA | MADPACQCGEDSCTCGADCKCSAGDCKCTKCQPKQQCKCTEACSCNPSS |

| | | | |
|---|---|---|---|
| Galeodes sp. | GspMT1 | SRA | MPDPTCKCQEDSCTCGSDCKCTASECKCTKCQPKQSCKCSGSCSCPPASS |
| Gluvia dorsalis | GdorMT1 | GDAP01009971 | MADPDCQCGEDTCTCGSDCKCSSGGCKCTKCRPKARCTDDHAEETSPSSS |
| Solpugema sp. | SspMT1 | SRA | MADPACQCGEDSCTCGPDCKCTSGECKCTKCQPKQACQCTESHGEESCPQSS |

*MT sequences were retrieved from NCBI databases (their accession numbers are provided) or reconstructed from raw sequence data obtained from the Sequence Read Archive (SRA).
