## Supplementary material for "The Evolutionary History of Metallothioneins in Horseshoe Crabs, Scorpions, Spiders and other Chelicerate Species": Legends of Supplemental Figures

### Figure legends

**Figure S1. Amino acid alignment of type 1 metallothioneins in chelicerates.** The alignment includes MT1 sequences from the classes Pycnogonida, Merostomata, and Arachnida, listed alphabetically. MT names correspond to those included in Table S1. Sequences were retrieved from NCBI database (accession numbers provided next to each MT name) or reconstructed form Sequence Read Archives (SRA). Conserved cysteine residues are highlighted in yellow. Below each alignment, the Cys motifs in the D1a (dark blue box) and D1b (light blue box) domains are shown. (A) Alignment of Pycnogonida MT1s showing a bidomain D1a/D1b organization. (B) Alignment of Merostomata MT1s displaying either a D1a/truncated-D1a or a D1a/D1a/t-D1a domain arrangement. (C) Alignment of Arachnida MT1s revealing a D1a/t-D1a structure or a monodomain D1a organization in ticks and some Opiliones and Solifugae species.

**Figure S2. Amino acid alignment of type 2 metallothioneins in spiders.** The alignment includes MT2 sequences from the Araneae order (Arachnida class), across its three suborders (Araneomorphae, Mesothelae and Mygalomorphae), and listed alphabetically within each group. MT names correspond to those included in Table S1. Sequences were retrieved from NCBI database (accession numbers provided next to each MT name) or reconstructed form Sequence Read Archives (SRA). Conserved cysteine residues are highlighted in yellow. Below the alignment, the Cys motifs and a highly conserved His in the D2a (red box) and D2b (pink box) domains are shown. Spider MT2 shows a bidomain D2a/D2b organization.

**Figure S3. Amino acid alignment of type 3 metallothioneins in spiders.** The alignment includes MT3 sequences from the Araneae order (Arachnida class), across its three suborders (Araneomorphae, Mesothelae and Mygalomorphae), and listed alphabetically within each group. MT names correspond to those included in Table S1. Sequences were retrieved from NCBI database (accession numbers provided next to each MT name) or reconstructed form Sequence Read Archives (SRA). Conserved cysteine residues are highlighted in yellow. Below each alignment, the Cys motifs in the D3 repeats (green box) are shown. (A) Alignment of MT3S forms, consisting of three imperfect D3 repeats. (B) Alignment of MT3L forms consisting of five imperfect D3 repeats (six, in Mygalomorphae MT3L sequences).

**Figure S4.** Deconvoluted ESI-MS spectra of MT proteins, recorded at pH 2.4, recombinantly produced in Zn-, Cd- and Cu-enriched media. (A) NgraMT1, LpolMT1a, LpolMT1b, IricMT1a, and HpauMT1 proteins. (B) LcoeMT2, LcoeMT3Sa, and LcoeMT3L proteins. Glycosylated species containing one (*), two (**) or even three (***) hexamer units have been detected.

**Figure S5. Amino acid alignment of metallothioneins from chelicerates, myriapods, mollusks and chordates.** The alignment includes the seven MT1s from Pycnogonida species with a D1a/D1b organization, three putative MT sequences from two myriapod species, *Lithobius forficatus* and *Glomeridella minima*, which exhibit an identical domain organization, ten mollusk MTs composed of $\beta$2 and $\beta$1 domains {Calatayud, 2021 #10915}, and seven chordate MTs (four human MTs, representing the vertebrate forms, and three MT sequences from different cephalochordate species) consisting of $\beta$ and $\alpha$ domains with 9-11 conserved cysteine residues {Calatayud, 2021 #10913}.
